# Conformational changes in Lassa virus L protein associated with promoter binding and RNA synthesis activity

**DOI:** 10.1101/2021.06.24.449696

**Authors:** Tomas Kouba, Dominik Vogel, Sigurdur R. Thorkelsson, Emmanuelle R. J. Quemin, Harry M. Williams, Morlin Milewski, Carola Busch, Stephan Günther, Kay Grünewald, Maria Rosenthal, Stephen Cusack

## Abstract

Lassa virus, which causes annual outbreaks in West Africa with increasing case numbers in recent years, is recognized by the WHO R&D blueprint as a significant threat for public health with high epidemic potential and no effective countermeasures. The viral large (L) protein, which contains the RNA-dependent RNA polymerase, is a key player for transcription of viral mRNA and genome replication. Here we present nine cryo-EM structures of Lassa virus L protein in the apo-, promoter-bound pre-initiation and active RNA synthesis states. We characterize distinct binding pockets for the conserved genomic 3’ and 5’ promoter RNAs and show how full-promoter binding induces a distinct pre-initiation conformation. In the apo- and elongation states, the endonuclease is inhibited by the binding of two distinct L protein peptides in the active site, respectively, whereas in the pre-initiation state, the endonuclease is uninhibited. In the stalled, early elongation state, a template-product duplex is bound in the active site cavity together with an incoming non-hydrolysable nucleotide. In this configuration, the full C-terminal region of the L protein, including the putative cap-binding domain, is highly ordered. The structural data are complemented by in vitro and cell-based studies testing a broad range of L protein mutants to probe functional relevance. These data advance our mechanistic understanding of how this flexible and multifunctional molecular machine is activated and will underpin antiviral drug development targeting the arenavirus L protein.

## Introduction

Lassa virus (LASV) is a segmented, negative strand RNA virus belonging to the family of *Arenaviridae* within the *Bunyavirales* order. It is a rodent-borne virus, endemic to West Africa and the causative agent of Lassa haemorrhagic fever, a febrile illness with increasing case numbers and a case fatality rate among hospitalized patients of ~15% in Nigeria in 2018 ^1^. Recent studies applying computational modelling predict a total number of ~900,000 human infections per year across West Africa ^2^. The large (L) protein of LASV is a multi-domain molecular machine that binds the conserved 3’ and 5’ ends (the ‘promoter’) of each of the two viral RNA (vRNA) genome segments (denoted L and S) and plays a central role in the viral life cycle, which is entirely cytoplasmic. The L protein contains RNA-dependent RNA polymerase (RdRp) activity and catalyses both viral transcription and genome replication. Each vRNA segment is template for the synthesis of two different types of RNA products: capped viral transcripts (of L and NP genes) as well as an unmodified full-length complementary RNA (cRNA) copy, which is an intermediate of viral genome replication. The cRNA is a template for the second stage of replication, the synthesis of vRNA genome copies, as well as the production of further mRNA (of GPC and Z genes) by transcription. Transcription is initiated using a capped primer derived from host mRNA by a yet to be elucidated ‘cap-snatching’ mechanism involving the intrinsic endonuclease (EN) of the L protein and possibly its cap-binding domain (CBD) ^3^. This results in viral mRNAs that have 1-7 host-derived nucleotides at the 5’ end ^4–6^. Viral genome replication is initiated by a prime-and-realign mechanism resulting in an extra G nucleotide at the 5’ end of the vRNA and cRNA ^7^. The first structural studies of the complete arenavirus L protein were conducted on Machupo virus (MACV), which is related to LASV but belongs to the group of New World arenaviruses found in South America. Negative stain electron microscopy studies at low resolution revealed a donut-like molecule with accessory appendages ^8^. In 2020, the first models of MACV and LASV L proteins were proposed based on cryo-electron microscopy (cryo-EM) with overall resolutions of ~3.6 Å and 3.9 Å, respectively ^9^. These structures revealed that arenavirus L proteins are structurally similar to the polymerases of the related La Crosse and influenza viruses ^10–14^. However, the reported arenavirus structures are incomplete and do not show the L protein in an active conformation. Here we present nine cryo-EM structures that provide insights into the conformational rearrangements of LASV L protein that occur upon its activation into a functional RNA synthesis elongation state. This comprehensive structural study is complemented by biochemical data from *in vitro* assays with purified L protein and selected mutants as well as functional data in cells using the LASV mini-replicon system ^15^. The results presented enhance our mechanistic understanding of the multifunctional LASV L protein and will guide targeted drug development approaches in the future.

## Results

### Overview of structures obtained

Nine cryo-EM structures of LASV L protein have been determined in the apo-state, bound to the 3’ end of the genomic vRNA alone, bound to the full promoter, comprising the highly complementary 3’ and 5’ ends of the vRNA, or in a stalled, early elongation state (Table 1).

**Table 1.**
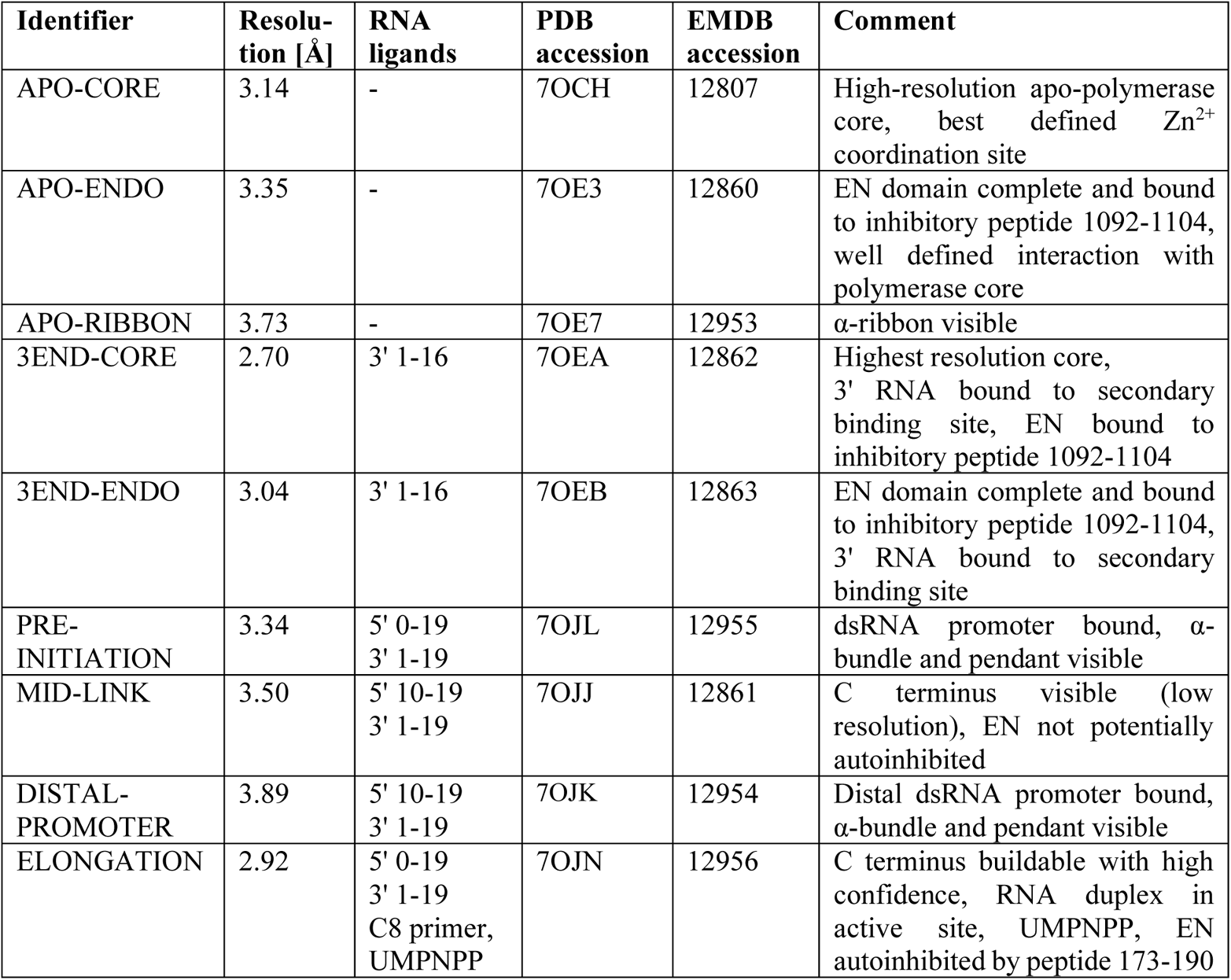
Overview about the different LASV L protein structures

From grids of the apo-state, two different 3D classes were separated. In the **APO-ENDO** structure, at 3.35 Å resolution (Fig. 1a), the N-terminal EN is clearly resolved, packing against the polymerase core and with a peptide (residues 1092-1104) from the central region of the L protein bound in its active site cavity, presumably inhibiting its activity. In the second class, denoted **APO-RIBBON** (Fig. 1a), at 3.73 Å resolution, the EN is not resolved, but residues 822-1110 of the L protein form an extended structure including an α-bundle (α-ribbon, 843-884, with a third helix 907-925 packing against it) that is not visible in the **APO-ENDO** structure. Masked refinement of the common regions of the two apo-structures yielded a map of the **APO-CORE** (Fig. 1a) with an improved resolution of 3.14 Å that allowed a more accurate model to be built.

**Figure 1.**
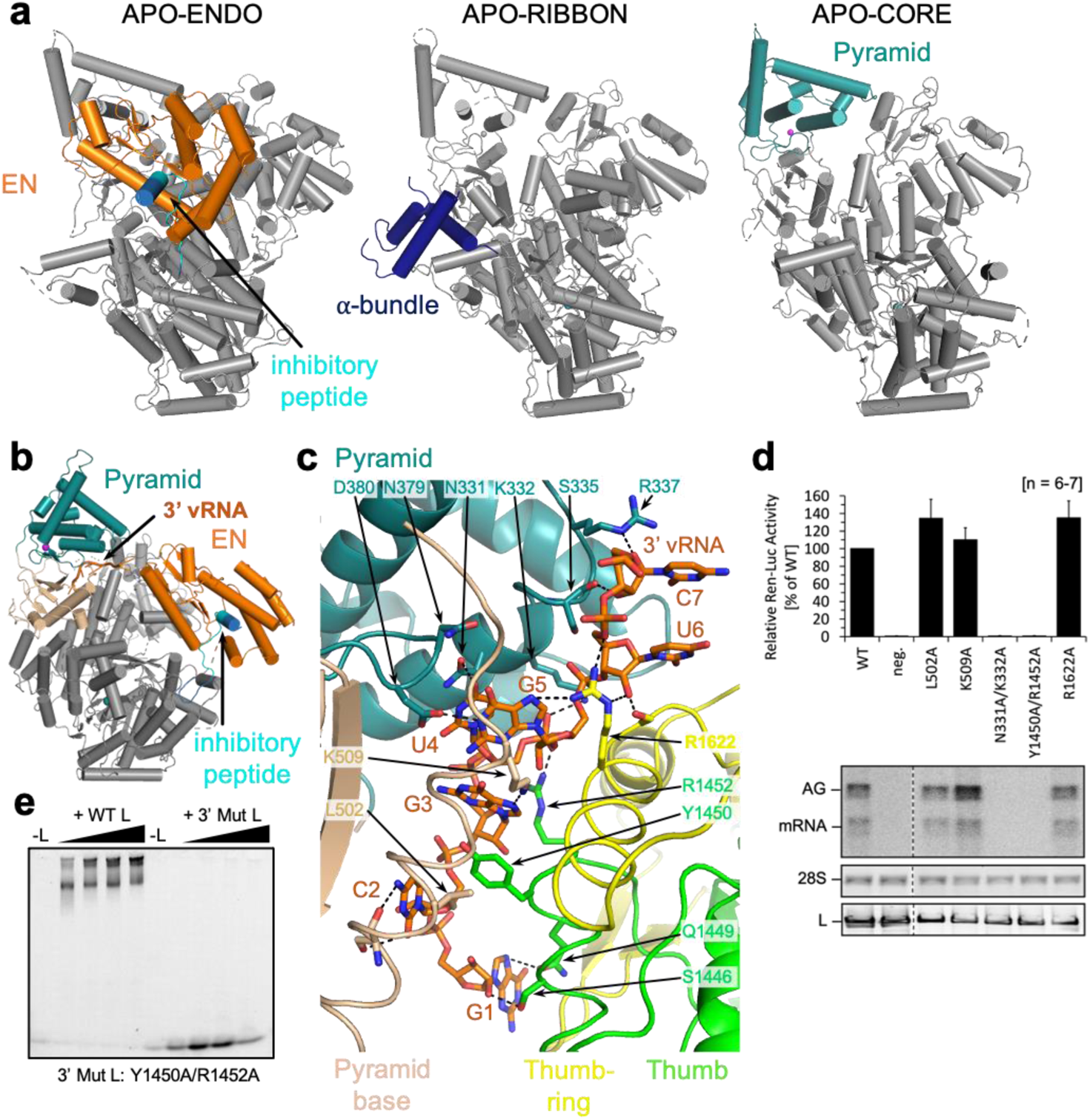
L protein in the apo-state and with 3’ viral RNA bound in the secondary binding site. **a.** Ribbon diagram presentations of the structures APO-ENDO, APO-RIBBON and APO-CORE. Each of the respective experimental maps resolves distinct regions of the L protein better than the others and those regions are shown in color and indicated by name in the respective structures. **b.** Overall structure of L protein 3END-CORE as a ribbon diagram with the 3’ vRNA bound below the pyramid domain. Pyramid (teal), pyramid base (wheat), EN domain (orange) and the inhibitory peptide (cyan) are highlighted. **c.** Close-up of the secondary 3’ vRNA binding site with the 3’ vRNA nucleotides 1-7 (orange), pyramid domain (teal), pyramid base (wheat) as well as thumb (green) and thumb-ring (yellow) domains. Important amino acids in the RNA:protein interface are shown as with respective labels. Hydrogen bonds are indicated by dotted lines. For selected regions secondary structure depiction was disabled to enhance visibility. **d.** LASV mini-replicon data for L proteins with mutations in the secondary 3’ RNA binding site presenting luciferase reporter activity (in standardized relative light units relative to the wild-type L protein (WT), mean average of 6-7 biological replicates) (top), Northern blotting results with signals for antigenomic viral RNA (AG), viral mRNA (mRNA) and 28S ribosomal RNA (28S) as a loading control (middle), and Western blot detection of FLAG-tagged L proteins (L) to demonstrate general expressibility of the mutants (bottom). **e.** Electrophoretic mobility shift assay of wild-type L protein (WT L) and mutant Y1450A/R1452A (Mut L) with 10 nt 3’ viral RNA. L protein concentrations ranging from 0-1 µM and 0.2 µM of fluorescently labelled 3’ vRNA (Supplementary Table 2) were used (see methods). (High-resolution figure available at the end of this PDF)

Upon incubation of L protein with nucleotides (nts) 1-16 of the vRNA 3’ end alone (structures denoted **3END-CORE**, 2.70 Å, **3END-ENDO,** 3.04 Å) (Fig. 1b), nucleotides 1-6 from the 3’ end bind specifically in a buried groove under the pyramid, a prominent feature in the N-terminal region of the L protein (Fig. 1c). This site corresponds to the secondary 3’ end-binding site previously described for influenza virus, La Crosse virus (LACV) and MACV polymerase proteins ^9,13, 16–18^. In these structures, the EN remains in the inhibited conformation as observed in the **APO-ENDO** structure (compare Figs. 1a and b). When the full vRNA promoter is bound (5’ end nts 0-19, including an additional G0 according to the product expected from the prime-and-realign initiation mechanism, 3’ end nts 1-19), a pre-initiation complex (**PRE-INITIATION**) is observed at 3.34 Å resolution (Fig. 2a, Supplementary Fig. 1). This structure reveals that the LASV vRNA promoter is organised similarly to those of influenza virus ^10^ and LACV ^13,14^ in comprising a single-stranded 5’ end folded as a hook, a distal duplex region and a single stranded 3’ end, only partially visible, directed towards the RNA synthesis active site (Fig. 2c). Overall, the protein conformation of the **PRE-INITIATION** structure resembles that of the **APO-RIBBON**, with an additional partially ordered insertion domain, previously denoted the pendant ^9^, packing against the 3’ strand of the promoter (see below) (Fig. 2a). Two further structures were obtained from a sample in which the L protein was incubated with a truncated promoter (5’ nts 10-19, 3’ nts 1-19), which lacks nucleotides 0-9 of 5’ end. One 3D class from this sample, obtained by focussed refinement on the promoter-bound region, (**DISTAL-PROMOTER,** 3.89 Å resolution), closely resembles the full promoter (**PRE-INITIATION**) structure, but additionally reveals a new position of the EN, without inhibitory peptide bound (Supplementary Fig. 1). The second 3D class from the same grid (**MID-LINK,** 3.50 Å resolution) (Supplementary Fig. 2), obtained by focussed refinement of the other end of the polymerase, shows density for the C-terminal region of the L protein beyond residue 1834. This allows tentative modelling of domains that resemble the mid-link and 627-domains of the influenza virus polymerase PB2 subunit. At very low resolution, an envelope of the putative cap-binding (CBD-like) domain is observed. A final structure (**ELONGATION**, 2.92 Å resolution) captures an early elongation state initiated with an uncapped primer and stalled after incorporation of four nucleotides by an incoming non-hydrolysable UTP analogue, UMPNPP (Figs. 3 and 4). In this structure, the promoter duplex is disrupted due to translocation of the template and a duplex of eight base pairs occupies the active site cavity (Fig. 4a). The complete C-terminal domain is well resolved, and, due to rotation of the CBD-like domain, is now in a bent configuration, rather than the extended configuration seen in the **MID-LINK** structure. The C-terminal domain, together with the EN, now in a third distinct position, forms a ring around the putative product exit channel (Fig. 4b).

**Figure 2.**
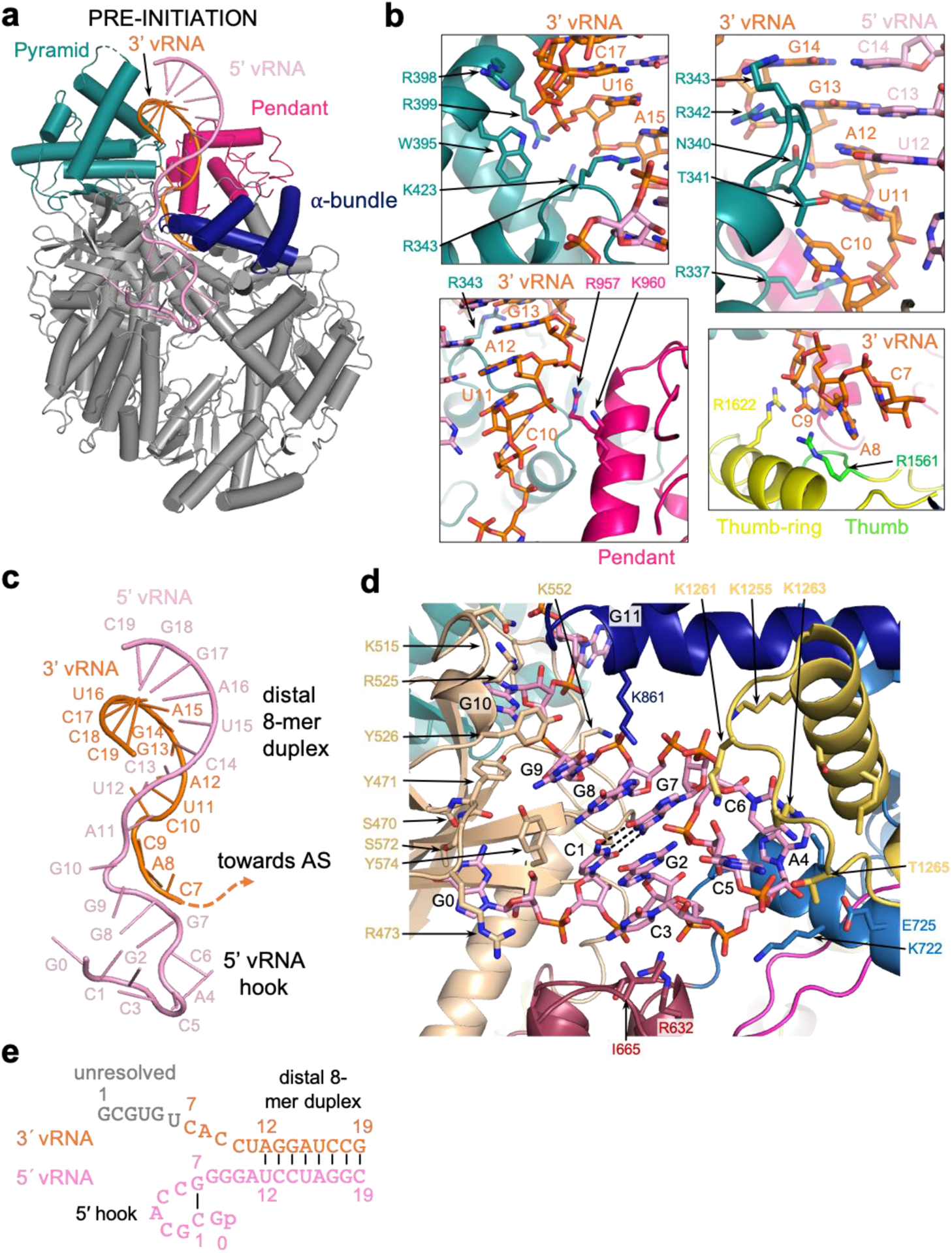
L protein in the pre-initiation state. **a.** Ribbon diagram of the PRE-INITIATION structure with pendant domain (pink), α-bundle (dark blue), 3’ vRNA nts 7-19 (orange) and 5’ vRNA nts 0-19 (pink) highlighted in color and indicated by name. **b.** Interactions of the L protein pyramid (teal), pendant (pink) thumb-ring (yellow) and thumb (green) domains with the 3’ vRNA are shown. Important amino acid side chains and the RNA nucleotides of 3’ and 5’ vRNA are shown as sticks with respective labels. **c.** Viral RNA observed in this structure with a 5’ vRNA hook structure composed of 5’ vRNA nts 0-9 and a distal duplex region involving 5’ vRNA nts 12-19 and 3’ vRNA nts 12-19. The 3’ vRNA nts 1-11 are directed towards the RdRp active site (towards AS) but not resolved. **d.** Close-up of the 5’ RNA hook binding site involving the fingers domain (blue), fingernode (light yellow), pyramid base (wheat) and helical region (raspberry). Residues important for the RNA:protein interface and nucleotides are shown as sticks and are labelled. **e.** Schematic presentation of the promoter RNA (3’ vRNA in orange, 5’ vRNA in pink) in the PRE-INITIATION structure. Nucleotides 1-6 of the 3’ vRNA, which are not resolved, are colored in grey. Distal duplex and 5’ hook regions are labelled. (High-resolution figure available at the end of this PDF)

**Figure 3.**
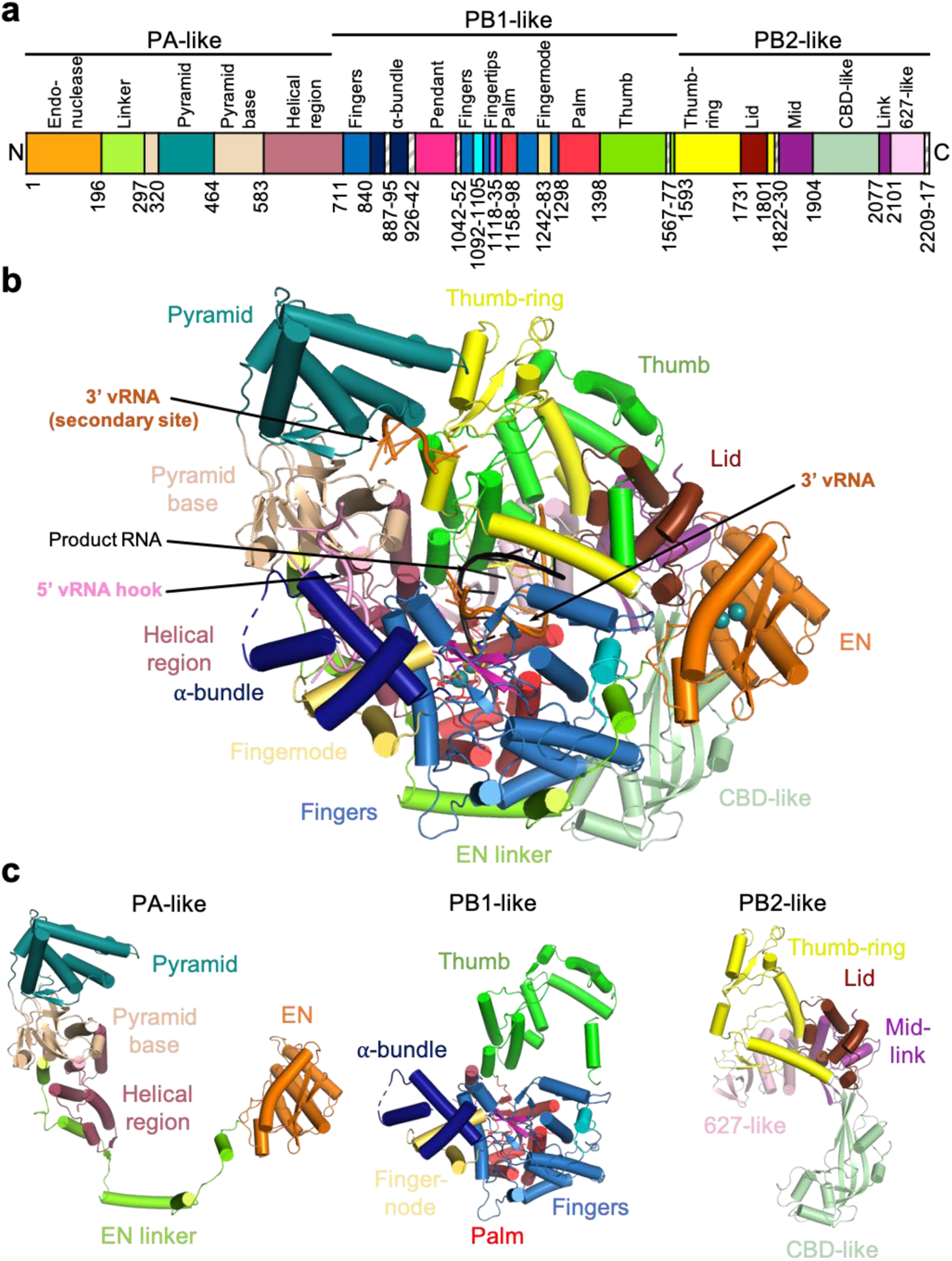
Overview of the L protein structure. **a.** Schematic linear presentation of the L protein domain structure. **b.** Complete ELONGATION structure of the L protein presented as a ribbon diagram in front view. Domains are colored according to (a) and labelled. 3’ vRNA is colored in orange, 5’ vRNA in pink and product RNA in black. See also Supplementary Movie 3 for a 3D impression of the L protein and its domains. **c.** Separate presentation of the PA-like, PB1-like and PB2-like regions of the L protein in the ELONGATION structure. (High-resolution figure available at the end of this PDF)

**Figure 4.**
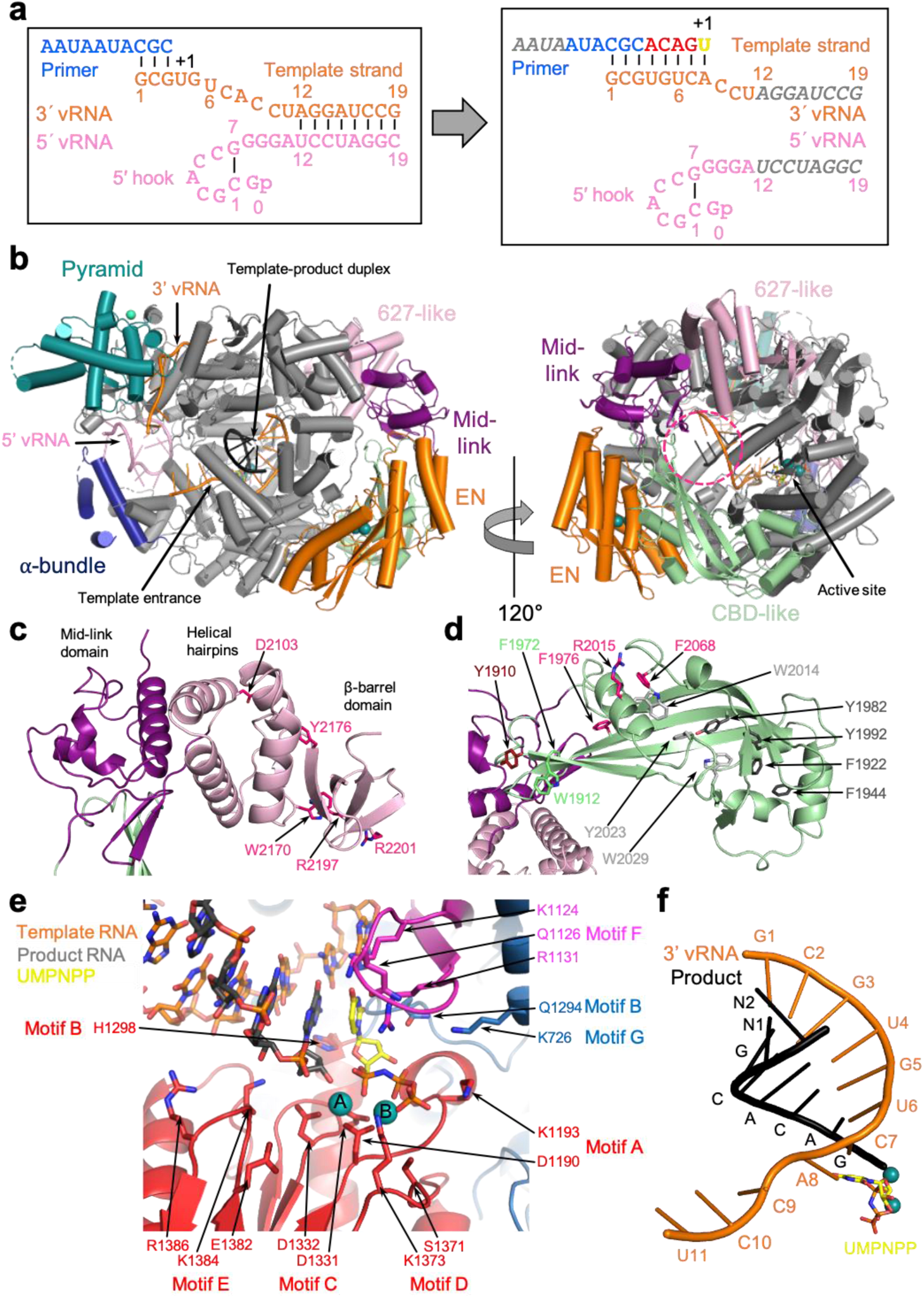
Elongation state of the L protein. **a.** Schematic presentation of the primed reaction carried out to obtain the ELONGATION structure with the L protein stalled in an early elongation state. Nucleotides that are not visible or not clearly assignable from the experimental map are shown in grey italics. **b.** ELONGATION structure of the L protein presented as a ribbon diagram in two views as indicated. EN, pyramid, α-bundle, mid-link, and 627-like domains are colored. 3’ vRNA is shown in orange, 5’ RNA in pink and product RNA in black. A dashed circle (hotpink) indicates the putative product exit. **c.** Close-up on the mid-link and 627-like domains with the respective structural features labelled and the side chains of amino acids shown to be selectively important for viral transcription by Lehmann et al. 2014 ^24^ shown as pink sticks. **d.** Close-up on the CBD-like domain with side chains of amino acids that have been tested in the LASV mini-replicon system shown as sticks (pink – selective role in viral transcription; light grey – no significant reduction of L protein activity upon mutation shown by Lehmann et al. 2014 ^24^; green – no or weak effect on L protein function upon mutation; dark red – potential selective role in viral transcription; dark grey – general defect of L protein activity upon mutation). Corresponding mini-replicon data are presented in Supplementary Fig. 8. **e.** Close-up of the polymerase active site with the template RNA (orange), the product RNA (dark grey), the non-hydrolysable UTP (yellow) and catalytic manganese ions (teal, A and B) involving the palm (red), fingers (blue), fingertips (magenta) and thumb (green) domains of the L protein. Important side chains are shown as sticks and conserved RdRp active site motifs (A-G) are labelled. **f.** Template-product duplex of the polymerase active site are shown as ribbon diagram with the product in black and the 3’ template in orange. (High-resolution figure available at the end of this PDF)

The different structures obtained (Table 1) reveal that LASV L, similar to influenza virus polymerase ^19^, has a number of domains flexibly linked to the polymerase core, allowing multiple configurations of the protein. No individual structure is complete (although the elongation structure model comprises ~90 % of the residues, lacking mainly the pendant domain), but integration of all the information leads to a coherent picture of the overall LASV L protein structure and the significant conformational changes that occur upon promoter binding and the subsequent transition into the active elongation state.

### Overview of LASV L protein structure

As previously established, MACV and LASV L proteins have an overall architecture similar to previously determined orthomyxovirus and bunyavirus polymerases ^9^. With reference to the heterotrimeric influenza virus polymerase, the LASV L protein can be conveniently divided into PA-like (1-687), PB1-like (688-1592) and PB2-like (1593-2217) regions (Figs. 3a and c, Supplementary Fig. 3).

The PA-like region has an N-terminal EN (1-195), whose structure and properties have previously been studied ^20,21^. Whereas previous crystal structures of the isolated LASV EN are not resolved beyond residue ~ 173, the full L protein structures show that the domain comprises an additional helix ending at 190. The EN is followed by an extended linker (196-296) that wraps around the polymerase core (Fig. 3, Supplementary Fig. 3). This connects to the pyramid base (297-319, 464-582) into which is inserted the prominent feature denoted the pyramid domain (320-463) (Figs. 1a and 3, Supplementary Fig. 3). The pyramid domain is specific to Old World arenaviruses and results from residue insertions compared to New World arenaviruses (e.g. MACV) that considerably lengthen the two helices spanning 386-435 (Supplementary Alignment file), giving it a characteristic angular shape. At the beginning of the pyramid domain, there is a structural zinc-binding site with coordinating ligands H316, C321, H364 and C366 (Supplementary Fig. 4). Mutational studies using the LASV mini-replicon system showed a general but incomplete reduction in L protein activity upon single site exchanges to alanine (Supplementary Fig. 4), which further emphasizes the structural role of this zinc-binding site. Indeed, sequence comparisons show that this site is specific to the LASV strain Bantou 289 and closely related strains, but not conserved in other LASV lineages or other arenaviruses (Supplementary Alignment file). Interestingly, the MACV L protein also contains a zinc-binding site but in a different location, at the pyramid base (Supplementary Fig. 4).

The PB1-like region (Figs. 3a, 3c, 4e, Supplementary Fig. 3) contains the canonical fingers, palm and thumb with associated conserved polymerase motifs A-F (motif G is 641-RY, motif H is K1237 ^13^). The catalytic triad of aspartates are D1190 (motif A), D1331 and D1332 (motif C). As previously noted ^9^, the fingertips (motif F, 1117-1137) are well structured even in the absence of bound promoter (Supplementary Fig. 5), unlike in influenza virus and LACV polymerases ^13^. The LASV L PB1-like region is considerably larger than influenza virus PB1 (882 residues compared to 756), mainly due to the so-called Lassa insertion (830-1069). This includes two flexibly linked modules: (i) a three-helix α-bundle (840-925), which includes an α-ribbon, and (ii) the compact pendant domain (943-1042), the latter being only partially visible in our structures (Fig. 3, Supplementary Fig. 3). The internal connection (887-895) between the α-ribbon and third helix of the α-bundle is disordered as are the flexible linkers before (926-942) and after (1042-1052) the pendant domain (Fig. 3a, Supplementary Fig. 6). These modules were previously observed in the MACV L structure ^9^, but in different positions and configurations (Supplementary Fig. 6). The PB1-like region has an additional insertion in the fingers, called the finger node (1242-1283), not present in influenza virus PB1, but very similar to the finger-node of LACV and likewise involved in binding the 5’ hook (see below) (Fig. 2d) ^13^. In all LASV structures, the region 1567-1577 is disordered. Moreover, it is so far unclear whether any protein segment might serve as a priming loop.

The PB2-like region (1593-2217) (Figs. 3a, 3c, 5c and 5d, Supplementary Fig. 3) has a similar overall organisation to influenza virus, LACV and Severe fever with thrombocytopenia syndrome bunyavirus (SFTSV) polymerases, with a ‘thumb-ring’, associated with the core and surrounding the thumb domain. Into the thumb-ring a ‘helical lid’ domain (1731-1800) is inserted, which in influenza virus polymerase forces strand-separation during RNA synthesis ^22^. This is followed by a short flexible linker to an array of C-terminal domains (1830-2217), visible at lower resolution in the **MID-LINK** structure (Supplementary Fig. 2), but fully buildable in the well-ordered **ELONGATION** structure (Figs. 3a-c and 4b-d). This includes the influenza-like split ‘mid-link’ domain (1831-1903, 2077-2100), into which is inserted the putative cap-binding domain (CBD-like, 1904-2076), followed by a 627-like domain that comprises two helical hairpins (2101-2168) and a terminal, compact β-barrel domain (2169-2208) (Figs. 3a-c and 4c-d, Supplementary Figs. 2 and 3). The mid-link and 627-like domains are juxtaposed in the same way in the **MID-LINK** and **ELONGATION** structures, suggesting that they are rigidly associated (Supplementary Fig. 2), although possessing some rotational freedom as a whole with respect to the thumb domain (Supplementary Fig. 7). In contrast, the CBD-like domain has considerable rotational freedom, with a difference of ~84° in its orientation with respect to the mid-link domain in the **MID-LINK** and **ELONGATION** structures, respectively (Supplementary Fig. 7). The C-terminal region was previously visualised in a dimeric form of MACV L protein (PDB:6KLH), but at insufficient resolution to build a correct model ^9^. The MACV cryo-EM density (EMD-0710) for the C-terminal region is fully compatible with the LASV C-terminal model and shows an extended configuration similar to that observed in the **MID-LINK** structure (Supplementary Fig. 7). The individual modules of the LASV C-terminal region have similar folds to that of the California Academy of Sciences reptarenavirus (CASV) L protein, despite low sequence homology (Supplementary Figs. 2 and 8) ^23^. However, the LASV CBD-like domain is considerably more elaborate, having ~170 residues compared to ~ 100 residues in CASV (Supplementary Fig. 8). The CASV CBD-like domain is minimalist, comprising a five-stranded mixed β-sheet with a transverse helical hairpin. In LASV, there are significant insertions in the β1-β2 and β3-β4 loops and in the loop of the helical hairpin, that fold together to extend the length of the domain (Fig. 4d, Supplementary Fig. 8). In the **ELONGATION** structure, the CBD-like domain is locked in position by interactions with the EN and core of the polymerase, with the helical hairpin insertion being particularly important for the latter interaction (Supplementary Fig. 9). In this conformation, the canonical cap-binding site between the C-terminal end of strand β1 and the principal transverse helix, as observed in cap-bound CBDs of SFTSV, Rift-Valley fever virus (RVFV) or influenza virus (Supplementary Fig. 8), appears to be partly blocked. Moreover, the CBD-like and 627-like helical-hairpin domains are particularly poorly conserved across arenavirus L proteins (as opposed to the mid-link and C-terminal β-barrel domain) (Supplementary Alignment file, Supplementary Fig. 8). Indeed, the most conserved region of the CBD-like domain is at the N-terminus of strand β1 (1909-GYAW in LASV). Past and recent mutational analyses of the potential cap-binding aromatic residues of this domain using the LASV mini-replicon system did not identify any transcription specific residues that might be responsible for cap-binding (Fig. 4d, Supplementary Fig. 8) ^24^. Neither have *in vitro* studies with isolated soluble domains of the CASV and LASV L protein C-terminal region detected any cap-binding activity ^23^, unlike for phenui- and orthomyxoviruses ^25–27^. It therefore remains an open question, whether in some, yet to be observed configuration of the L protein, a functional cap-binding site is formed.

**Figure 5.**
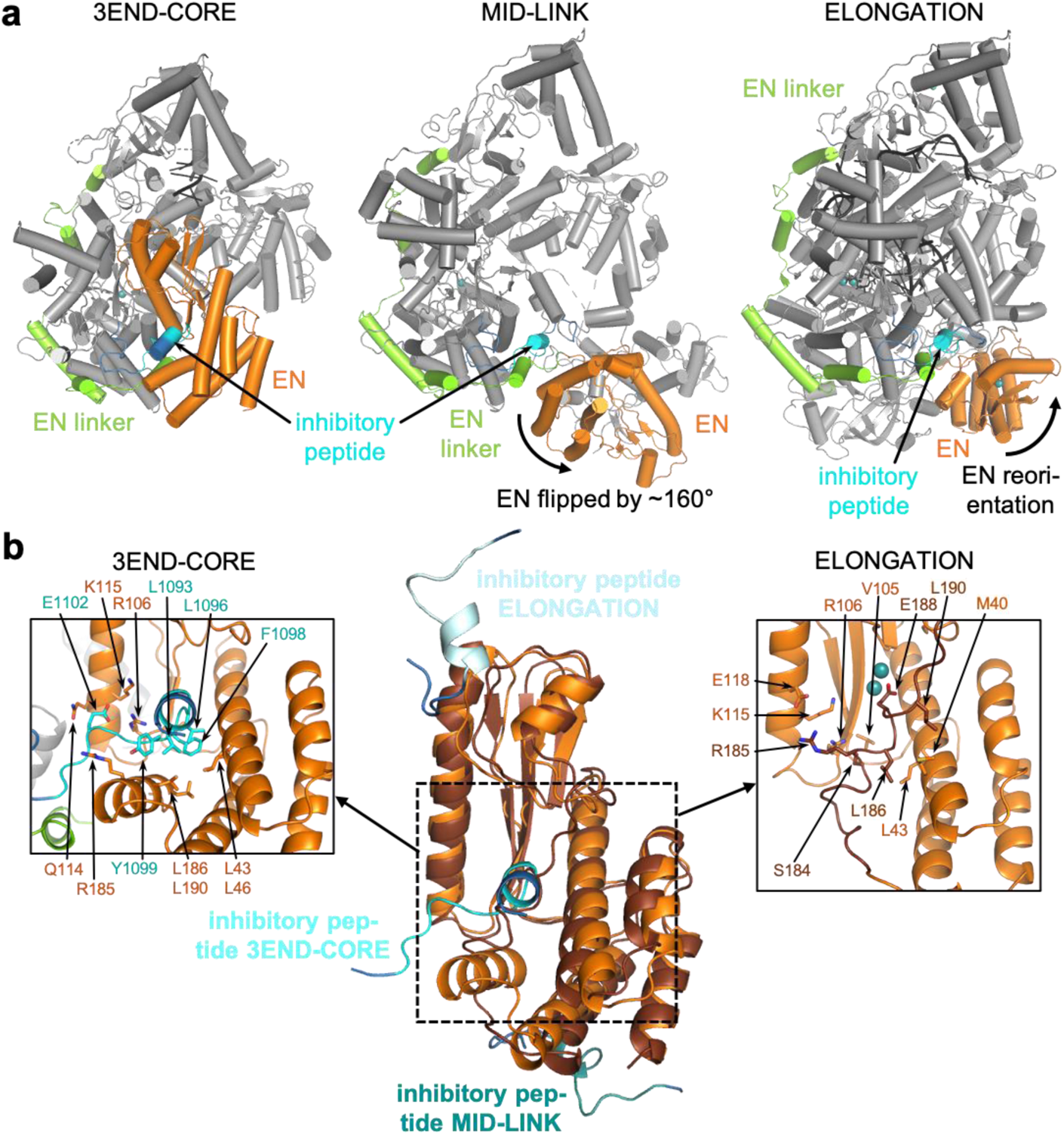
Conformational flexibility of the endonuclease domain. **a.** Overview of the three different conformations of the EN (orange) observed in the 3END-CORE, MID-LINK, and ELONGATION structures. The EN linker (green) and the inhibitory peptide (cyan) are highlighted as well. **b.** In the middle panel a superimposition of the EN domains of 3END-CORE (orange), corresponding also to the overall conformation of the EN domain in the MID-LINK structure, and ELONGATION (brown) with the respective positions of the inhibitory peptides in teal, cyan and light blue, respectively, is shown. Connections to the fingers domain are indicated in blue. The position of the inhibitory peptide of 3END-CORE is the same as in the APO-ENDO structure, similarly is the same position of the EN observed in both MID-LINK and DISTAL-PROMOTER structures. Right and left panels show close-ups of the autoinhibited EN active sites in the ELONGATION and 3END-CORE structures, respectively. Important residues of the protein:protein interactions are labelled and side chains are shown as sticks. (High-resolution figure available at the end of this PDF)

### Positional flexibility and regulation of the endonuclease

We observe three quite different locations of the EN, in each case differently packed against the core of the L protein (Fig. 5). In the **APO-ENDO** and **3END-ENDO** structures, the EN interacts with the core regions 1137-1142 and 1592-1604 (Supplementary Fig. 10). Mutation to alanine of F1592, which packs on P109 of the EN, resulted in a slight reduction in general L protein activity in the LASV mini-replicon system. A severe, general loss of 90% of activity was observed for L protein mutant P109G (Supplementary Fig. 10). The EN active site itself is exposed to the outside, but access is blocked by residues 1092-1105 from the PB1-like region, which we refer to as the ‘inhibitory peptide’ (Figs. 5a and b). The arena-conserved 1096-LCFYS motif (Supplementary Alignment file) is intimately bound in the EN active site pocket (Fig. 5b, Supplementary Figs. 10 and 11) and would thus prevent any substrate RNA binding there (e.g. superposition of LCMV EN structure with bound inhibitor shows overlap ^28^) (Supplementary Fig. 11). A similar interaction with EN is observed in the apo-MACV structure (PDB:6KLD) (Supplementary Fig. 4).

In the **MID-LINK** and **DISTAL-PROMOTER** structures, the EN has flipped by ~160° around a hinge between G195-G199. The EN active site faces away from the rest of the polymerase and is exposed to the solvent, free of the inhibitory peptide (Figs. 5a and b). Instead, inhibitory peptide residues 1087-1099, as well as PB2-like segments 1759-1770 (lid), 1852-1860 and 1894-1896 (mid), 2077-2081 and 2089-2091 (link) pack against the back of the EN, stabilising it in its new location. For these two observed positions of the EN, the total buried surface area is comparable, 3005 Å^2^ (**APO-ENDO**, autoinhibited) and 2721 Å^2^ (**MID-LINK**, free), compatible with there being an equilibrium between the two states as observed in the two different apo-structures.

In the **ELONGATION** structure, the EN is stabilised in a third position (Figs. 5a and b) with its active site auto-inhibited by a completely different mechanism involving the C-terminal region of the EN (173-190). This is redirected so that the 181-188 helix binds in and blocks the EN active site groove, with E188 co-ordinating, together with E51 and D89 ^21^, two cations in the active site (Fig. 5b, Supplementary Fig. 10). The inhibitory peptide remains at the same place with respect to the polymerase core, but due to the reorientation of the EN it packs against a different site on the EN, with, for instance, K1094 making a salt-bridge with E70 and Y1099 stacking against P81 (Fig. 5b, Supplementary Fig. 11). Diverse other regions of the L protein also contact the EN (Fig. 3b, Supplementary Fig. 9) and the total buried surface area of the EN in this location is 4060 Å^2^.

To investigate the function of the inhibitory peptide, we performed a mutational analysis of residues 1092-1105 as well as interacting residues of the EN domain, as observed in the **APO-ENDO** and **3END-CORE** structures (Supplementary Figs. 10 and 11). We observed a severe general defect in L protein function for a number of mutants both in the EN domain (L43G/N, L46G/N, V105G, R106K, K115A, R185A, L186G, L190G/N) and in the ‘inhibitory peptide’ (L1093S, L1096A/N, C1097G, F1098A/S, Y1099A, E1102A) (Supplementary Fig. 11). For hantavirus L protein it was shown that an active EN can lead to RNA degradation and therefore lower protein expression levels ^29^. To exclude that the general defect of the mutants of the ‘inhibitory peptide’ interaction is caused by RNA degradation due to an elevated activity of LASV EN, we combined the previous mutations with the EN inactivating mutation D89A ^30^, without observing any change in phenotype (Supplementary Fig. 12). Additionally, using *in vitro* polymerase assays only residual polymerase activity was detectable for L protein mutant E1102A and no activity for mutant Y1099A (Supplementary Fig. 13). We conclude that the ‘inhibitory peptide’ and other tested residues involved in the interaction play a general role in L protein activity but are not selectively important for transcription, this being consistent with the diversity of interactions we see for these residues when comparing all observed conformations of the L protein (Fig. 5). Comparing the apo- and pre-initiation structures suggests that either promoter binding or mutations in the inhibitory peptide release the EN from autoinhibition. We tested this hypothesis by assaying purified L proteins with mutations in the inhibitory peptide for EN activity *in vitro*, using capped or uncapped RNA substrates and with either no promoter, 3’ end only, 5’ end only or both promoter RNAs present (Supplementary Fig. 14). The EN active site mutant E102A and addition of the nuclease inhibitor DPBA served as negative controls. We found that, for the wild-type L protein, the only situation where weak EN activity is reproducibly detectable is when the 5’ end only or both promoter ends are bound, and the same is true for the L protein with mutations Q114A or E1102A, which are probably not sufficient to disrupt inhibitory peptide binding.

In summary, in both the apo- and early elongation states, the EN is autoinhibited, but by different mechanisms involving binding in the active site of either the ‘inhibitory peptide’ 1092-1105 or the C-terminal helix of the EN, respectively. Whilst 5’ end or full promoter binding partly activates the EN, consistent with the structure and presumed functional role in cap-dependent transcription priming of the pre-initiation state, its low intrinsic activity *in vitro* under any conditions tested by us and others ^20,21,31^ suggests that the mechanism of EN activation may be more complex than expected from the currently available structural data.

### 3’ end binding in the secondary site

Incubation of LASV L protein with vRNA 3’ end nucleotides 1-16 yielded the currently highest resolution structure (**3END-CORE**, 2.70 Å). It features specific binding of nucleotides 1-7 in a tunnel under the pyramid (Figs. 1b and c). This site corresponds to the secondary 3’ end site previously observed for influenza virus polymerase ^16,17^, LACV L ^13^ and MACV L ^9^. The excellent cryo-EM density enables unambiguous base identification of nucleotides 2-5 (CGUG) of the 3’ end and placing of several water molecules in the protein-RNA interface (Supplementary Fig. 15). However, G1 and nucleotides 6-7 (UC) have poor density. In the MACV L structure, G1 is better defined perhaps due to its stabilisation by stacking on Tyr534, which is substituted by Leu540 in LASV L (Supplementary Alignment file). Nucleotides 3-5 of the 3’ end form a particularly compact arrangement with a direct interaction between G3 O6 and G5 N2, and U4 stacking underneath (Fig. 1c, Supplementary Fig. 15).

Several specific protein-RNA interactions are made with conserved arenavirus residues such as K332, D380, L502, K509 from the pyramid and Y1450, R1452 and S1626 from the thumb and thumb-ring domains, thus involving the PA-, PB1 and PB2-like regions, as in influenza virus and LACV (Fig. 1c). Mutational analysis of the residues interacting with the 3’ end in the secondary binding site using the LASV mini-replicon system revealed a general defect in L protein function upon introduction of double mutations N331A/K332A and Y1450A/R1452A, whereas mutations L502A, K509A and R1622A did not interfere with L protein activity (Fig. 1d). Purified L protein mutant Y1450A/R1452A exhibited significantly reduced 3’ end binding ability compared to wild-type L protein (Fig. 1e). However, this mutant maintained polymerase activity in the presence of the 19 nt 3’ and 20 nt 5’ promoter RNAs and, in contrast to the wild-type L protein, showed polymerase activity with only the 19 nt 3’ promoter RNA present (Supplementary Fig. 16). This strongly suggests that in the wild-type L protein, in the absence of the 5’ end, the 3’ end is tightly sequestered in the secondary binding site and does not enter the active site (see discussion). In the presence of a 47 nt hairpin RNA containing the connected 3’ and 5’ promoter sequences of LASV, L-Y1450A/R1452A showed significantly reduced polymerase activity (Supplementary Fig. 16). This shows that 3’ end binding in the secondary site is required for efficient RNA synthesis, either to sequester the template 3’ end after passing through the active site and/or to prevent the unbound 3’ end from forming double stranded RNA with the template 5’ end or the product RNA.

### Full promoter binding

The **PRE-INITIATION** structure shows the full promoter (5’ nts 0-19, 3’ nts 1-19) bound to the LASV L protein and reveals several significant conformational changes that occur upon promoter binding (Fig. 2 Supplementary Fig. 1). The LASV vRNA 5’ and 3’ ends are highly complementary over 19 nucleotides with only two mismatches at positions 6 and 8 (Supplementary Table 2). In addition, the 5’ end carries an extra nucleotide (G0) arising from a prime-and-realign mechanism during replication initiation ^4,5,32–34^. As expected, when bound to the L protein, the promoter does not adopt a fully double-stranded conformation but forms a structure resembling that observed for influenza virus and LACV (Figs. 2a, c, d and e, Supplementary Fig. 17). Nucleotides 12-19 from both strands form a distal 8-mer canonical A-form duplex, whereas nucleotides 1-11 of each end are single-stranded, which is consistent with previous mutational studies ^35^. Nucleotides 0-9 of the 5’ end form a compact hook structure linked to the duplex region by nucleotides 10-11. The internal secondary structure of the hook differs from that of influenza virus and LACV polymerases by only having one canonical base-pair (C1-G7) upon which G2 and C3 are consecutively stacked on one side and G8 and G9 on the other side (Fig. 2d, Supplementary Fig. 17). The loop of the hook comprises nucleotides 4-6, with A4 stacking on C6. The hook makes extensive interactions with conserved residues from numerous different loops from both the PA-like and PB1-like regions of the L protein (e.g. residues 470-474, 518-526, 860-861 and 1255-1265) (Fig. 2d, Supplementary Movie 1). Several of these loops only become structured upon promoter binding. Of note, three aromatic residues are involved, Y471 (conserved in Old World arenaviruses only), Y526 (Y or F in all arenaviruses) and Y574 (conserved in all arenaviruses) (Supplementary alignment file). Y471 interacts with the phosphate of G10 and Y574 with the base of G9. Y526 extends and stabilises the central stacked backbone of the hook by packing on G9, with I665 playing a similar role on the other end by stacking against C3. G0 is base-specifically recognised by the carbonyl-oxygen of S470 and is sandwiched between P297 and R473, which could potentially interact with the terminal triphosphate (not present in the 5’ RNA used for this structure) (Supplementary Fig. 17). Mutational analysis of the 5’ binding site using the LASV mini-replicon system reveals that most mutations cause a severe general defect in L protein activity (Supplementary Fig. 1). Complete loss of function was observed for mutants Y474A, V514G/K515A, R525A/Y526A and K681A. An activity reduction to ~20% was observed for mutants R473A/T474A, Q551A/K552A, Y574A, and K1263A/T1265A. These results indicate that 5’ end binding is important for the general function of LASV L protein during both viral transcription and replication.

The 3’ strand of the distal duplex specifically interacts with two regions of the L-protein pyramid (Figs. 2b and 6a). One is the 340-loop, which orders upon binding, with the arenavirus-conserved motif 340-NTRR making several major groove contacts with bases and phosphates of the 3’ strand nucleotides 12-15 (and R337 with phosphate of C10). The second interacting region involves the Old World arenavirus-specific extended helices at the top of the pyramid, including W395, R399 (conserved in all Old World arenaviruses) and K423 contacting the backbone phosphates 3’ strand nucleotides 16-17 (Fig.2b, Supplementary Movie 1). The single-stranded 3’ end nucleotides 11 to 7 are directed towards the polymerase active site, but 1-6 are not visible (Figs. 2a, c and e). Arenavirus-conserved R1561 interacts base-specifically with C9, which also stacks on chemically conserved R1622, both residues being from the thumb domain (Fig. 2b, Supplementary Movie 1).

**Figure 6.**
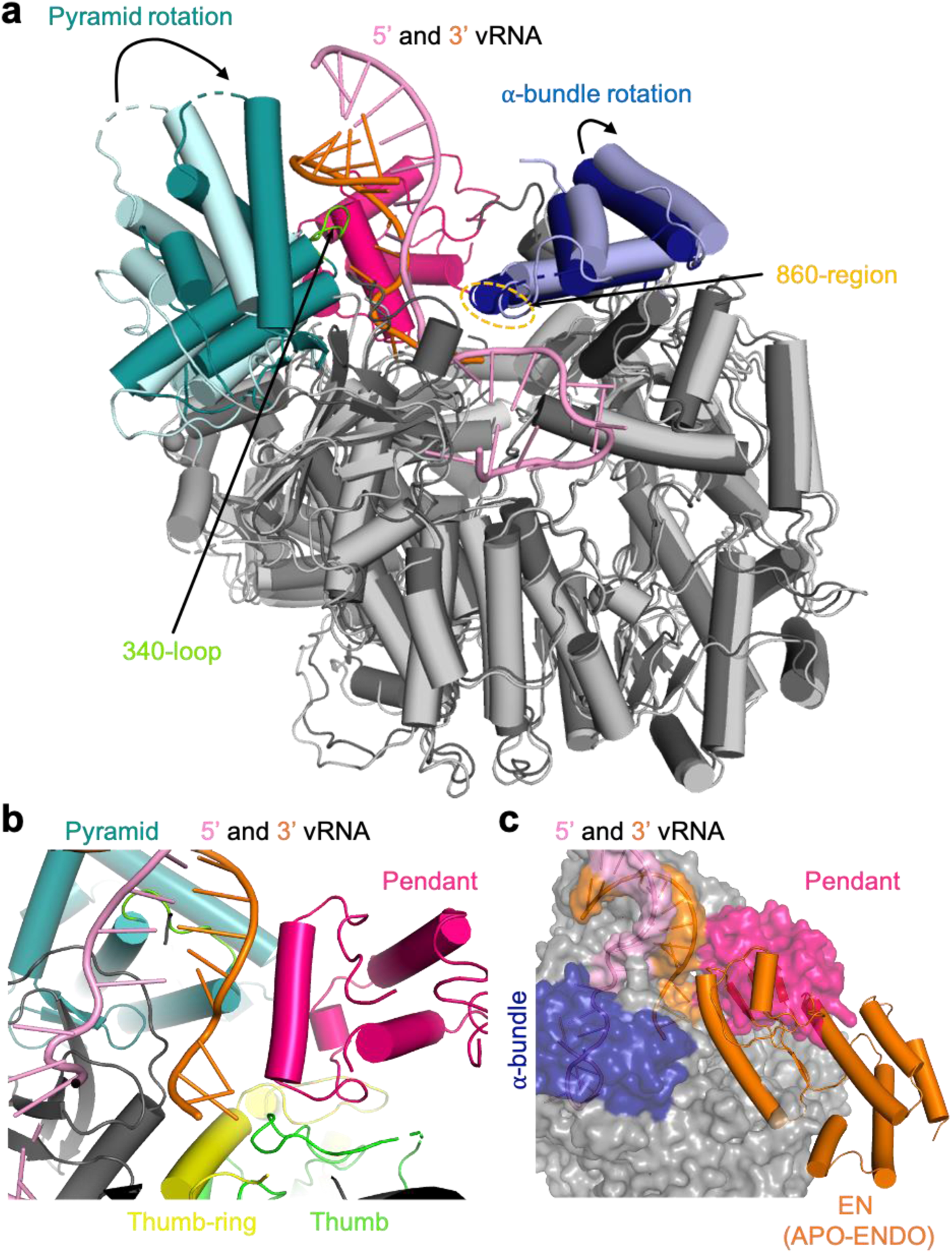
Global rearrangements upon promoter binding. **a.** PRE-INITIATION (dark grey, teal, dark blue, hotpink) and APO-RIBBON (light grey, light cyan, light blue) structures are superimposed. The pyramid and α-bundle rotations between apo- and promoter-bound structures are indicated as well as the 340-loop and the 860-region. Promoter RNA is shown in pink and orange. **b.** Close-up of the interaction site between promoter RNA (pink and orange) and the pendant (hotpink), thumb-ring (yellow) and thumb (green) domains in the PRE-INITIATION structure. Pyramid domain shown in teal. **c.** Superposition of the PRE-INITIATION and APO-ENDO structures. PRE-INITIATION is presented as transparent surface in grey with the pendant (hotpink) and α-bundle (blue) as well as the 5’ (pink) and 3’ (orange) vRNA highlighted in color. The EN domain of the APO-ENDO structure is shown as orange ribbon, which overlaps with the pendant domain volume of the PRE-INITIATION structure. (High-resolution figure available at the end of this PDF)

Apart from the induced fit ordering of several promoter-interacting loops, there are also more global rearrangements. Firstly, distal duplex binding causes a major rotation of the entire pyramid by ~21.6°, enabling the summit helices to contact the 3’ strand as described above (Fig. 6a). This rotation does not occur when just the 3’ end is bound in the secondary site. Secondly, the α-bundle rotates slightly (4.5°, **PRE-INITIATION** versus **APO-RIBBON**) to allow interaction of the 860-region (860-861) with the 5’ hook (Fig. 6a). Thirdly, the pendant domain (943-1040) becomes stably positioned by packing against the rotated pyramid domain and the thumb and thumb-ring (Fig. 6b). The pendant domain helix spanning residues 953-965, runs parallel to 3’ nucleotides 9-12, but with only two direct interactions. R957 contacts the phosphate of A12 contributing to stabilisation of the 3’ end interacting 340-loop and K960 is close to phosphate of U11 (Fig. 2b). The pendant domain was first visualised in the MACV apo-L structure in a very different position that would superpose with the promoter duplex region as well as the α-bundle, as observed in the promoter-bound LASV L structure (Supplementary Fig. 6). Similarly, the α-bundle of MACV apo-L (with its different topology, see Supplementary Fig. 6) superposes with the pendant domain in LASV L, showing that these flexible and linked domains must rearrange from the apo-state observed in MACV L. Finally, the position of the pendant domain in the LASV **PRE-INITIATION** structure is incompatible, due to significant steric overlap, with the location of the inhibited EN in the **APO-ENDO** or **3END-ENDO** structures (Fig. 6c). This important observation provides a plausible rationale for full promoter binding (as opposed to just the 3’ end binding in the secondary site) inducing a flip of the EN to the alternative uninhibited location observed in the **MID-LINK** and **DISTAL-PROMOTER** structures (see above).

### Elongation structure

To determine a structure of functionally active LASV polymerase in early elongation, we incubated promoter bound L protein with a 10-mer uncapped primer, 5’-AAUAAUACGC-3’ together with ATP, GTP, CTP and non-hydrolysable UMPNPP (denoted U_PNPP_). Biochemical analysis (Supplementary Fig. 18) shows that various products are formed depending on whether (i) the 3’ terminal triplet of the primer hybridises with the 3’ end of the template (3’-GCGUGUCA…) giving 14-mer 5’-AAUAAUA**<u>CGC</u>**ACAGU_PNPP_ or 18-mer: 5’-AAUAAUA**<u>CGC</u>**ACAG[U]GGAU_PNPP_ products or (ii) just the 3’ terminal nucleotide hybridises, giving 16-mer 5’-AAUAAUACG**<u>C</u>**GCACAGU_PNPP_ or 20-mer 5’-AAUAAUACG**<u>C</u>**GCACAG[U]GGAU_PNPP_ products (here, bold indicates primer hybridization, red colouring: incorporated nucleotides, [U] misincorporation at an A in the template) (Supplementary Fig. 18). In each case, the longer product is formed if the A8 in the template is read-through by misincorporation. The most prominent products are the 14-mer and 18-mer. Upon plunge freezing on EM grids and performing 3D single particle reconstruction, the sample gave a major class showing the stalled (i.e. pre-incorporation), early elongation state at 2.92 Å resolution. The high quality of the density allows unambiguous assignment of the template and product sequences. As expected, incoming UMPNPP is observed at the +1 position base pairing with A8 of the template. There is good density for seven bases of the product (5’-….N_1_N_2_CGCACAG), with the preceding two, which are unpaired with the template, having poorer density, prohibiting unambiguous identification (Supplementary Fig. 19). However, since the map density corresponding to N_2_ looks more like an A, the product is most likely the 14-mer. Products that would contain mismatches due to read-through, may be less stably bound to the polymerase. The active site cavity contains an 8-mer duplex from position +1 to −7 (+1 corresponding to the UMPNPP) (Figs. 4a, e and f), whereas as observed for related viral polymerases ^14,22^ a 10-mer duplex (positions +1 to −9) is expected to fill the active site cavity before strand-separation. Nucleotides 1-11 of the template are visible, corresponding to positions +4 to −7. The duplex region of the promoter has melted due to translocation of the template but nucleotides 0-11 of the 5’ end remain bound in the hook conformation as described above. Consistent with the distal duplex being absent, the pyramid is not rotated and a second template 3’ end (nucleotides 1-7) is actually bound in the secondary site (Fig. 4b). Whilst this is likely an artefact of performing the RNA synthesis reaction with excess template, it does confirm that secondary 3’ end binding is compatible with elongation and consistent with the template docking in this site after exiting the active site cavity, as observed for influenza virus polymerase ^18^. The pendant domain is not visible and the α-bundle only has weak density, probably due to the absence of the distal promoter duplex.

The configuration of the polymerase active site as well as the binding of the incoming nucleotide and template are canonical, involving conserved motifs A-D and the fingertips (xmotif F) (Fig. 4e, Supplementary Movie 2). The triphosphate and terminal 3’ OH of the product strand are co-ordinated by two manganese ions (A and B), held in place by D1190 (motif A) and 1331-DD (motif C). K1373 of motif D also contacts the γ-phosphate. Motif F residues R1131, positioned by Q1294 (motif B), and L1133 stack under the incoming nucleotide template bases at the +1 position, respectively, while K1124 (motif F) contacts the O4 of the incoming nucleotide (Supplementary Movie 2).

Compared to the unoccupied active site in the PRE-INITIATION structure, only minor adjustments to the active site loops occur, the most significant being displacement of the central β-strands of the fingertips loop by about 2 Å to make room for the +1 base-pair to stack on R1131 and L1133 (Supplementary Fig. 20). Unlike in influenza virus polymerase ^22^, motif B does not change conformation between the occupied and unoccupied states of the active site. However, to accommodate the growing template-product duplex, the helical lid (1731-1805) has to be displaced out of the active site cavity by about 8 Å (Supplementary Fig. 20). Coupled with the lid movement, the sharply kinked (requiring conserved Gly1595) pair of consecutive helices α52-α53 (1579-1611) also translate in the same direction, with helix α53 forming one side of the active site cavity close to the distal part of the product strand. Interestingly, the first visible base of the product (position −9) is packed against T1583 from α52, which could therefore play a role in strand separation rather than the helical lid itself (Supplementary Fig. 20).

More generally, the conformation of the active elongating polymerase is stabilised by a number of new interactions between distant regions of the L protein sequence. For instance, in the new position of the helical lid, residues 1764-1766 contact the EN at Phe85 (close to the EN active site), contributing to the interactions which stabilise the EN in its third location (see above). Residues 811-820, disordered in all other structures, interact with EN linker 195-199, again only possible with the EN in its new location. Similarly, the inhibitory peptide residues 1087-1091 change conformation to allow simultaneous interaction with the EN (1089-TT with D129 and S82) and with the kink between α52-α53 (A1091 with T1591) (Fig. 5a, Supplementary Fig. 11). Most dramatic, is the stabilisation of the entire C-terminal domain, which, together with the exposed end of the palm, forms a ring with a ~ 30 Å diameter central pore, a putative product exit channel (Fig. 4b). The EN buttresses the proximal part of the CBD-like domain (e.g. Q32 with A1911) as well as the mid-link domain (e.g. A171 with K1895, E34 with K1891), whereas the distal part of the CBD-like domain interacts with numerous loops from the polymerase core including EN linker residues 230-232 (e.g. H232-Q2045), fingers domain residues 793-799, 802-805 (e.g. V802-Y2030), 1215-1216 (e.g. D1216-K2062, K1215-E2053), and palm domain residues 1314-1318 (e.g. Y1314-Q2045) (Supplementary Fig. 9). The total buried surface area between the polymerase core and the CBD-like domain is 2884 Å^2^. The extreme C-terminal 627-like domain (mainly the β-barrel and to a lesser extent, the helical hairpins) make interactions with multiple regions, notably 1715-1722 and 1812-1816 of the thumb-ring (e.g. F1715-Y2176/V2145, F1716-V2189, D1722-S2191/S2192, R1816-D2143, S1812/L1815-G2175), residues 691-694 of the helical region (e.g. M691-G2193) and the palm domain 1390-1392 loop (e.g. W1390-R2197) (Supplementary Fig. 9). The total buried surface area of the β-barrel domain is 1472 Å^2^. Interestingly, W2170, R2197 and R2201, whose mutation leads to a transcription-specific defect ^24^, are intimately involved in the interface together with W1390 in the 1390-loop. Even though mutation of W1390 to alanine in a previous study did not impair overall L protein activity ^36^, from the structure we would not expect a small hydrophobic alanine residue to disturb the remaining contacts between these domains. Additionally, residues G1391 and D1392 were shown to be selectively important for viral transcription ^36^, further emphasizing the importance of this interaction site. Residue Y2176, also identified as being selectively important for viral transcription ^24^ interacts with the thumb-ring residue F1715 (Supplementary Fig. 9). These data suggest that the configuration of the C-terminal region and its interaction with the core, as observed in the **ELONGATION** structure is critical for transcription.

## Discussion

Previous biochemical studies on LASV L ^7^ and MACV L ^8,37^ proteins have revealed certain features of promoter binding to arenavirus polymerases and the impact on RNA synthesis activity. In RNA binding experiments, it was shown that MACV L makes a tight complex with the 3’ promoter strand with the identity of nucleotides 2-5 being particularly important ^8^. This corresponds exactly with the binding specificity of the 3’ end secondary site seen in our structural analysis. For both MACV and LASV, the most efficient *in vitro* RNA synthesis activity was observed using both 19-mer strands in 1:1 ratio as in the native promoter ^7,37^. For MACV L protein weak activity, which could be enhanced with a GpC primer, was also observed in presence of only the 3’ strand ^8^. The more specific requirements found for optimal unprimed RNA synthesis by LASV L were (i) the presence of the terminal non-templated G0 base on the 5’ strand (i.e. 0-19); (ii) the two mismatches at positions 6 and 8 in the S segment promoter (only one in the L segment), rather than a perfect proximal duplex; (iii) a sufficiently long distal duplex region, preferably the full 19-mer ^7^. For MACV, it was further shown that the G0 phosphates were not essential and that enhancement was achieved with 3’ truncated 5’ ends (e.g. maximal activity for 0-12 mer), although these experiments differed in that a GpC primer was systematically used ^37^. These observations are consistent with our structural analysis as well as the notion that the default mode of binding of the 3’ end alone is in the secondary site and that full-promoter binding or a primer is required to dislodge it and permit RNA synthesis. More recently, it has been confirmed for both LASV and MACV L that *de novo* RNA synthesis is indeed enhanced by the presence of the 5’ end, but, surprisingly, it was reported that cap-dependent transcription was inhibited by the 5’ end ^9^. This raises the question of the exact role of the 5’ end binding in arenavirus L proteins.

In LACV and influenza virus, 5’ end hook binding is required to order the fingertips loop into a functional configuration in the polymerase active site ^13,14^. This does not appear to be the case for arenavirus L proteins ^9^ (Supplementary Fig. 5). In addition, for influenza virus it has been shown that 5’ end binding stimulates EN activity, probably by favouring the transcription active configuration of the polymerase over the replicase conformation ^19^. Evidence given above that 5’ end (or full promoter) binding stimulates EN activity suggests that a similar conformational change mechanism may operate for arenavirus L proteins. Finally, whereas 5’ end binding is required in orthomyxoviruses for poly(A) tail generation during transcription ^18^, arenavirus L proteins terminate transcription by a very different mechanism without poly(A) tail synthesis ^38–41^. To investigate the functional consequences of 5’ end hook binding further, we performed polymerase activity assays either (i) with the 3’ promoter end alone, (ii) the 3’ end together with the 5’ (nts 0-19) end or (iii) the 3’ end together with a truncated 5’ (nts 0-12) end. In each case, the assays were performed in the presence or absence of 3 nt or 10 nt long uncapped primers (Supplementary Fig. 21). We used uncapped primers as we could not detect any difference in primed product formation between uncapped (hydroxylated or tri-phosphorylated 5’) and cap0-capped primers for LASV L (Supplementary Fig. 22). Under the conditions of the reaction, no products were formed by the 3’ end alone unless a primer was present (Supplementary Fig. 21). Adding the promoter 5’ end (nts 0-19) led to strong product formation even in the absence of primer. When the 5’ end was truncated (nts 0-12), unprimed product formation was significantly reduced, whereas primed product formation was comparable to the corresponding conditions with the full 5’ (nts 0-19) RNA. In the absence of the 5’ end, structural and biochemical data show that the 3’ end preferentially binds tightly in the secondary binding site. Bearing this in mind, we interpret our activity results to show that 5’ end binding and/or the presence of a primer (capped or uncapped), that can hybridise with and stabilise the 3’ end in the polymerase active site, stimulates RNA synthesis activity, presumably by promoting 3’ end binding in the active site rather than the secondary site. Indeed, the major rotation of the pyramid domain induced by distal duplex binding shears the two sides of the 3’ end-binding groove and prevents closure around the RNA, thus disfavouring secondary site 3’ end binding when the full promoter is bound. Since, the promoter duplex no longer exists during elongation, this mechanism does not prevent the template 3’ end rebinding in the secondary site after going through the active site as observed for influenza polymerase ^18^.

In conclusion, our structural and functional results support the hypothesis that full-promoter binding, including the 5’ hook and distal duplex, induces the functionally ready pre-initiation state via the conformational changes described above, at the same time as releasing the EN from autoinhibition (Fig. 7). Given that our activity results are independent of whether the primer is capped or uncapped (Supplementary Fig. 22), we do not think they shed light on true cap-dependent transcription per se. It remains unclear, whether the assays presented by Peng et al. truly reflect cap-dependent transcription, as the respective control (i.e. non-capped primer) was not included ^9^. Indeed, we have not been able to recapitulate true cap-dependent transcription with the LASV polymerase, due to its weak or non-existent endonuclease and cap-binding activities, emphasising that the mechanism of cap-snatching and cap-dependent transcription for arenavirus L protein remains enigmatic. This is even more remarkable considering the shortness of the capped primers used ^3^, which it is difficult to imagine being bound in a canonical way by the CBD-like domain as well as reaching into the active site to hybridise with the template, a situation reminiscent of Thogoto virus polymerase (Guilligay et al, 2014).

**Figure 7.**
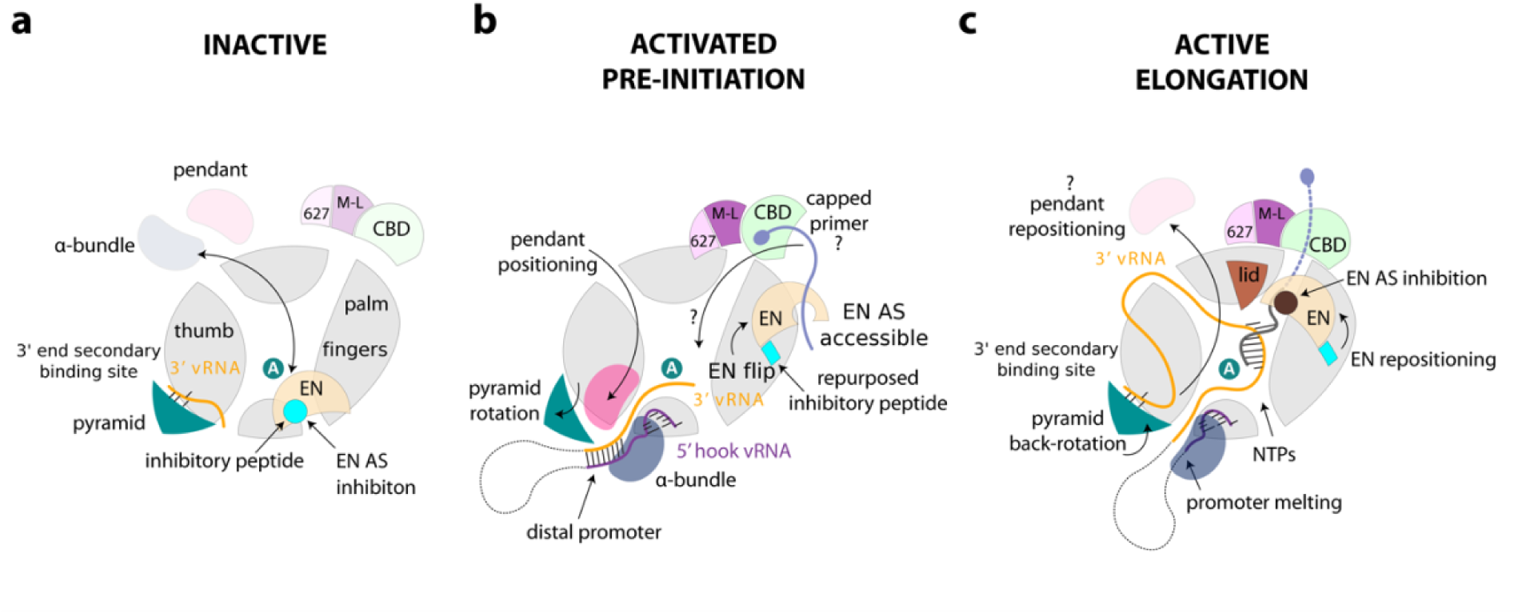
Schematic diagram of conformational changes in Lassa virus L protein associated with promoter binding and RNA synthesis activity. **a.** In the inactive state, the Lassa virus L protein tightly binds to the 3’ vRNA end base-specifically into a district secondary 3’ end binding site in-between the pyramid and thumb domain. On the surface of the L protein core, there is mutually exclusive positioning (black double-arrow) of either the α-bundle and pendant or the EN domain. When placed on the core, the EN domain is however autoinhibited by the inhibitory peptide in this configuration. b. Upon full promoter (i.e. 3’ and 5’ vRNA with a distal duplex region) binding, major conformational changes occur. Positioning of the distal promoter favours tight association of the α-bundle and pendant, and this together forces the pyramid domain to rotate, which is incompatible with 3’ vRNA in the secondary 3’ end binding site. The 3’ vRNA in the full promoter is extending towards the L protein active site (marked by the white A in the teal sphere). The 5’ end nucleotides 1-10 are bound in a hook-like conformation outside of the active site in a specific groove. The EN flips and binds to the other end of the L protein core via the inhibitory peptide, which also repositions and refolds for this purpose. In this configuration, the EN is in close vicinity to the CBD-like domain. The EN active site is accessible and could potentially cleave an incoming capped primer. How the capped primer is associated to the CBD-like domain and how it is navigated towards the active site remains elusive. c. Upon transition to the elongation state, the distal promoter duplex melts and the pendant domain is released, which allows the pyramid to rotate back and re-establish the secondary 3’ end binding site. We presume the 3’ vRNA template, after exiting the active site, wraps around the L protein core and rebinds to the secondary 3’ end binding site. The EN repositions once again and together with the mid-link and CBD-like domains forms a distinct ring around the putative product exit channel. The EN active site is again autoinhibited but by another peptide than observed in the inactive conformation. (High-resolution figure available at the end of this PDF)

The autoinhibited EN conformations appear in L protein configurations where the EN activity is not expected to be required and indeed could potentially be detrimental (Fig. 7). On the other hand, in the promoter bound pre-initiation state, EN activity is required ^30,31^, presumably to generate capped primers. Correspondingly, in this state, we observe that the EN is not autoinhibited, although how it might act in collaboration with the putative CBD is unclear. Whilst these biochemical experiments support the structure-based hypothesis that 5’ end only or full promoter binding activates the EN due to its displacement by the pendant and α-bundle domains, the observed EN activity is very weak (Supplementary Fig. 14), consistent with the barely detectable *in vitro* activity of the isolated EN domain ^20,21,31^. This suggests that some other L protein configuration or possibly a host factor may be required to fully activate the EN inside infected cells. Similarly, interaction with a currently unknown host cap-binding protein may be required to present capped RNAs to the L protein. Identification of how arenavirus L proteins access host capped RNAs is a key requirement for further understanding of the mechanism of transcription.

## Methods

### Expression and purification of LASV L protein

The L gene of LASV Bantou 289 (accession no. MK044799) containing a StrepII-tag at an internal position (after residue 407, 407strep) or a StrepII-His tandem tag at the C terminus (Cstrep) was cloned into an altered pFastBacHT B vector as described previously ^7^. If indicated, mutations were introduced by mutagenic PCR before cloning. Using DH10EMBacY E. coli cells ^42,43^, recombinant baculoviruses were produced and subsequently used for protein expression with the protocol being identical as described previously ^7^. The harvested Hi5 insect cells were resuspended in Buffer A (50 mM HEPES(NaOH) pH 7.0, 1 M NaCl, 10% (w/v) Glycerol and 2 mM dithiothreitol), supplemented with 0.05% (v/v) Tween20 and protease inhibitors (Roche, cOmplete mini), lysed by sonication and centrifugated two times (20,000 x g for 30 min at 4°C). Soluble protein was loaded on Strep-TactinXT beads (IBA) and eluted with 50 mM Biotin (Applichem) in Buffer B (50 mM HEPES(NaOH) pH 7.0, 500 mM NaCl, 10% (w/v) Glycerol and 2 mM dithiothreitol). L protein-containing fractions were pooled and diluted 1:1 with buffer C (20 mM HEPES(NaOH) pH 7.0) before loading on a heparin column (HiTrap Heparin HP, GE Healthcare). Proteins were eluted with Buffer A and concentrated using centrifugal filter units (Amicon Ultra, 30 kDa MWCO). The proteins were subsequently used for biochemical assays and structural studies. For endonuclease assays, the L proteins were further purified by size-exclusion chromatography (Superose 6, GE Healthcare) in buffer B. Pure L proteins were concentrated as described above, flash frozen and stored at −80°C.

### *In vitro* LASV-L complex reconstitution for cryo-EM

#### APO-structures

The LASV L-Cstrep protein was first injected onto a Superdex 200 Increase 3.2/300 column (GE Healthcare) equilibrated in 40 mM HEPES pH 7.4 (4 °C), 500 mM NaCl, 10 mM MgCl_2_ and 1 mM TCEP. 50 μl fractions were collected and the protein was eluted at 2 μM concentration. Protein was diluted to 0.7 μM and aliquots of 3 μl were applied to Quantifoil R1.2/1.3 Au 300 mesh grids, immediately blotted for 2 s and plunged into liquid ethane using an FEI Vitrobot IV (4 °C, 100% humidity).

#### PROMOTER-DUPLEX and MID-LINK structures

The LASV L-Cstrep protein was first injected onto a Superose 6 Increase 3.2/300 column (GE Healthcare) equilibrated at 4°C in 40 mM HEPES pH 7.4, 400 mM NaCl, 10 mM MgCl_2_ and 1 mM TCEP. 50 μl fractions were collected and the protein was eluted at 2 μM concentration. Protein was diluted to ~0.9 μM and mixed with 1.3-fold molar excess of truncated promoter vRNAs (5’ nts 10-19, 3’ nts 1-19) (Supplementary Table 2) for 10 minutes at 4 °C. Aliquots of 3 μl were applied to Quantifoil R2/2 Au 300 mesh grids, immediately blotted for 2 s and plunge frozen into liquid ethane using an FEI Vitrobot IV (4 °C, 100% humidity).

#### 3’END structures

The LASV L-Cstrep protein was first injected onto a Superose 6 Increase 3.2/300 column (GE Healthcare) equilibrated at 4°C in 40 mM HEPES pH 7.4, 250 mM NaCl, 10 mM MgCl_2_ and 1 mM TCEP. 50 μl fractions were collected and the protein was eluted at 2 μM concentration. Protein was diluted to ~1.8 μM and mixed with 3-fold molar excess of 3’ (1-16) vRNA (Supplementary Table 2) for 10 minutes at 4 °C. Aliquots of 3 μl were applied to Quantifoil R2/2 Au 300 mesh grids, immediately blotted for 2 s and plunge frozen into liquid ethane using an FEI Vitrobot IV (4 °C, 100% humidity).

#### PRE-INITIATION structure

The LASV L-Cstrep protein with a concentration of 1.4 µM in assay buffer (100 mM HEPES(NaOH) pH 7.0, 50 mM NaCl, 50 mM KCl, 2 mM MnCl_2_ and 2 mM dithiothreitol) was mixed sequentially with single stranded 5’ (0-19) vRNA and single stranded 3’ (1-19) vRNA in 1.2-fold and primer St1 in 7.1-fold molar excess (all RNAs are listed in Supplementary Table 2). After 45 min incubation on ice, the reaction was started by addition of NTPs (0.25 mM GTP/ATP). After incubation at 30°C for 2 h, 3 µL of the reaction was applied to glow-discharged Quantifoil R 2/1 Au G200F4 grids, immediately blotted for 2 s using an FEI Vitrobot Mk IV (4°C, 100% humidity, blotting force –10) and plunge frozen in liquid ethane/propane cooled to liquid nitrogen temperature.

#### ELONGATION structure

The LASV L-Cstrep protein with a concentration of 3 µM in assay buffer (100 mM HEPES(NaOH) pH 7.0, 50 mM NaCl, 50 mM KCl, 2 mM MnCl_2_ and 2 mM dithiothreitol) was mixed sequentially with single stranded 5’ (0-19) vRNA and single stranded 3’ (1-19) vRNA in 1.7-fold and primer C8 in 3.3-fold molar excess (all RNAs are listed in Supplementary Table 2). After 45 min incubation on ice, the reaction was started by addition of NTPs (0.25 mM GTP/ATP/UMPNPP and 0.125 mM CTP). After incubation at 30°C for 2 h, 3 µL of the reaction was applied to glow-discharged Quantifoil R 2/1 Au G200F4 grids, immediately blotted for 2 s using an FEI Vitrobot Mk IV (4°C, 100% humidity, blotting force –10) and plunge frozen in liquid ethane/propane cooled to liquid nitrogen temperature.

### Electron microscopy

#### APO-, DISTAL-PROMOTER, MID-LINK, PRE-INITIATION and ELONGATION structures

The grids were loaded into an FEI Tecnai Krios electron microscope at the Centre for Structural Systems Biology (CSSB) Cryo-EM facility, operated at an accelerating voltage of 300 kV and equipped with K3 direct electron counting camera (Gatan) positioned after a GIF BioQuantum energy filter (Gatan). Cryo-EM data were acquired using EPU software (FEI) at a nominal magnification of x105,000, with a pixel size of 0.85 or 0.87 Å per pixel. Movies of a total fluence of ~50 electrons per Å^2^ were collected at ~1 e-/Å^2^ per frame. A total number of 15,488 (APO-); 13,462 (DISTAL-PROMOTER and MID-LINK); 13,204 (PRE-INITIATION); 10,368 (ELONGATION) movies were acquired at a defocus range from −0.4 to −3.1 μm (Supplementary Table 1).

#### 3’ END-structures

The grids were loaded into an FEI Tecnai Krios electron microscope at European Synchrotron Radiation Facility (ESRF) beamline CM01 ^44^, operated at an accelerating voltage of 300 kV and equipped with K2 Summit direct electron counting camera (Gatan) positioned after a GIF Quantum energy filter (Gatan). Cryo-EM data were acquired using EPU software (FEI) at a nominal magnification of x165,000, with a pixel size of 0.827 Å per pixel. Movies of a total fluence of ~50 electrons per Å^2^ were collected at ~1 e-/Å^2^ per frame. A total number of 6,616 movies were acquired at a defocus range from −0.3 to −2.8 μm (Supplementary Table 1)

#### Cryo-EM image processing

All movie frames were aligned and dose-weighted using MotionCor2 program (Supplementary Cryo-EM processing overview file EM-1, -3, -5, -7, -9). Thon rings from summed power spectra of every 4e-/Å^2^ were used for contrast-transfer function parameter calculation with CTFFIND 4.1 ^45^. Particles were selected with WARP ^46^. The further 2D and 3D cryo-EM image processing was performed in RELION 3.1 ^47^. First, particles were iteratively subjected to two rounds of 2D-classification (Supplementary Cryo-EM processing overview file EM-1, -3, -5, -7, -9) at 2x binned pixel size. Particles in classes with poor structural features were removed.

#### 3D analysis of the APO-structures

Two times binned particles (1,491 k) were subjected to two rounds of 3D classifications with image alignment (Supplementary Cryo-EM processing overview file EM-2). The first round of 3D classification was restricted to ten classes and performed using 60 Å low-pass filtered initial model constructed from 16 most populated 2Ds class averages (Supplementary Cryo-EM processing overview file EM-2). Particles in classes with poor structural features were removed. The second classification (into ten classes) was done during two rounds of 25 iterations each, using regularization parameter T = 4. In the second round, local angular searches were performed at 3.5 ° to clearly separate structural species. Three major 3D species were identified: the bare CORE-like, the ENDO-like and RIBBON-like.

In the first branch of 3D classification, the focus was on the structure of the CORE of the LASV-L protein. All identified species were pooled together (428.8 k particles) and CORE-focused global 3D refinement was performed. Another round of CORE-focused 3D classification was performed based on the global refinement with local angular searches at 0.9° to clearly separate structural species. The most defined class (67.4 k particles) was further CORE-focused 3D auto-refined and iteratively aberration-corrected. For Bayesian-polishing only the first 23 frames were used.

In the second branch of 3D classification, the focus was on the structure of the ENDO-like species. The ENDO-like species from CSSB DATA 1 and DATA 2 were pooled together (209.4 k particles) and globally 3D auto-refined. A global 3D classification was performed based on the global refinement with local angular searches at 0.9 °. The most defined class (74 k particles) was further core-focused 3D auto-refined as described for the CORE-like specie.

In the third branch of 3D classification, the focus was on the structure of the RIBBON-like species (35.4 k particles). The RIBBON structure was obtained in a similar way as the ENDO one.

#### 3D analysis of the 3’ END-structures

Two times binned particles (654.8 k) were subjected to global 3D auto-refinement with 60 Å low-pass filtered APO-ENDO structure as initial model. In the first branch of 3D classification, the focus was on the structure of the 3’ end binding site. A specific mask containing the 3’ vRNA secondary binding site and pyramid domain area was created (yellow, Supplementary Cryo-EM processing overview file EM-4) and focused 3D classification without angular assignment was performed, using regularization parameter T = 4. The most defined class (194 k particles) was globally 3D auto-refined, iteratively aberration-corrected and Bayesian-polished (only the first 23 frames were used). The resulting global refinement was then subjected to core-focused 3D classification, with angular assignment using regularization parameter T = 8. The most defined class (84.5 k particles) was CORE-focused 3D auto-refined.

In the second branch of 3D classification, the focus was on the structure of the ENDO-like species. A specific mask containing the ENDO area was created (cyan, Supplementary Cryo-EM processing overview file EM-4) and ENDO-focused 3D classification without angular assignment was performed, using regularization parameter T = 4. The most defined class (159 k particles) was globally 3D auto-refined, iteratively aberration-corrected and Bayesian-polished (only the first 26 frames were used). The resulting global refinement was then subjected to ENDO-focused (cyan mask, Supplementary Cryo-EM processing overview file EM-4) 3D classification, without angular assignment using regularization parameter T = 12. The most defined class (40.2 k particles) was globally 3D auto-refined.

#### 3D analysis of the DISTAL-PROMOTER and MID-LINK structures

Two times binned particles (2,081 k) were subjected to two rounds of three-dimensional classifications with image alignment (Supplementary Cryo-EM processing overview file EM-8). The first round of 3D classification was restricted to twelve classes and performed using 60 Å low-pass filtered APO-ENDO structure as initial model. Particles in classes with poor structural features were removed. The second classification (into ten classes) was done during two rounds of 25 iterations each, using regularization parameter T = 4. In the second round, local angular searches were performed at 3.5° to clearly separate structural species. Three major 3D species were identified: DISTAL-PROMOTER-like, the MID-LINK-like and ENDO-like. The ENDO-like particles were processed together with CSSB DATA 1 (Supplementary Cryo-EM processing overview file EM-2).

The DISTAL-PROMOTER-like and MID-LINK-like species were pooled (65 k particles) and globally 3D auto-refined. In the first branch of 3D classification, the focus was on the structure of the DISTAL-PROMOTER-like specie. A specific mask containing the DISTAL-PROMOTER area was created (pink, Supplementary Cryo-EM processing overview file EM-8) and DISTAL-PROMOTER-focused 3D classification without angular assignment was performed, using regularization parameter T = 4. The most defined class (23.7 k particles) was further DISTAL-PROMOTER-CORE-focused (purple mask, Supplementary Cryo-EM processing overview file EM-8) 3D auto-refined as described for the CORE-like specie.

In the second branch of 3D classification, the focus was on the structure of the MID-LINK-like specie. A specific mask containing the MID-LINK area was created (orange, Supplementary Cryo-EM processing overview file EM-8) and MID-LINK-focused 3D classification without angular assignment was performed, using regularization parameter T = 4. The most defined class (40.7 k particles) was further MID-LINK-CORE-focused (yellow mask, Supplementary Cryo-EM processing overview file EM-8) 3D auto-refined as described for the CORE-like species.

#### 3D analysis of the PRE-INITIATION structure

Particles (2,470 k) were subjected to two rounds of reference-free 2D classification. Particles in classes with secondary structure features were selected (1,016 k particles) and used for an *ab initio* volume reconstruction and then 3D refined using the latter *ab initio* 60 Å low-pass filtered volume reconstruction as initial model. The particles were astigmatism corrected with CTFrefine. The particles were then subjected to a 3D classification restricted to twelve classes using T = 4 and 7.5 ° sampling for 25 iterations and then 3.5 ° sampling for an additional 10 iterations. Classes with comparable structural features were combined (319 k particles), 3D refined, aberration-corrected and Bayesian-polished then 3D refined again. The refined particles were then subjected to further 3D classification without image alignment and particles from the most defined class (119 k particles) were used for final 3D auto-refinement (Supplementary Cryo-EM processing overview file EM-5 and EM-6).

#### 3D analysis of the ELONGATION structure

Particles (2,452k) were binned four times and subjected to one round of reference free 2D classification. Particles in classes with secondary structure features were selected (579 k particles) and subjected to 3D classification restricted to ten classes using T = 4 and 7.5 ° sampling for 25 iterations with the 60 Å low-pass filtered DISTAL-PROMOTER volume as a reference. Classes with comparable structural features were combined (122 k particles) and 3D refined to 3.5 Å resolution. The 20 Å low-pass filtered refined volume was then used as a reference for 3D classification of all extracted particles (2,452k) restricted to eight classes using T = 4 and 7.5 ° sampling for 35 iterations. The most defined class (426 k particles) was selected for 3D refinement and then subjected to further 3D classification without image alignment restricted to six classes. The most defined class (79 k particles) was 3D refined, aberration-corrected and Bayesian-polished then finally 3D refined using SIDESPLITTER ^48^ (Supplementary Cryo-EM processing overview file EM-9 and EM-10).

#### Final steps

All final cryo-EM density maps were generated by the post-processing feature in RELION and sharpened or blurred into MTZ format using CCP-EM ^49^. The resolutions of the cryo-EM density maps were estimated at the 0.143 gold standard Fourier Shell Correlation (FSC) cut-off (Supplementary Cryo-EM processing overview file). A local resolution (Supplementary Cryo-EM processing overview file) was calculated using RELION and reference-based local amplitude scaling was performed by LocScale ^50^.

#### Model building

The APO-CORE structure was constructed *de novo* with iterative rounds of model-building with Coot ^51^ and real-space refinement with Phenix ^52^. Subsequent structures used this as a basis for further model extension. Secondary structure prediction using JPRED ^53^ based on multiple sequence alignment of both New World and Old World arenaviruses (Supplementary Alignment file) was particularly helpful in guiding model building. Considerable care was taken to cross-check between structures for consistency of sequence assignment and to ascertain connectivity. This also enabled better completion of models in lower resolution maps by transfer of structural elements that could be more accurately modelled in a higher resolution map. A homology model based on the MACV pendant domain (PDB: 6KLD), rebuilt to correct for sequence misalignments (using the original map, EMD-0707), was used to help build the LASV pendant domain. Unexpectedly, despite apparent sequence homologies, the LASV α-bundle 827-VVVNK…IIDQY-925 has a completely different arrangement of helices (topologically impossible to align in 3D) than that of the equivalent region in MACV L 820-VVIPK…QVALA-917 (Supplementary Fig. 6), both structures being confirmed by good quality maps. This might partly explain why in the LASV L structure previously published ^9^ (PDB:6KLC), based on a lower resolution 3.9 Å map, the α-bundle is built in the reverse direction.

Buried surface areas were determined using the Protein interfaces, surfaces and assemblies’ service PISA at the European Bioinformatics Institute (http://www.ebi.ac.uk/pdbe/prot_int/pistart.html) ^54^. An overview of the *Segment based Manders’ Overlap Coefficient* (SMOC) scores ^55^ for each of the structures is provided in Supplementary Fig. 23. Structure presentation was done using PyMOL (Schrödinger) and UCSF ChimeraX ^56^.

### Electrophoretic mobility shift assay

3’ RNA 1-10 nt (Supplementary Table 2) was chemically synthesized with a fluorophore at the 5’ end (5’ Cyanine3) (Biomers). Reactions containing 0-1 µM L protein and 0.2 µM labelled single-stranded RNA were set up in binding buffer (100 mM HEPES(NaOH), pH 7.0, 100 mM NaCl, 50 mM KCl, 2mM MnCl_2_, 2 mM dithiothreitol, 0.1 μg/μL Poly(C) RNA (Sigma), 0.1 µg/µL bovine serum albumine and 0.5 U/μl RNasin (Promega)) and incubated on ice for 30 min. RNA bound protein complexes were separated from unbound RNA by native gel electrophoresis at 4°C, using 5% polyacrylamide Tris-glycine gels. Fluorescence signals were detected in the gel with the VILBER LOURMAT FUSION SL4 using the Starlight Module with an excitation wavelength of 523 nm and a 609 nm emission filter.

### Endonuclease Assay

An RNA 16mer was chemically synthesized with either 5’ Cap (TriLink BioTechnologies) or 5’ Triphosphate (Chemgenes). For labeling pCp-Cy5 (Cytidine-5’-phosphate-3’-(6-aminohexyl)phosphate (Jena bioscience), was ligated to the 3’ end of the 16 nt RNA using T4 RNA ligase (Thermo Scientific). The resulting labelled 17 nt RNA substrates (Supplementary Table 2) were separated from excess pCp-Cy5 by denaturing PAGE (7 M urea, 25% acrylamide 0.5-fold Tris-borate-EDTA). The clearly blue colored product bands were excised from the gel. The gel pieces were grounded and the RNA was extracted two times with Tris-borate buffer. The pure labelled RNA was precipitated with 90% Ethanol from the supernatant after addition of Ammonium acetate (2.5 M), washed two times with 90% Ethanol and dissolved in DEPC treated H_2_O. Reactions containing 0.5 µM L protein were incubated, sequentially with 2.5 pmol of either single stranded 5’ promoter RNA, 3’ promoter RNA (Supplementary Table 2) or both, on ice for 15 min in 5 µL assay buffer (100 mM HEPES(NaOH) pH 7.0, 100 mM NaCl, 50 mM KCl, 2 mM MnCl_2_, 0.5 U/μl RNasin (Promega), 2 mM dithiothreitol, and 0.1 µg/µL bovine serum albumin). After addition of ~0.3 µM labelled RNA the mix was incubated at 37°C for 120 min. The reaction was stopped by adding an equivalent volume of RNA loading buffer (98% formamide, 18 mM EDTA, 0.025 mM SDS) and heating the samples at 95°C for 5 min. Products were separated by denaturing PAGE on 7 M Urea, 25% polyacrylamide Tris-borate-EDTA (0.5-fold) gels and 0.5-fold Tris-borate buffer. Fluorescence signals were detected in the gel with the VILBER LOURMAT FUSION SL4 using the Starlight Module with an excitation wavelength of 624 nm and a 695 nm emission filter.

### Polymerase Assay

#### Standard Polymerase Assay

If not indicated otherwise, 0.5 µM L protein was incubated sequentially with 1 µM of single stranded 5’ promoter RNA (Supplementary Table 2) and 1 µM single stranded 3’ promoter RNA (Supplementary Table 2) in assay buffer (100 mM HEPES(NaOH) pH 7.0, 50 mM NaCl, 50 mM KCl, 2 mM MnCl2, 0.5 U/μl RNasin (Promega), 2 mM dithiothreitol) on ice for 15 min. The reaction was started by addition of NTPs (0.25 mM UTP/ATP/CTP and 0.125 mM GTP supplemented with 166 nM, 5 µCi [α]^32^P-GTP) in a final reaction volume of 10 µL. After incubation at 30°C for 2 h the reaction was stopped by adding an equivalent volume of RNA loading buffer (98% formamide, 18 mM EDTA, 0.025 mM SDS, xylene cyanol and bromophenol blue) and heating the sample at 95°C for 5 min. Products were separated by native gel electrophoresis using 25% polyacrylamide 0.5-fold Tris-borate-EDTA gels and 0.5-fold Tris-borate running buffer. Signals were visualized by phosphor screen autoradiography using a Typhoon scanner (GE Healthcare).

#### Primer-dependent Polymerase Assay

Primer GCG, C1, St1 and C8 (Supplementary Table 2) were chemically synthesized with 5’-hydroxy ends (Biomers), primer C8ppp (Supplementary Table 2) with 5’ Triphosphate modification (Chemgenes). An N^7^-MeGppp (Cap0) was introduced at the 5’ terminus of C8ppp using the ScriptCap m7G Capping System (CELLSCRIPT) with 1 nmol C8ppp oligo using the manufacturers standard protocol. After addition of Ammonium acetate (2.5 M) the capped RNA was precipitated with Ethanol (90%), washed two times with Ethanol (90%), dried and dissolved in DEPC treated H_2_O. For primer-dependent reactions, 10 µM of the respective primer was added to LASV L bound to promoter RNA and the mix was again incubated on ice for 15 min. The reaction was started by addition of NTPs (0.25 mM UTP/ATP/CTP and 0.125 mM GTP supplemented with 166 nM, 5 µCi [α]^32^P-GTP) in a final reaction volume of 10 µL.

#### LASV mini-replicon system

The experiments were performed in the context of the T7 RNA polymerase-based LASV mini-replicon system essentially as described previously ^15,24,36^. L genes were amplified using mutagenic PCR from a pCITE2a-L template to either yield wild-type or mutated L gene expression cassettes. L gene PCR products were further gel purified when additional unspecific bands were visible in agarose gels and quantified spectrophotometrically. BSR-T7/5 cells stably expressing T7 RNA polymerase ^57^ were transfected per well of a 24-well plate with 250 ng of minigenome PCR product expressing Renilla luciferase (Ren-Luc), 250 ng of L gene PCR product, 250 ng of pCITE-NP expressing NP, and 10 ng of pCITE-FF-luc expressing firefly luciferase as an internal transfection control. At 24 h post transfection, either total cellular RNA was extracted for Northern blotting using an RNeasy minikit (Qiagen) or cells were lysed in 100 µL of passive lysis buffer (Promega) per well, and firefly luciferase and Ren-Luc activity were quantified using the dual-luciferase reporter assay system (Promega). Ren-Luc levels were corrected with the firefly luciferase levels (resulting in standardized relative light units [sRLU]) to compensate for differences in transfection efficiency or cell density.

For Northern blot analysis, 750 - 2000 ng of total cellular RNA was separated in a 1.5% agarose-formaldehyde gel and transferred onto a Roti®-Nylon plus membrane (pore size 0.45 μm, Carl Roth). After UV crosslinking and methylene blue staining to visualize 28S rRNA the blots were hybridized with a ^32^P-labelled riboprobe targeting the Ren-Luc gene. Transcripts of Ren-Luc genes and complementary replication intermediate RNA of the minigenome were visualized by autoradiography using an FLA-7000 phosphorimager (Fujifilm). To verify expression of L protein mutants in BSR-T7/5 cells, the cells were transfected with 500 ng of PCR product expressing C-terminally 3xFLAG-tagged L protein mutants per well in a 24-well plate. Cells were additionally infected with modified vaccinia virus Ankara expressing T7 RNA polymerase (MVA-T7) ^58^ to boost the expression levels and thus facilitate detection by immunoblotting. After cell lysis and electrophoretic separation in a 3-8% Tris-acetate polyacrylamide gel, proteins were transferred to a nitrocellulose membrane (GE Healthcare). FLAG-tagged L protein mutants were detected using peroxidase-conjugated anti-FLAG M2 antibody (1:10,000) (A8592; Sigma-Aldrich) and bands were visualized by chemiluminescence using Super Signal West Femto substrate (Thermo Scientific) and a FUSION SL image acquisition system (Vilber Lourmat).

## Supporting information

Supplementary Alignment file

Supplementary Cryo-EM processing overview file

## Acknowledgements

We acknowledge the European Synchrotron Radiation Facility for provision of beam time on CM01 and we would like to thank Daouda A.K. Traore for assistance. Furthermore, we want to thank Sophia Reindl and Nadja Hüttmann for advice and technical support in the early phases of this project; Michael Hons, Wojtek Galej and Erika Pellegrini for access to the Glacios at EMBL Grenoble; Carolin Seuring and Cornelia Cazey for access to Cryo-EM facility at CSSB; Aymeric Peuch and Wolfgang Lugmayr for help with using the joint EMBL-IBS and the CSSB partition on the DESY computer cluster. We acknowledge funding of this project by the Leibniz Association’s Leibniz competition programme (grant K72/2017 to S.G., K.G. and S.C.), the Federal Ministry of Education and Research of Germany (grant 01KI2019 to M.R.), the Wilhelm und Maria Kirmser-Stiftung, the Alexander von Humboldt foundation (individual fellowship to E.Q.) as well as the EMBL Interdisciplinary Postdocs (EI3POD) initiative co-funded by Marie Skłodowska-Curie (grant 664726 to T.K.). Part of this work was performed at the Cryo-EM Facility at CSSB, supported by the UHH and DFG (grants INST 152/772-1 and 774-1).

This is a preprint of an article published in *Nature Communications*. The final authenticated version is available online at: https://doi.org/10.1038/s41467-021-27305.

## Funding

Leibniz Association, Leibniz competition programme [K72/2017]; Federal Ministry of Education and Research of Germany [01KI2019], Wilhelm und Maria Kirmser-Stiftung; Part of this work was performed at the Cryo-EM Facility at CSSB, supported by the UHH and DFG [INST 152/772-1 and 774-1]; Individual fellowship from the Alexander von Humboldt foundation (to E.Q.); T.K. holds a fellowship from the EMBL Interdisciplinary Postdocs (EI3POD) initiative co-funded by Marie Skłodowska-Curie [664726].

## Data availability

Data are available from the corresponding authors. Coordinates and structure factors or maps have been deposited in the wwwPDB or EMDB:

Apo-structure of Lassa virus L protein (well-resolved polymerase core) [APO-CORE] EMD-12807, PDB ID 7OCH

Apo-structure of Lassa virus L protein (well-resolved endonuclease) [APO-ENDO] EMD-12860, PDB ID 7OE3

Apo-structure of Lassa virus L protein (well-resolved α-ribbon) [APO-RIBBON] EMD-12953, PDB ID 7OE7

Lassa virus L protein bound to 3’ promoter RNA (well-resolved polymerase core and 3’ RNA secondary binding site) [3END-CORE] EMD-12862, PDB ID 7OEA

Lassa virus L protein bound to 3’ promoter RNA (well-resolved endonuclease) [3END-ENDO] EMD-12863, PDB ID 7OEB

Lassa virus L protein in a pre-initiation conformation [PRE-INITIATION] EMD-12955, PDB ID 7OJL

Lassa virus L protein with endonuclease and C-terminal domains in close proximity [MID-LINK] EMD-12861, PDB ID 7OJJ

Lassa virus L protein bound to the distal promoter duplex [DISTAL-PROMOTER] EMD-12954, PDB ID 7OJK

Lassa virus L protein in an elongation conformation [ELONGATION] EMD-12956, PDB ID 7OJN

Source data are provided with this paper.

## Author contributions

T.K., D.V., E.Q., S.G., K.G., M.R. and S.C. conceived and supervised the project. C.B. and M.M. carried out cloning. D.V. and C.B. expressed and purified the proteins. T.K., D.V. and S.T. prepared the cryo-EM grids. T.K., S.T. and E.Q. collected and processed the cryo-EM data, D.V., H.W., M.R. and S.C. built and validated the models; D.V. and H.W. performed the *in vitro* experiments, M.M. performed the cell-based mini-genome experiments, T.K., D.V., H.W. and M.R. compiled the figures, M.R. and S.C. wrote the manuscript with input from all co-authors.

## Competing interests statement

The authors declare no competing interests.

## Supplementary tables and figure captions

**Supplementary Table 1: Cryo-EM data collection, refinement and validation statistics.** This table provides the statistics for the data collection, refinement and structure validation of the cryo-EM structures. Refinement statistics were generated using Phenix comprehensive validation software.

**Supplementary Table 2: Synthetic RNA oligos used in the assays.** The table lists the RNA oligonucleotides that were used in the described assays. The specific sequence, length in nucleotides and the identifier used to label the RNA in the experimental descriptions are given. Underlined nucleotides in the 5’ (nts 0-19) and 3’ (nts 1-19) indicate mismatches between the two otherwise complementary RNA strands. All RNAs were synthesized by Biomers, TriLink BioTechnologies or Chemgenes.

**Supplementary Figure 1.**
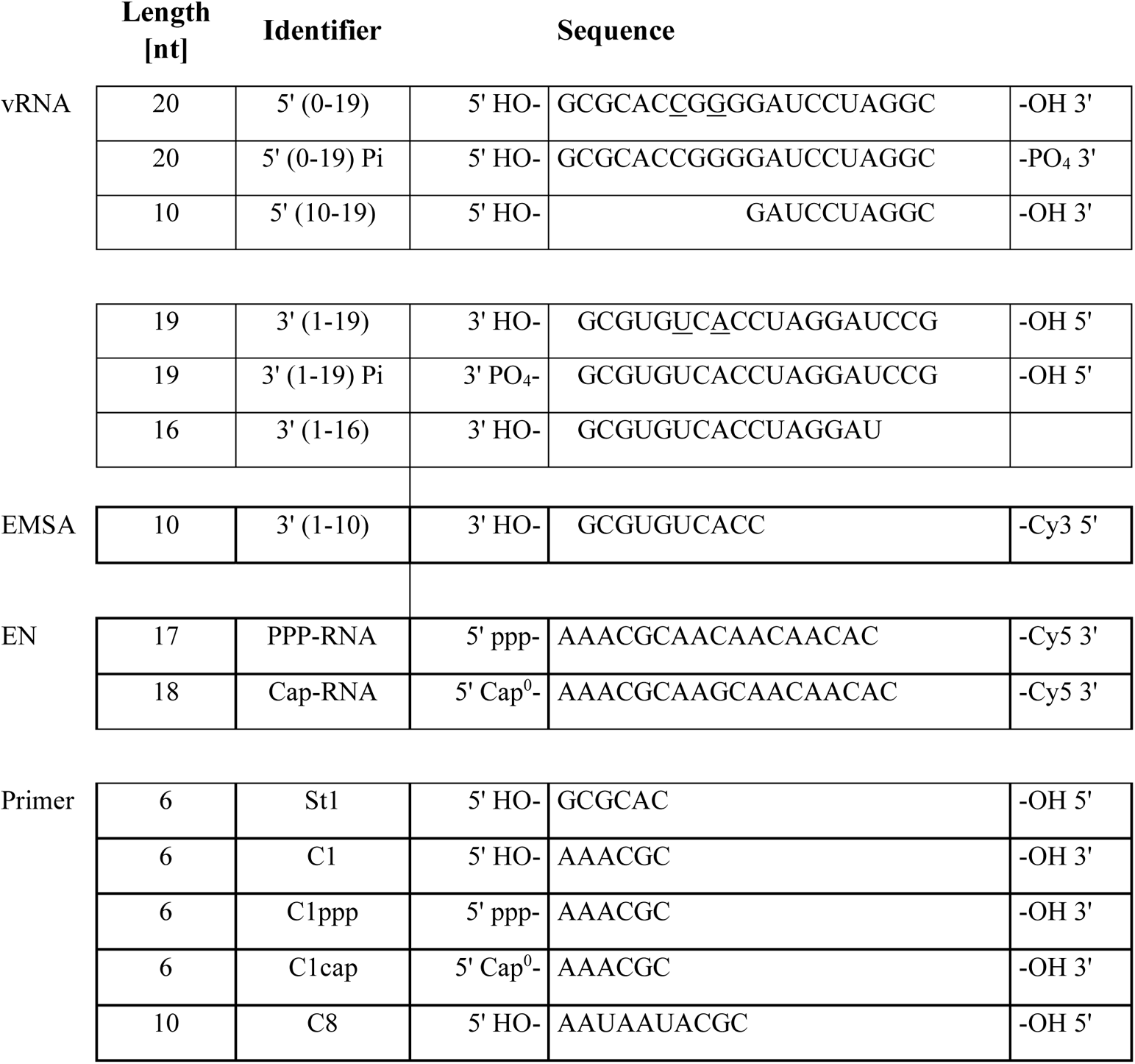
PRE-INITIATION and DISTAL-PROMOTER structures. **a.** An overview of the PRE-INITIATION structure is shown with the promoter RNA highlighted in orange (3’ vRNA) and pink (5’ vRNA). **b.** LASV mini-replicon data for L proteins with mutations in the 5’ hook RNA binding site presenting luciferase reporter activity (in standardized relative light units relative to the wild-type L protein (WT), mean average of 3 biological replicates), Northern blotting results with signals for antigenomic viral RNA (AG), viral mRNA (mRNA) and 28S ribosomal RNA (28S) as a loading control, and Western blot detection of FLAG-tagged L proteins (L) to demonstrate general expressibility of the mutants. **c.** The DISTAL-PROMOTER structure is shown with the promoter RNA highlighted in orange (3’ vRNA) and pink (5’ vRNA). **d.** The conformation of the EN domain (orange) and EN linker (green) is highlighted in a ribbon diagram presentation of the DISTAL-PROMOTER structure. Note that the EN is not resolved in the PRE-INITIATION structure.

**Supplementary Figure 2. MID-LINK structure. a.** The cryo-EM map of the MID-LINK structure is shown from two perspectives as indicated. Domains are colored and labelled according to Fig. 3. Additional low-resolution density for the CBD-like domain is indicated by a dashed green circle. **b.** Comparison of only the mid-link (violet) and 627-like (pink) domains of MID-LINK and ELONGATION structures as well as the CASV crystal structure of the L protein C-terminal region (PDB:5MUS).

**Supplementary Figure 3. Comparison of related polymerase structures.** The structures of influenza virus (PDB: 6T0V, elongation state), LACV (PDB: 6Z8K, elongation state), LASV (ELONGATION), and Severe fever with thrombocytopenia syndrome virus (SFTS virus, PDB:7ALP, apo-form) are shown side by side. Structures were divided according to influenza virus PA, PB1 and PB2 subunits and corresponding elements are shown in the same color.

**Supplementary Figure 4. Structural Zinc-binding site in the pyramid domain. a.** Close-up of the Zinc-binding site of APO-CORE structure with the protein chain shown as sticks (teal) and the Zinc ion as sphere (cyan) together with the experimental map shown as a grey mesh. Co-ordination is indicated by red doted lines. **b.** LASV mini-replicon data for L proteins with mutations in and around the Zinc-binding site presenting luciferase reporter activity (in standardized relative light units relative to the wild-type L protein (WT), mean average of 4 biological replicates), Northern blotting results with signals for antigenomic viral RNA (AG), viral mRNA (mRNA) and 28S ribosomal RNA (28S) as a loading control, and Western blot detection of FLAG-tagged L proteins (L) to demonstrate general expressibility of the mutants. **c.** Overview of the L protein of MACV (PDB:6KLE) as a ribbon diagram. Corresponding domains are colored according to Fig. 3 and labelled. **d.** Close-up of the MACV Zinc-binding site. Colors according to (c).

**Supplementary Figure 5. Polymerase active site in ELONGATION and APO-CORE structures.** Superimposition of the RdRp active sites of the ELONGATION (colored) and APO-CORE (grey) structures as ribbon diagram. Conserved RdRp motifs are shown in distinct colors and with labels. Product RNA (grey) and non-hydrolysable UTP (UMPNPP, yellow) of the ELONGATION structure are shown as sticks.

**Supplementary Figure 6. Pendant and α-bundle domains.** Comparison of MACV L (PDB: 6KLE), LASV DISTAL-PROMOTER and LASV PRE-INITIATION structures as ribbon diagrams with the pendant (pink) and α-bundle/ribbon (dark blue) domains shown in color and indicated by name.

**Supplementary Figure 7. Conformation of the C-terminal domains. a.** Colored cryo-EM maps of MID-LINK and ELONGATION structures from three perspectives highlighting the position of the C-terminal mid-link (violet), CBD-like (palegreen) and 627-like (pink) domains. Other domains are colored as in Fig. 3. Possible rotation of the C-terminal domains in respect to the core is indicated. **b.** Rotation of the CBD-like domain (palegreen) relative to the mid-link (violet) domain is indicated. Mid-link and 627-like domains of MID-LINK and ELONGATION have been superposed and CBD-like domain of the ELONGATION structure is shown as ribbon diagram. Density in the cryo-EM map for the CBD-like domain in the MID-LINK structure is visible in grey. **c.** Similarity of the conformation of the CBD-like domain (palegreen) in MACV L protein (light grey, PDB:6KLH) and the LASV MID-LINK structure (dark grey), where the position of the CBD-like domain at low resolution is marked by a dashed circle. Mid-link, CBD-like and 627-like domains are shown in color (violet and pink for LASV, dark red, cyan and hotpink for MACV).

**Supplementary Figure 8. (Putative) cap-binding domains. a.** Comparison between different putative and functional CBDs: Influenza virus (PDB:2VQZ), SFTSV (PDB:6XYA), RVFV (PDB:6QHG), LACV (PDB:6Z8K), CASV (PDB:5MUS) and LASV (ELONGATION). Corresponding secondary structure elements of ribbon diagrams are displayed in the same colors. **b.** Side-by-side comparison between reptarenavirus (CASV) and mammarenavirus (LASV, ELONGATION structure) CBD-like domains. Insertions of the LASV domain compared to the CASV domain are colored in blue and red. Secondary structure elements are labelled. **c.** LASV mini-replicon data for L proteins with mutations of aromatic residues potentially involved in cap-binding presenting luciferase reporter activity (in standardized relative light units relative to the wild-type L protein (WT), mean average of 4 biological replicates), Northern blotting results with signals for antigenomic viral RNA (AG), viral mRNA (mRNA) and 28S ribosomal RNA (28S) as a loading control, and Western blot detection of FLAG-tagged L proteins (L) to demonstrate general expressibility of the mutants. **d.** Conservation of the C-terminal domain within arenaviruses according to the Supplementary Alignment file illustrated using the ConSurf web server ^59^. Color legend is provided.

**Supplementary Figure 9. Domain interactions of the C-terminal region. a.** The cryo-EM map of the ELONGATION structure is presented with colors of domains according to Fig. 3. In close-ups of different perspectives, the neighboring domains of the mid-link, 627-like and CBD-like domains are visible. **b.** Close-ups of the interaction between residues identified as selectively important for viral transcription ^24^ of the 627-like domain with the palm (red) and thumb-ring (yellow) domains. Important main and side chains are shown as sticks with respective labels.

**Supplementary Figure 10. Interactions of the EN domain. a.** Close-ups of the interactions of residue P109 in the 3END-CORE, MID-LINK and ELONGATION structures with important side chains shown as sticks, labelled respectively. **b.** LASV mini-replicon data for L proteins with mutations P109A and F1592A presenting luciferase reporter activity (in standardized relative light units relative to the wild-type L protein (WT), mean average of 7 biological replicates), Northern blotting results with signals for antigenomic viral RNA (AG), viral mRNA (mRNA) and 28S ribosomal RNA (28S) as a loading control, and Western blot detection of FLAG-tagged L proteins (L) to demonstrate general expressibility of the mutants. **c.** Ligplot diagram ^60^ of the interface between the inhibitory peptide and the EN domain in the 3END-CORE structure generated by PDBsum ^61^. **d.** Close-up of the EN active site in the ELONGATION structure showing residues E188, E51 and D89 co-ordinating the divalent metal ions. Structure is shown as a ribbon diagram, important residues as sticks and with respective labels.

**Supplementary Figure 11. The EN and the inhibitory peptide. a.** Close-ups of the inhibitory peptide (cyan) and interacting residues in the 3END-CORE, ELONGATION and MID-LINK structures. EN is shown in orange, EN linker in green. **b.** LASV mini-replicon data for L proteins with mutations of residues involved in the interface between EN (orange background) and the inhibitory peptide (cyan background) (comp. Supplementary Fig. 10) presenting luciferase reporter activity (in standardized relative light units relative to the wild-type L protein (WT), mean average of 4 biological replicates), Northern blotting results with signals for antigenomic viral RNA (AG), viral mRNA (mRNA) and 28S ribosomal RNA (28S) as a loading control, and Western blot detection of FLAG-tagged L proteins (L) to demonstrate general expressibility of the mutants. **c.** Superimposition of the 3END-CORE EN domain (orange) with the crystal structure of Lymphocytic choriomeningitis virus (LCMV) EN domain (PDB:5LTN, yellow) solved in complex with the specific inhibitor 2,4-dioxo-4-phenylbutanoic acid (DPBA, green) bound to the EN. DPBA and the inhibitory peptide (cyan) are occupying the same space, i.e. the EN active site. Divalent metal ions (blue) and DPBA are shown as spheres. **d.** Superimposition as in (c) but with the LCMV structure shown as surface electrostatics calculated by APBS electrostatics plugin within PyMOL (Schrödinger).

**Supplementary Figure 12. Mutations of the interface of EN and the inhibitory peptide in combination with EN active site mutation D89A.** LASV mini-replicon data for L proteins with double mutations of residues involved in the interface between EN (orange background) and the inhibitory peptide (cyan background) as well as EN active site mutation D89A presenting luciferase reporter activity (in standardized relative light units relative to the wild-type L protein (WT), mean average of 7 biological replicates), Northern blotting results with signals for antigenomic viral RNA (AG), viral mRNA (mRNA) and 28S ribosomal RNA (28S) as a loading control, and Western blot detection of FLAG-tagged L proteins (L) to demonstrate general expressibility of the mutants. As a control the phenotype of a selective transcription defect expected with a single D89A mutation is shown as well (right panels).

**Supplementary Figure 13. *In vitro* polymerase activity of selected L protein mutants.** The influence of the mutations Q114A in the EN, and Y1099A and E1102A in the inhibitory peptide on the polymerase activity of purified LASV L was tested *in vitro* and compared to the wild-type L protein (WT). The reactions were carried out with only the 3’ vRNA (nts 1-19) present or with the 3’ vRNA (nts 1-19) Pi and 5’ vRNA (nts 0-19) Pi under standard polymerase assay conditions (see Methods). Products were separated by denaturing gel electrophoresis and visualized by autoradiography.

**Supplementary Figure 14. *In vitro* endonuclease activity of selected L protein mutants.** The ability to degrade 3’ Cy5 labelled RNA with a 5’ Cap (m7GTP-RNA-Cy5) or a 5’ triphosphate (PPP-RNA-Cy5) was tested for wild-type LASV L (WT), the mutant E1102A in the inhibitory peptide and the EN mutant Q114A. 2,4-Dioxo-4-Phenylbutanoic Acid (DPBA), known to inhibit viral endonucleases and the endonuclease inactive mutant E102A served as negative controls. 500 nM of the respective LASV L was incubated with ~0.3 μM of the fluorescently labelled RNA substrates (Supplementary Table 2) in assay buffer at 37°C for 2h (see Methods). Reaction products were separated on a denaturing polyacrylamide gel and fluorescence signals were detected with a VILBER LOURMAT FUSION SL4 imaging system using the Starlight Module with an excitation wavelength of 624 nm and a 695 nm emission filter.

**Supplementary Figure 15. 3’ RNA bound to the secondary binding site.** Experimental map (grey mesh) of the 3’ RNA (orange sticks) and the water molecules (red mesh and spheres) in the 3END-CORE structure from two perspectives as indicated.

**Supplementary Figure 16. In vitro activity of the wild-type L protein vs. mutant Y1450A/R1452A. a.** The influence of the double-mutation Y1450A/R1452A of the 3’ vRNA secondary binding site on the polymerase activity of purified LASV L was tested *in vitro* and compared to the wild type (WT). The reactions were carried out under standard polymerase assay conditions (see Methods) either with (i) only the 3’ vRNA (nts 1-19) Pi or (ii) 5’ vRNA (nts 0-19) Pi present, (iii) together with both 3’ and 5’ vRNAs or (iv) in the presence of a 47 nt hairpin RNA containing the connected 3’ and 5’ promoter sequences. Products were separated by denaturing gel electrophoresis and visualized by autoradiography. **b.** SDS-PAGE analysis of the purified LASV L proteins.

**Supplementary Figure 17. 5’ vRNA hook. a.** Experimental map of the 5’ vRNA hook (grey mesh) around the RNA shown as sticks from two perspectives as indicated. **b.** Close-up of the interaction between the pyramid base and the G0 nucleotide of the 5’ vRNA. Hydrogen bonds are indicated by dotted lines. Important main and side chains are shown as sticks with respective labels. **c.** Comparison between the 5’ vRNA hook structures of LASV (nts 0-9, PRE-INITIATION, pink), LACV (nts 1-10, PDB: 6Z6G, blue), and influenza virus (nts 1-10, PDB: 6T0V, green) presented as sticks. Hydrogen bonds between bases are shown as dotted black lines and bases are labelled.

**Supplementary Figure 18. Primed RNA synthesis by LASV L protein. a.** The influence of the tri-nucleotide GCG and the C8 primer (Supplementary Table 2) on the polymerase activity of purified LASV L was tested *in vitro* compared to the unprimed (*de novo*) reaction. The reactions were carried out in presence of both 3’ vRNA (nts 1-19) Pi and 5’ vRNA (nts 0-19) Pi under standard polymerase assay conditions (see Methods). Products were separated by denaturing gel electrophoresis and visualized by autoradiography. **b.** For polymerase stalling, LASV L protein was incubated *in vitro* with C8 primer (Supplementary Table 2) and increasing amounts of the non-hydrolysable UTP analogue UMPNPP under primer dependent polymerase assay conditions (see Methods). The reaction was started by addition of NTPs (0.25 mM GTP/ ATP/ 0.125 mM CTP and 0.03-0.25 mM UMPNPP). Products were separated by denaturing gel electrophoresis and visualized by autoradiography. Possible products are shown on the right. Positions for potentially mis-incorporated nucleotides (N) or stalling by UMPNPP (U_NPP_) are indicated.

**Supplementary Figure 19. Template and product RNA in the ELONGATION structure. a.** Experimental map of the 3’ RNA template (grey mesh) around the RNA shown as orange sticks. **b.** Experimental map of the product RNA (grey mesh) around the RNA (black) and the non-hydrolysable UTP (UMPNPP, yellow) shown as sticks.

**Supplementary Figure 20. Active site comparison between ELONGATION and PRE-INITIATION structures. a.** Close-up of the RdRp active site of the ELONGATION (black) and PRE-INITIATION (grey) structures. Product RNA and non-hydrolysable UTP (UMPNPP) are shown as sticks. Motif F of the ELONGATION structure is highlighted in magenta. **b.** Close-up of the RdRp active site towards the product exit channel with the product RNA shown as grey ribbon. Thumb-ring and lid domains of the ELONGATION structure are shown in yellow and brown, respectively. PRE-INITIATION structure shown in grey. Corresponding α-helices 52, 53 and 59 are labelled. Residue T1583 stacking on the terminal nucleotide of the product RNA is shown as sticks.

**Supplementary Figure 21. Role of the 5’ RNA.** The influence of the 5’ vRNA (nts 0-19) and 5’ vRNA (nts 0-12, corresponding to the hook) on the polymerase activity of purified LASV L was tested *in vitro* for the unprimed (*de novo*) or primed reaction using the tri-nucleotide GCG or the C8 primer (Supplementary Table 2). The reactions were carried out under standard polymerase assay conditions (see Methods) either with (i) only the 3’ vRNA (nts 1-19) present, (ii) together with 5’ vRNA (nts 0-19) or (iii) 5’ vRNA (nts 0-12). Reactions with LASV L and C8, 5’ vRNA (nts 0-19) or 5’ vRNA (nts 0-12) only are provided as additional controls. Products were separated by denaturing gel electrophoresis and visualized by autoradiography.

**Supplementary Figure 22. *In vitro* polymerase activity of LASV L in presence of capped and uncapped primers.** The influence of the 5’ modification of the primer C1: 5’-OH, 5’-ppp and 5’-Cap0 (Supplementary Table 2) on the polymerase activity of purified LASV L was tested *in vitro* compared to the unprimed (*de novo*) reaction. The reactions were carried out in presence of both vRNAs (3’ vRNA (nts 1-19) Pi and 5’ vRNA (nts 0-19) Pi) under primer dependent polymerase assay conditions (see Methods). Products were separated by denaturing gel electrophoresis and visualized by autoradiography. The 1:10 dilution of the Cap0-C1 primer gives more intense product bands than the undiluted primer probably as there is remaining GTP from the capping reaction (even after precipitation), which is incorporated instead of radioactive GTP.

**Supplementary Figure 23. Comparison of LASV L Protein Structures by TEMPy SMOC Score.** Schematic representation of the domain structure of the LASV L protein as in Fig. 3 (top panel). For each LASV L protein structure, the SMOC scores per residue were calculated by TEMPy within CCP-EM ^49,62^ providing the respective resolution as described in Supplementary Table 1 and plotted revealing the quality of the local fit (bottom panels). The domain structure of LASV L protein as shown in the top panel has been added as background to each SMOC plot to highlight which domains are missing/present in each structure. Absolute numbers of SMOC scores are resolution-dependent, thus the y-axis differs between the panels.

**Supplementary Movie 1. The PRE-INITIATION structure.** This movie highlights the interactions between the L protein and the 3’ and 5’ promoter RNA. Hydrogen bonds and electrostatic interactions are shown as dotted black lines.

**Supplementary Movie 2. The ELONGATION structure.** This movie presents the L protein stalled in an early elongation state with a non-hydrolysable UTP (UMPNPP) in the RdRp active site. Hydrogen bonds and electrostatic interactions are shown as dotted black lines. The movie further shows the putative product and template exit channels highlighting an interaction of the thumb domain with the RNA potentially involved in strand separation of template and product RNA duplex.

**Supplementary Movie 3. Overview of the L protein structure.** This movie introduces the different domains of the LASV L protein (ELONGATION structure).

**Supplementary Cryo-EM processing overview file.** This file includes representative micrographs, 2D-class averages, Fourier shell correlation curves, local resolution distributions and summaries of the cryo-EM 3D classification and 3D refinement schemes for all cryo-EM data in this study.

**Supplementary Alignment file. Alignment of full-length L protein sequences of Old World and New World arenaviruses.** L protein sequences (virus abbreviations, GenBank IDs listed) were aligned using ClustalOmega (https://www.ebi.ac.uk/Tools/msa/clustalo/) ^63^ with manual adjustments. The alignment was presented by ESPript ^64^ with secondary structure information given for LASV Bantou 289 L protein (PDB 7OJN). Regions of the L protein are labelled and marked with different colors according to Fig. 3. Conserved RdRp motifs are labelled according to Fig. 4e. All numbers given refer to LASV Bantou 289 L protein sequence.

## Notes

https://www.rcsb.org/structure/7OJN

https://www.rcsb.org/structure/7OCH

https://www.rcsb.org/structure/7OE3

https://www.rcsb.org/structure/7OE7

https://www.rcsb.org/structure/7OEA

https://www.rcsb.org/structure/7OEB

https://www.rcsb.org/structure/7OJJ

https://www.rcsb.org/structure/7OJK

https://www.rcsb.org/structure/7OJL

### Competing Interest Statement

The authors have declared no competing interest.

### Summary of Updates

We added a sentence to the acknowledgements section referring to the final version of this manuscript, which was published in Nature Communications. The final authenticated version is available online at: https://doi.org/10.1038/s41467-021-27305

