## Supplementary Alignment file for "Conformational changes in Lassa virus L protein associated with promoter binding and RNA synthesis activity"

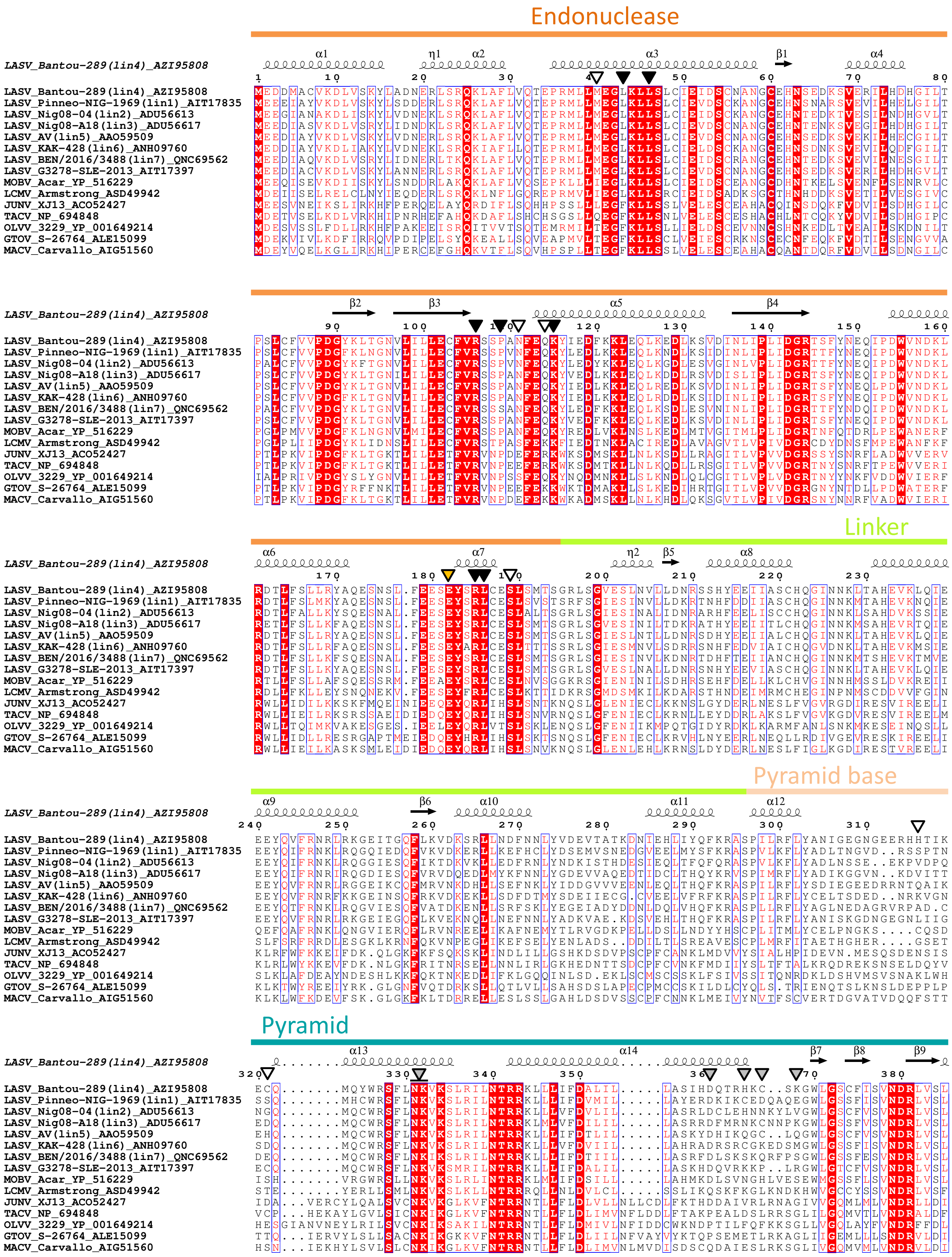


▼ – general defect in L protein activity upon mutation observed in Lassa virus mini-replicon system

▼ (yellow) – selective transcriptional defect of L protein upon mutation observed in Lassa virus mini-replicon system

▽ – no or weak effect on L protein activity upon mutation observed in Lassa virus mini-replicon system

▽ (grey) – intermediate effect on L protein activity upon mutation observed in Lassa virus mini-replicon system

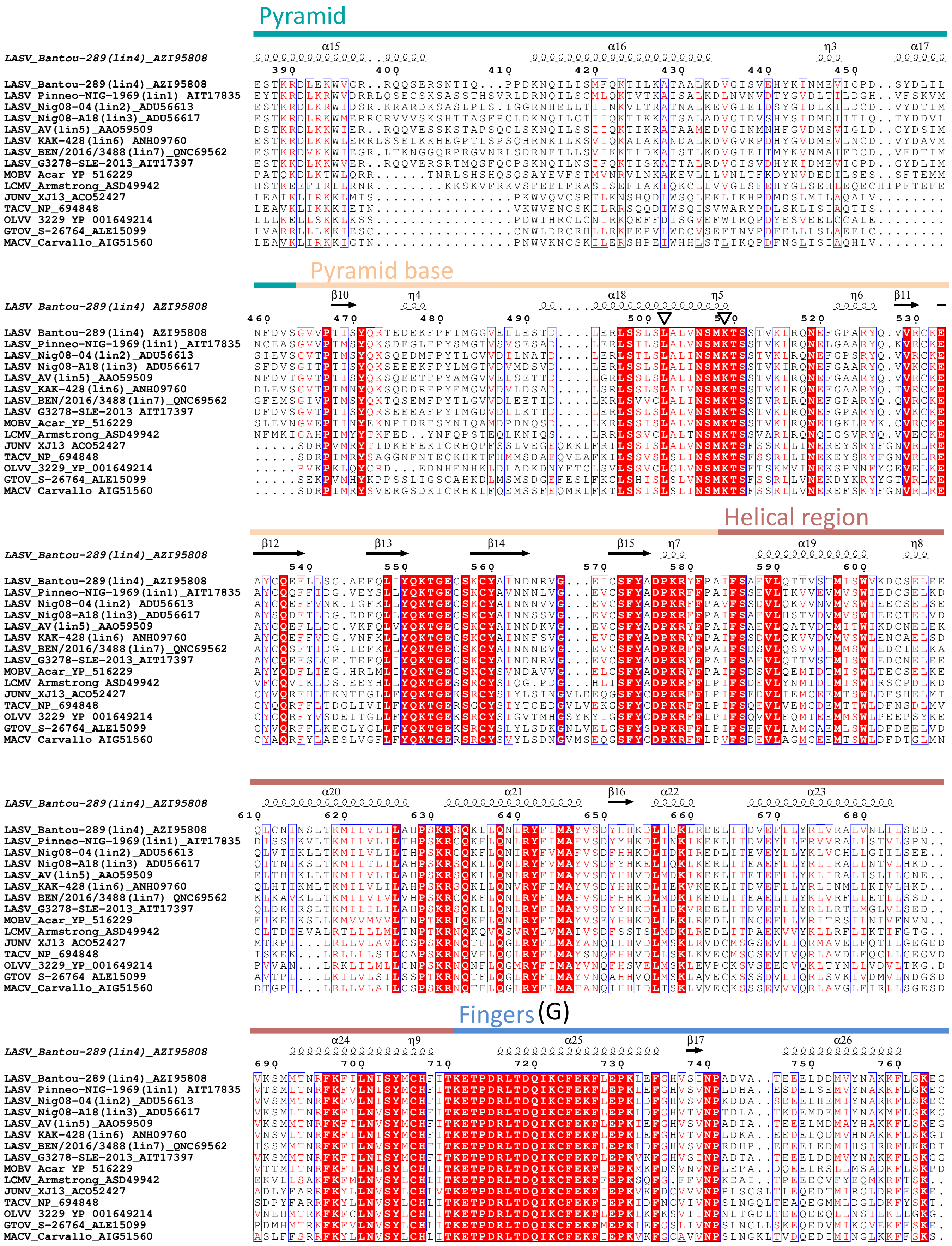


▼ – general defect in L protein activity upon mutation observed in Lassa virus mini-replicon system

▼ – selective transcriptional defect of L protein upon mutation observed in Lassa virus mini-replicon system

▽ – no or weak effect on L protein activity upon mutation observed in Lassa virus mini-replicon system

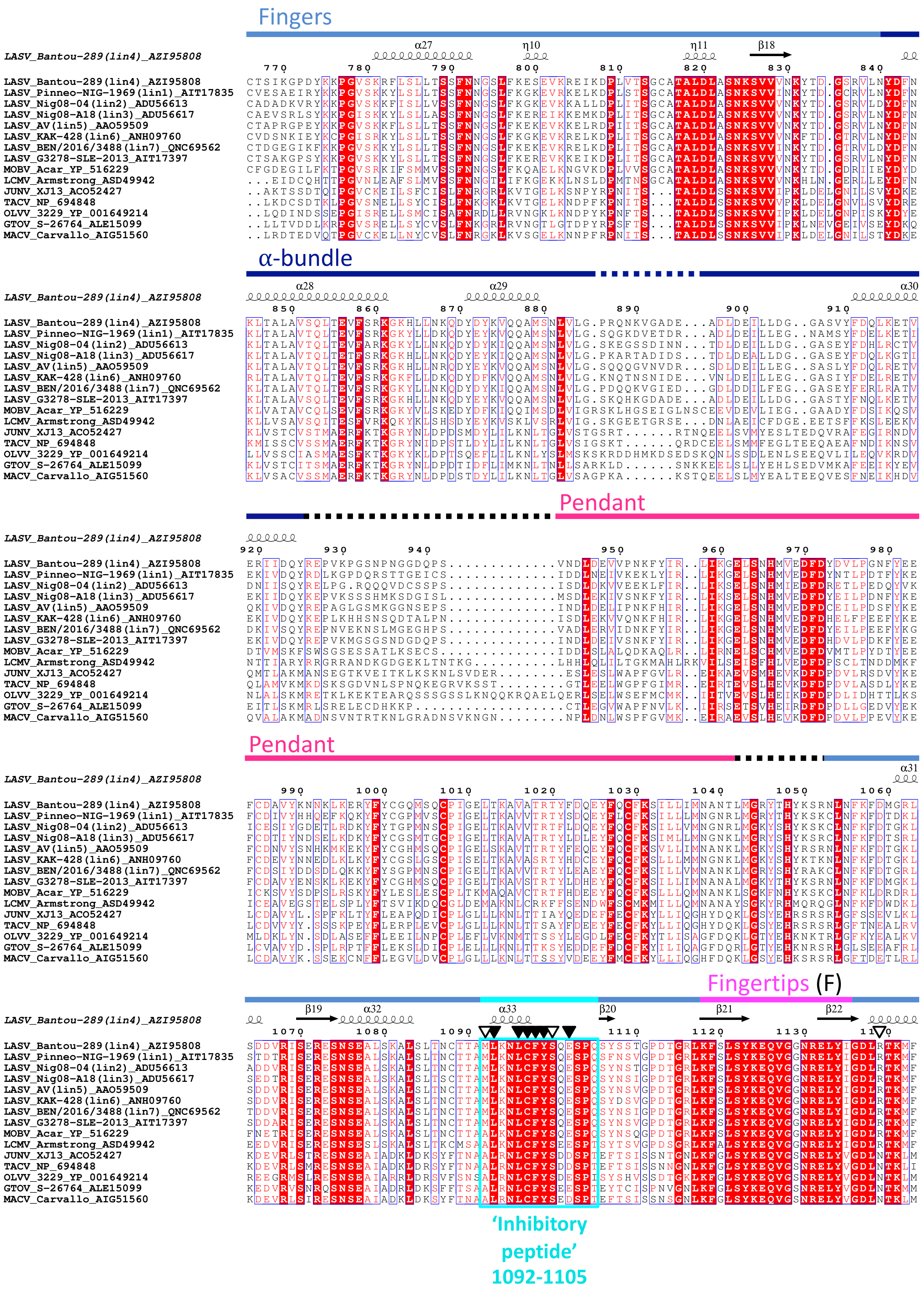


▼ – general defect in L protein activity upon mutation observed in Lassa virus mini-replicon system

▼ – selective transcriptional defect of L protein upon mutation observed in Lassa virus mini-replicon system

▽ – no or weak effect on L protein activity upon mutation observed in Lassa virus mini-replicon system

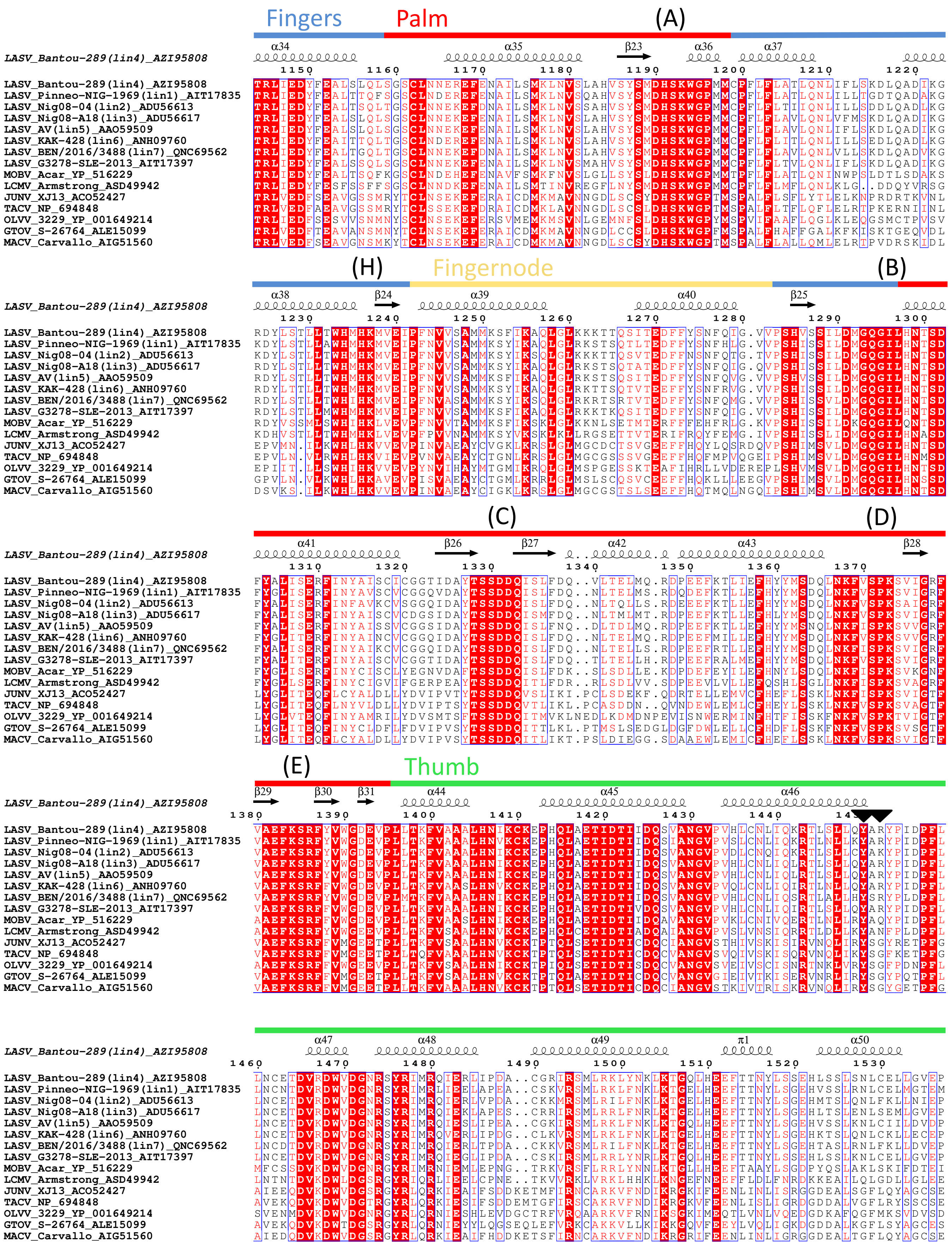


▼ – general defect in L protein activity upon mutation observed in Lassa virus mini-replicon system

▼ – selective transcriptional defect of L protein upon mutation observed in Lassa virus mini-replicon system

▽ – no or weak effect on L protein activity upon mutation observed in Lassa virus mini-replicon system

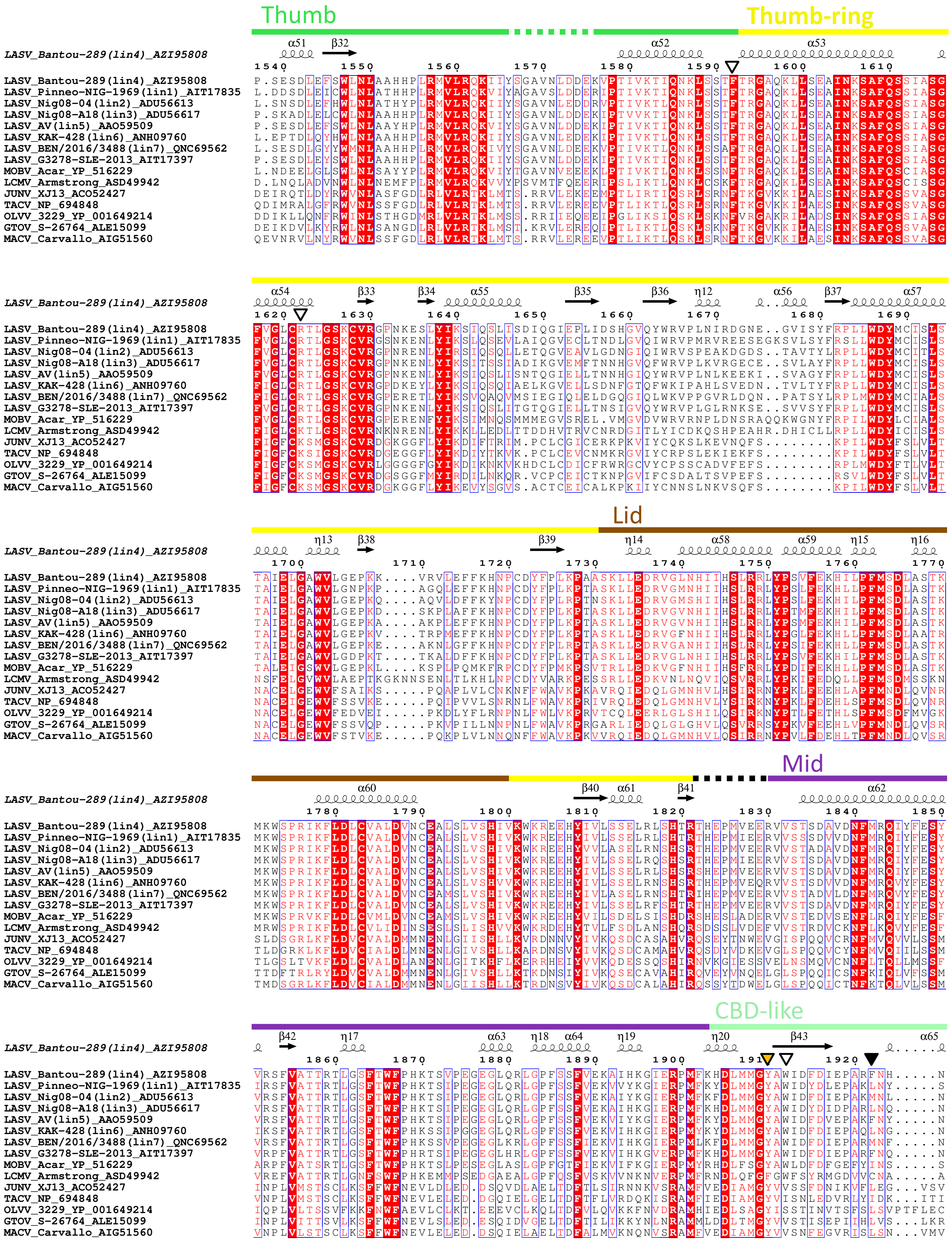


▼ – general defect in L protein activity upon mutation observed in Lassa virus mini-replicon system

▼ (orange) – selective transcriptional defect of L protein upon mutation observed in Lassa virus mini-replicon system

▽ – no or weak effect on L protein activity upon mutation observed in Lassa virus mini-replicon system

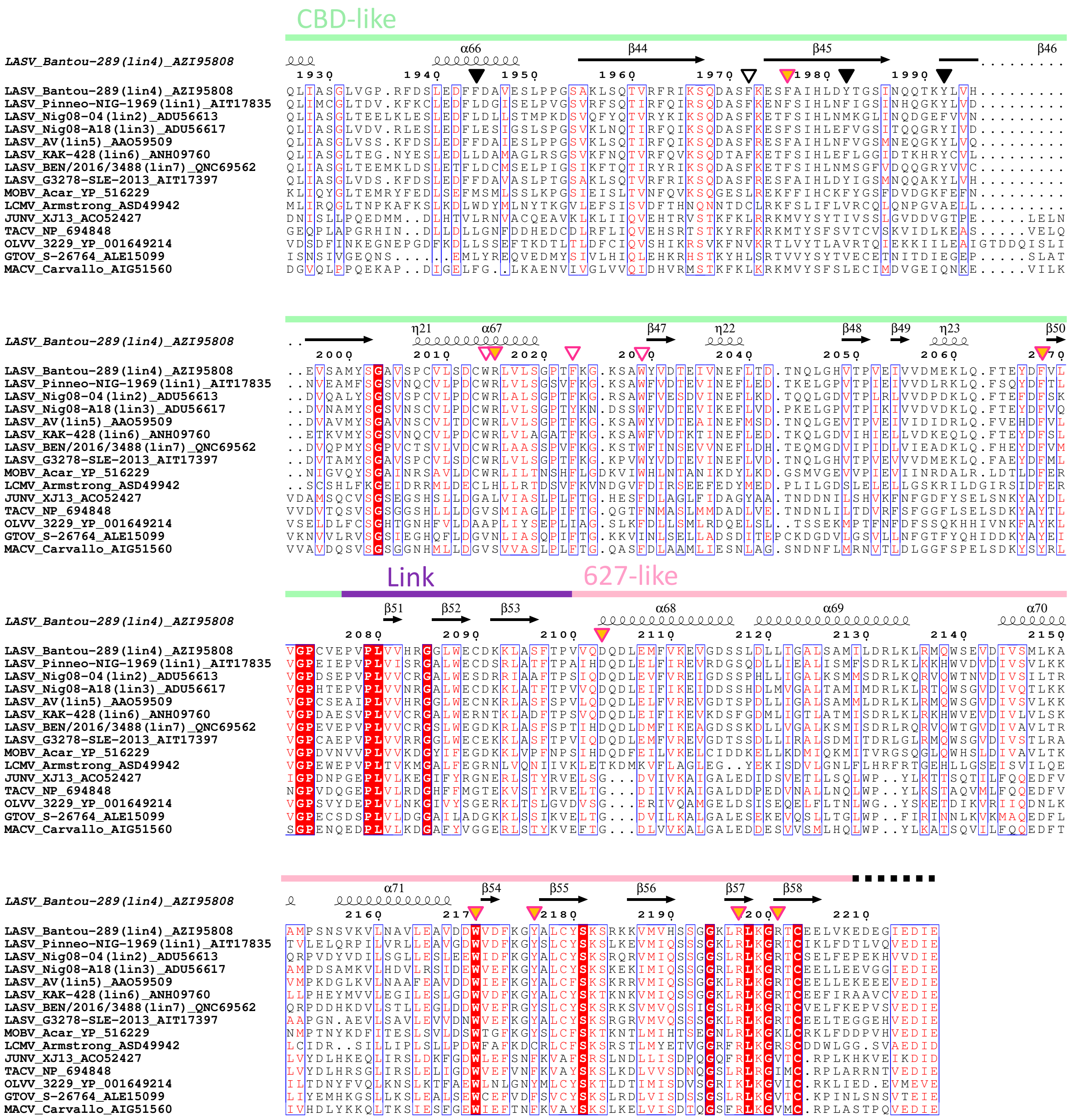


▽ – selective transcriptional defect of L protein upon mutation observed in Lassa virus mini-replicon system (Lehmann et al. 2014)

▽ – no or weak effect on L protein activity upon mutation observed in Lassa virus mini-replicon system (Lehmann et al. 2014)

▼ – general defect in L protein activity upon mutation observed in Lassa virus mini-replicon system

▼ – selective transcriptional defect of L protein upon mutation observed in Lassa virus mini-replicon system

▽ – no or weak effect on L protein activity upon mutation observed in Lassa virus mini-replicon system

▽ – intermediate effect on L protein activity upon mutation observed in Lassa virus mini-replicon system
