## Supplementary Cryo-EM processing overview file for "Conformational changes in Lassa virus L protein associated with promoter binding and RNA synthesis activity"

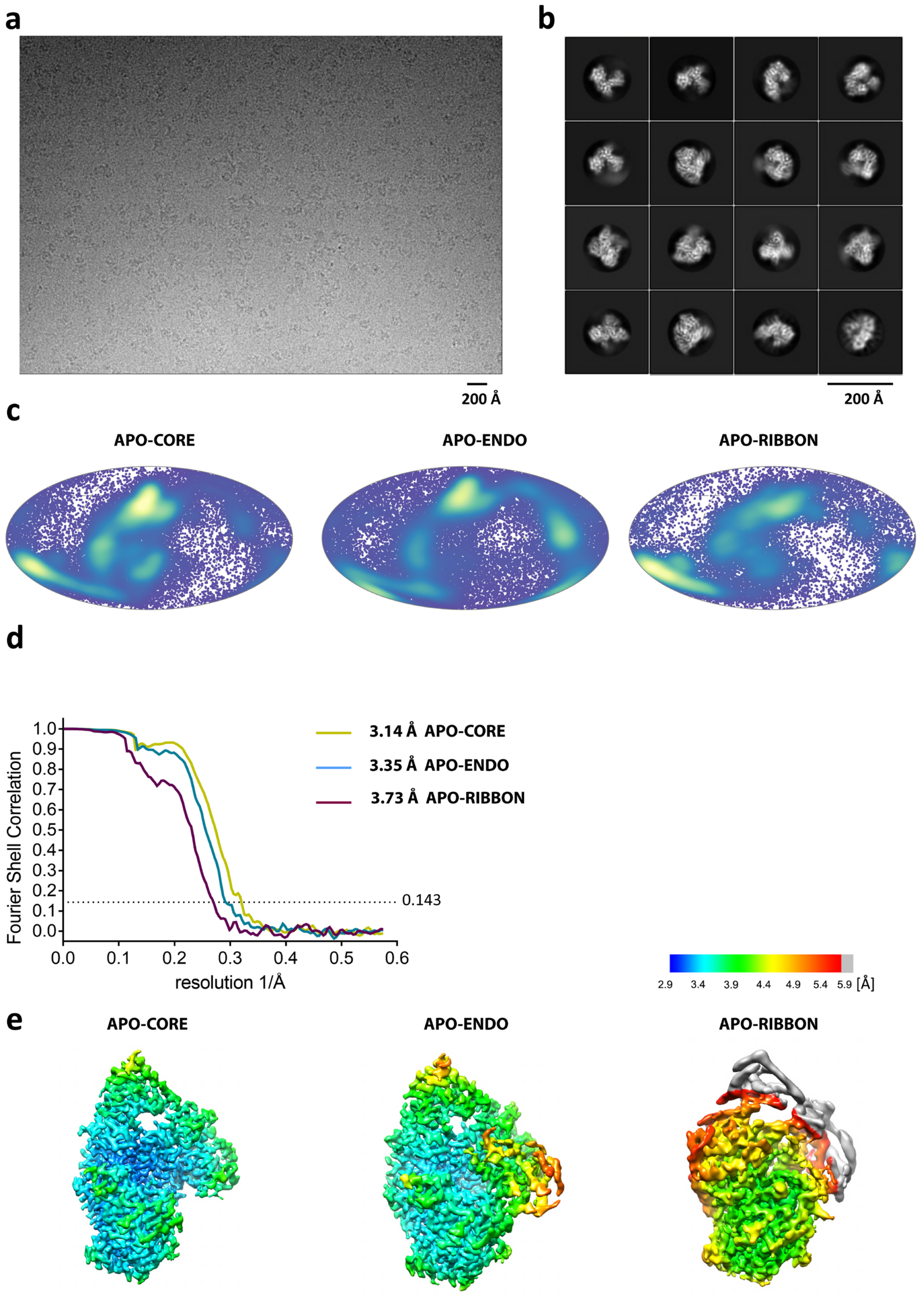


**Figure EM-1: Cryo-EM of APO LASV L protein**

**a**, Representative micrograph of APO LASV L protein in free standing ice after MotionCor2 [1] correction at defocus of ~2.5 µm. **b**, 2D-class averages of the APO LASV L protein complex. **c**, Angular distribution for particle projections of the APO-CORE, APO-ENDO and APO-RIBBON, respectively, visualized on a globe-like plane. **d**, Fourier shell correlation (FSC) curves for the APO-CORE (yellow), APO-ENDO (cyan) and APO-RIBBON (purple), respectively. The plot of the FSC between two independently refined half-maps shows the overall resolution of the two maps as indicated by the gold standard FSC 0.143 cut-off criteria[2]. **e**, Surface representation of local resolution distribution of the APO-CORE, APO-ENDO and APO-RIBBON, respectively. Maps are colored according to the local resolution calculated within the RELION software package. Resolution is as indicated in the color bar.

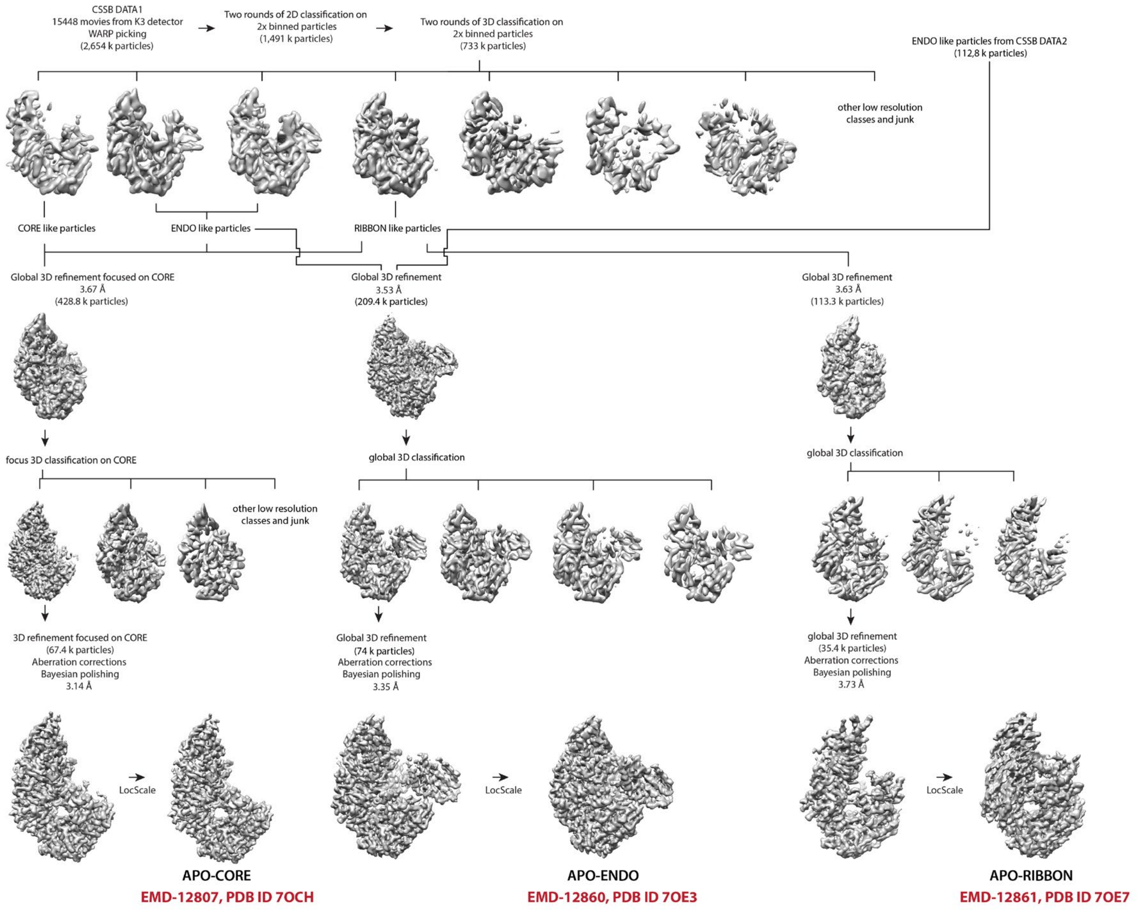


**Figure EM-2: Cryo-EM data 3D classification and refinement scheme of APO LASV L protein.**

Summary of the cryo-EM 3D classification and refinement scheme of the APO LASV L protein. CSSB DATA1 2D classes with well-defined secondary structure features were merged (733k particles). The merged particles were classified into ten 3D classes with angular assignment. Incomplete, low resolution, and damaged particle classes were excluded from further data analyses. Three most prominent 3D classes of the APO LASV L protein corresponding to APO-CORE, APO-ENDO and APO-RIBBON were identified. APO-CORE was focus-3D classified around the CORE region of the LASV L protein and the best defined class was focus 3D-refined. APO-ENDO was merged with the same class from CSSB DATA2, globally 3D-refined, and further 3D classified. Best defined APO-ENDO class was globally 3D-refined. APO-RIBBON was processed similarly to APO-ENDO. Final cryo-EM maps were refined and post-processed with their respective masks in RELION 3.1 [3,4] and filtered by LocScale [5].

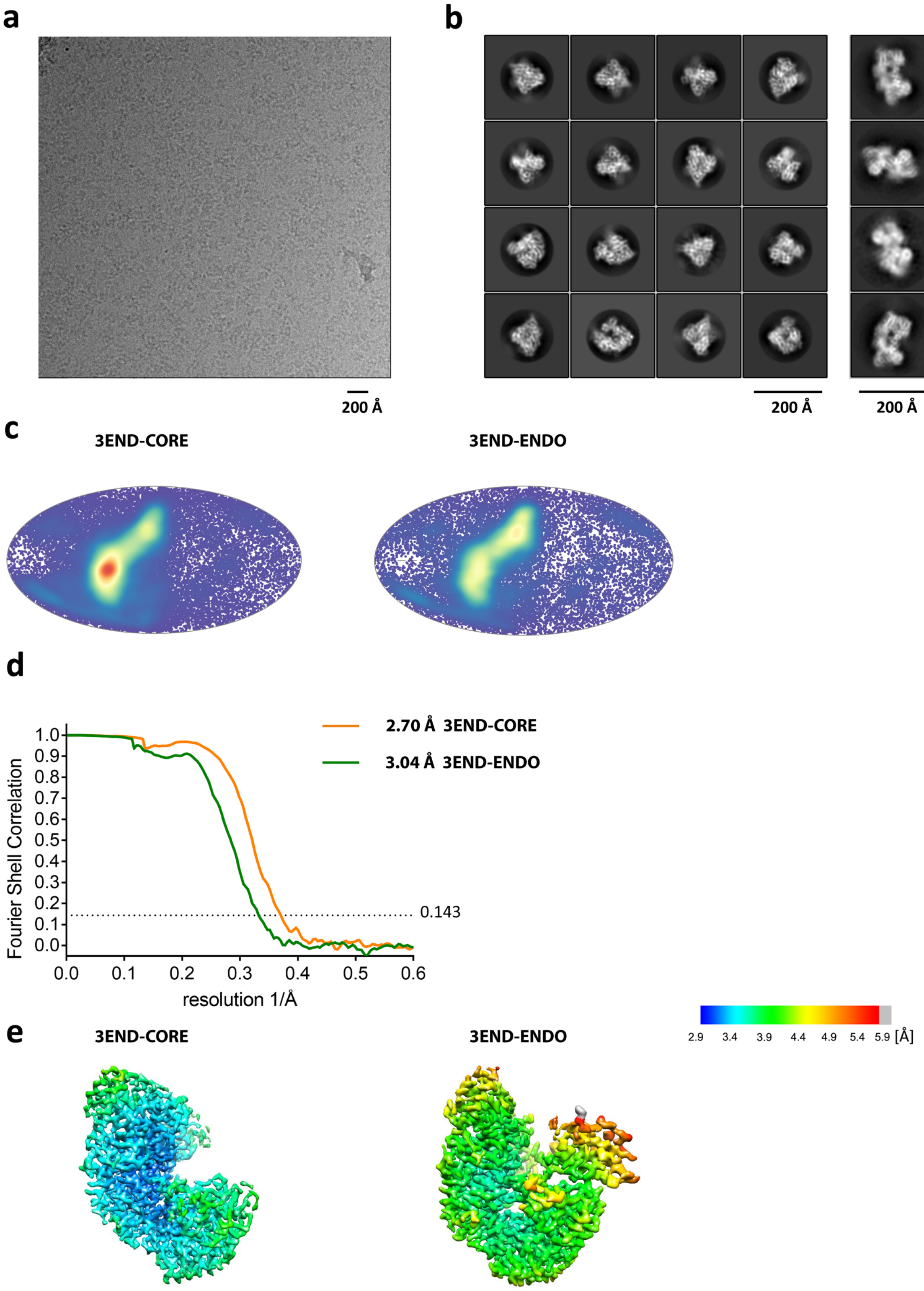


**Figure EM-3: Cryo-EM of LASV L protein with vRNA 3' end alone**

**a**, Representative micrograph of LASV L protein with vRNA 3' end alone in free standing ice after MotionCor2[1] correction at defocus of ~2.5 µm. **b**, 2D-class averages of the of LASV L protein with vRNA 3' end alone - in monomeric (**left**) and dimeric (**right**) form. **c**, Angular distribution for particle projections of the 3END-CORE and 3END-ENDO, respectively, visualized on a globe-like plane. **d**, Fourier shell correlation (FSC) curves for the 3END-CORE (orange), 3END-ENDO (green), respectively. The plot of the FSC between two independently refined half-maps shows the overall resolution of the two maps as indicated by the gold standard FSC 0.143 cut-off criteria[2]. **e**, Surface representation of local resolution distribution of the 3END-CORE, 3END-ENDO, respectively. Maps are colored according to the local resolution calculated within the RELION software package. Resolution is as indicated in the color bar.

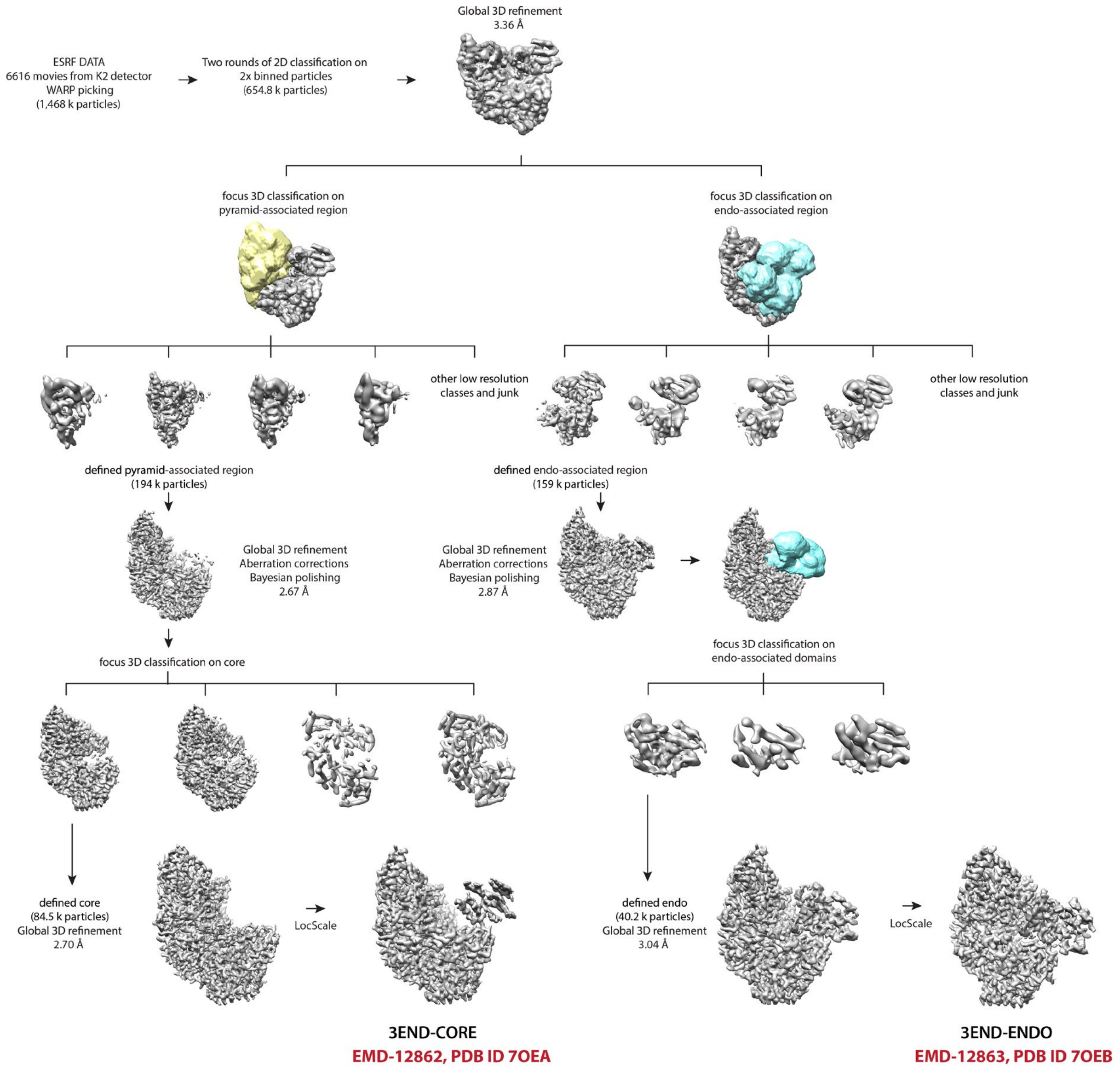


**Figure EM-4: Cryo-EM data 3D classification and refinement scheme of LASV L protein with vRNA 3' end alone**

Summary of the cryo-EM 3D classification and refinement scheme of of LASV L protein with vRNA 3' end alone. ESRF DATA 2D classes with well-defined secondary structure features were merged (654.8k particles). The merged particles were globally 3D-refined and then in parallel focus 3D-classified around the pyramid-associated domains (**left**, yellow) and endo-associated domains (**right**, blue). The most defined class of pyramid-associated domains was globally 3D-refined and further 3D-classified around the whole core region. The most defined class was globally 3D-refined and assigned as 3END-CORE. The most defined class of endo-associated domains was globally 3D-refined and further 3D-classified strictly around the endo domain. The most defined endo domain class was globally 3D-refined and assigned as 3END-ENDO. Final cryo-EM maps were refined and post-processed with their respective masks in RELION 3.1[3,4] and filtered by LocScale [5].

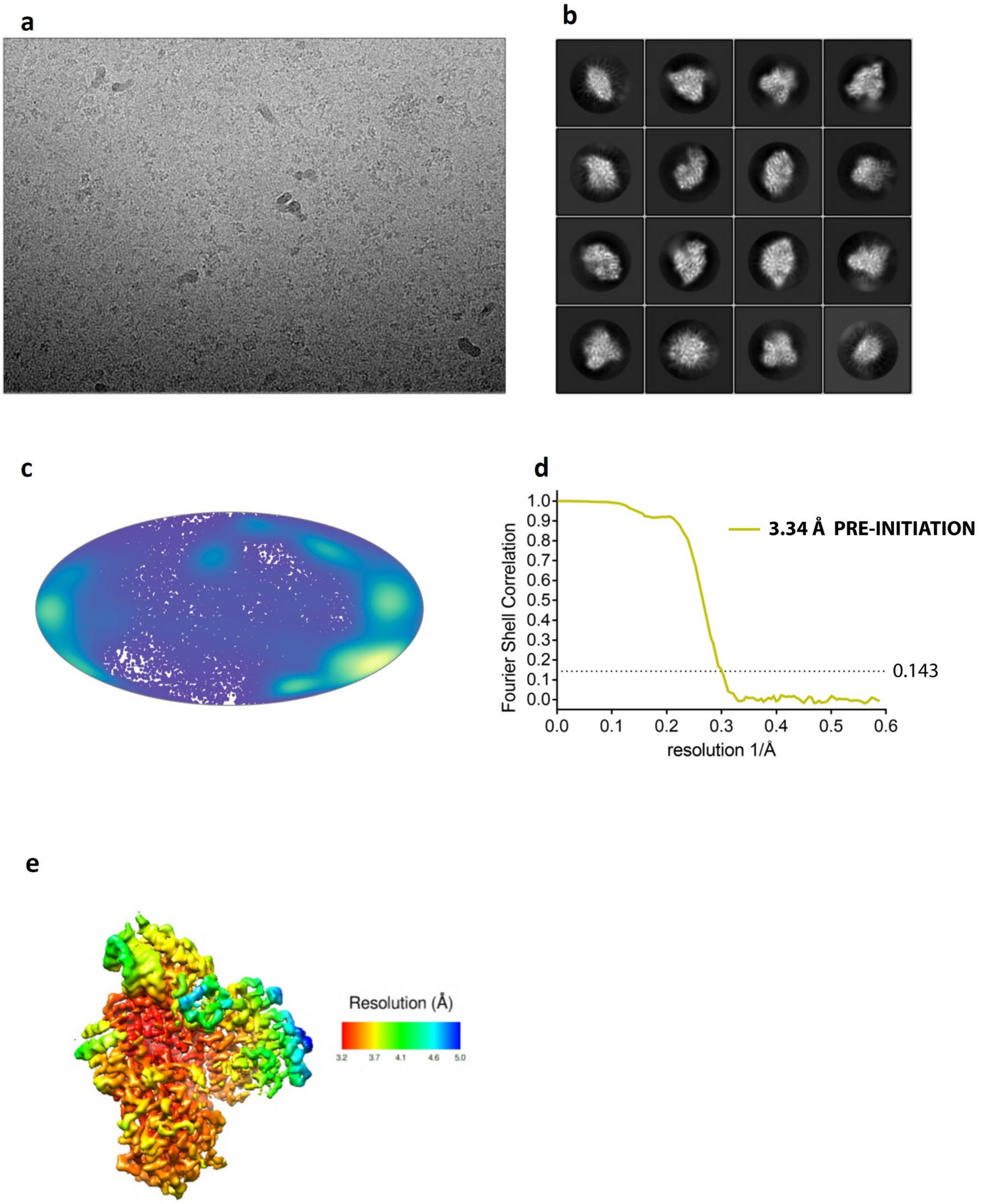


**Figure EM-5: Cryo-EM of LASV L protein PRE-INITIATION complex**

**a**, Representative micrograph of LASV L protein with 3’ and 5’ vRNAs in free standing ice after MotionCor2[1] correction at defocus of ~2.5 µm. **b**, 2D-class averages of the of LASV L protein with 3’ and 5’ vRNAs **c**, Angular distribution for particle projections of PRE-INITIATION visualized on a globe-like plane. **d**, Fourier shell correlation (FSC) curves for PRE-INITIATION. The plot of the FSC between two independently refined half-maps shows the overall resolution of the two maps as indicated by the gold standard FSC 0.143 cut-off criteria[2]. **e**, Surface representation of local resolution distribution of PRE-INITIATION. The map is colored according to the local resolution calculated within the RELION software package. Resolution is as indicated in the color bar.

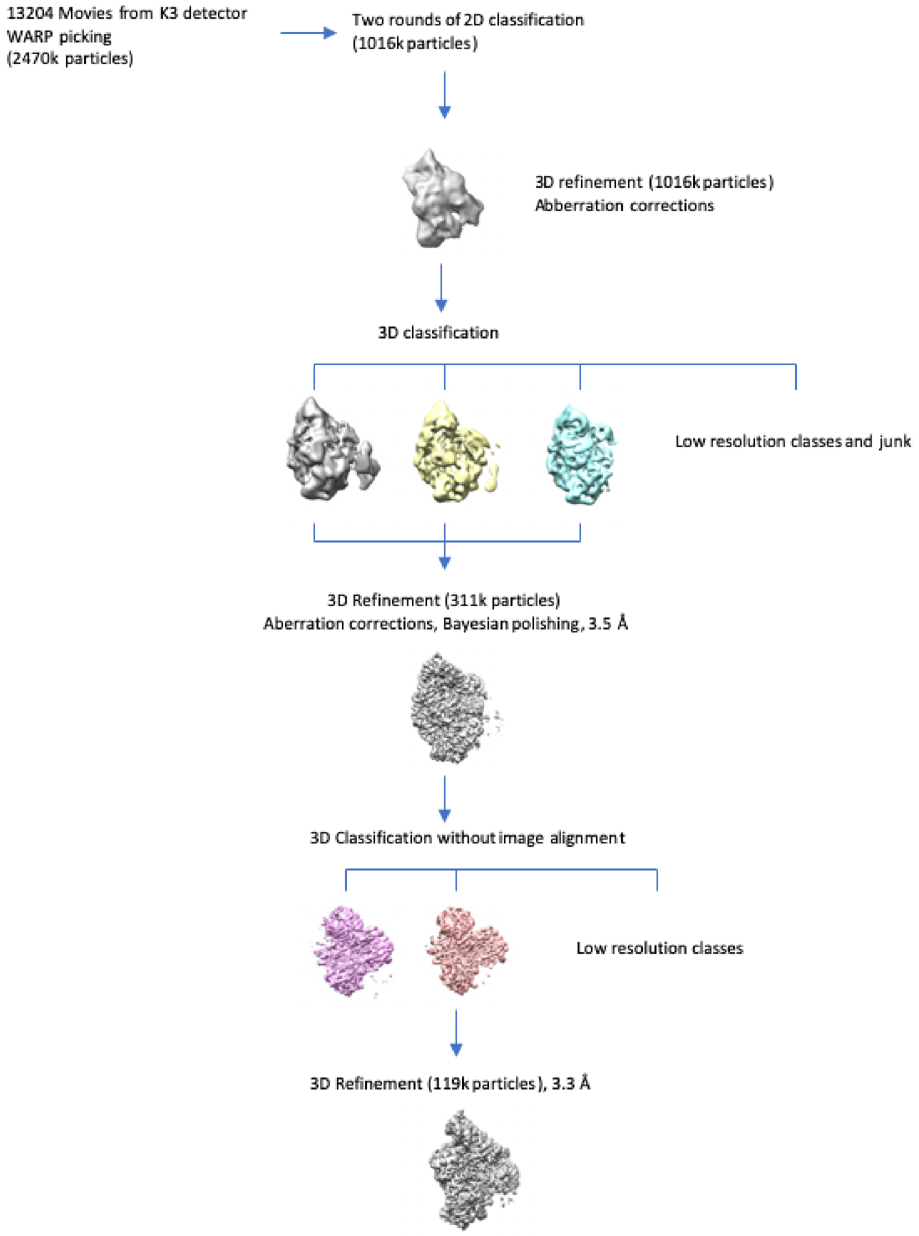


**Figure EM-6: Cryo-EM data 3D classification and refinement scheme of LASV L protein PRE-INITIATION complex**

Summary of the cryo-EM 3D classification and refinement scheme of the PRE-INITIATION LASV L protein. CSSB DATA3 2D classes with well-defined secondary structure features were merged (1016k particles). The merged particles were classified into twelve 3D classes with angular assignment. Incomplete, low resolution, and damaged particle classes were excluded from further data analyses. Three most prominent 3D classes were merged globally 3D-refined, aberration corrected and Bayesian polished [3,4] then further 3D classified. The best defined class was globally 3D-refined. Final cryo-EM map was refined and post-processed with their respective mask in RELION 3.1.

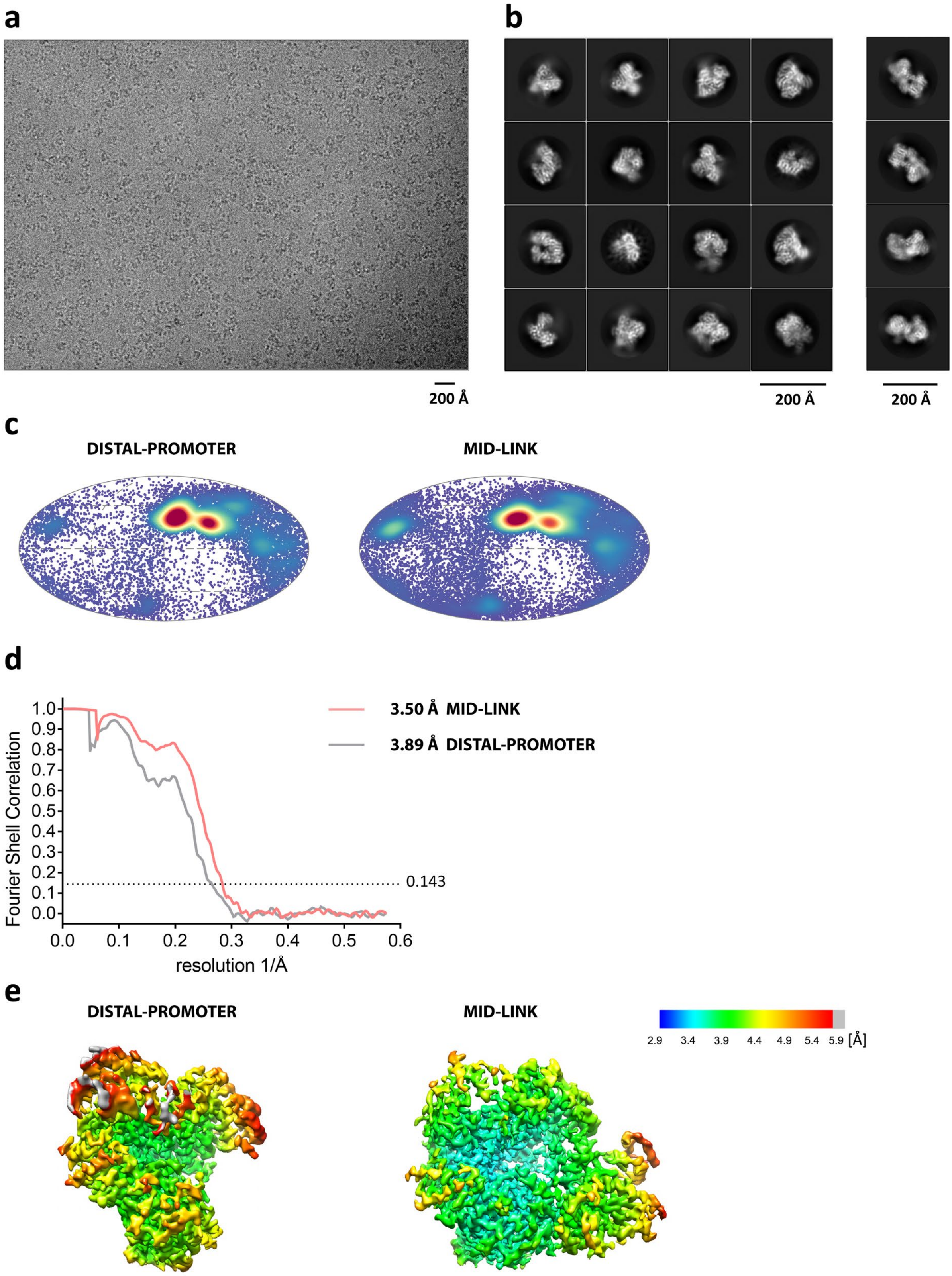


**Figure EM-7: Cryogenic electron microscopy of LASV L protein incubated with a truncated promoter**

**a**, Representative micrograph of LASV L protein incubated with a truncated promoter in free standing ice after MotionCor2[1] correction at defocus of ~2.5 µm. **b**, 2D-class averages of the of LASV L protein incubated with a truncated promoter complex in monomeric (**left**) and dimeric (**right**) form. **c**, Angular distribution for particle projections of the DISTAL-PROMOTER, MID-LINK, respectively, visualized on a globe-like plane. **d**, Fourier shell correlation (FSC) curves for the DISTAL-PROMOTER (grey), MID-LINK (pink), respectively. The plot of the FSC between two independently refined half-maps shows the overall resolution of the two maps as indicated by the gold standard FSC 0.143 cut-off criteria[2]. **e**, Surface representation of local resolution distribution of the DISTAL-PROMOTER, MID-LINK, respectively. Maps are colored according to the local resolution calculated within the RELION software package. Resolution is as indicated in the color bar.

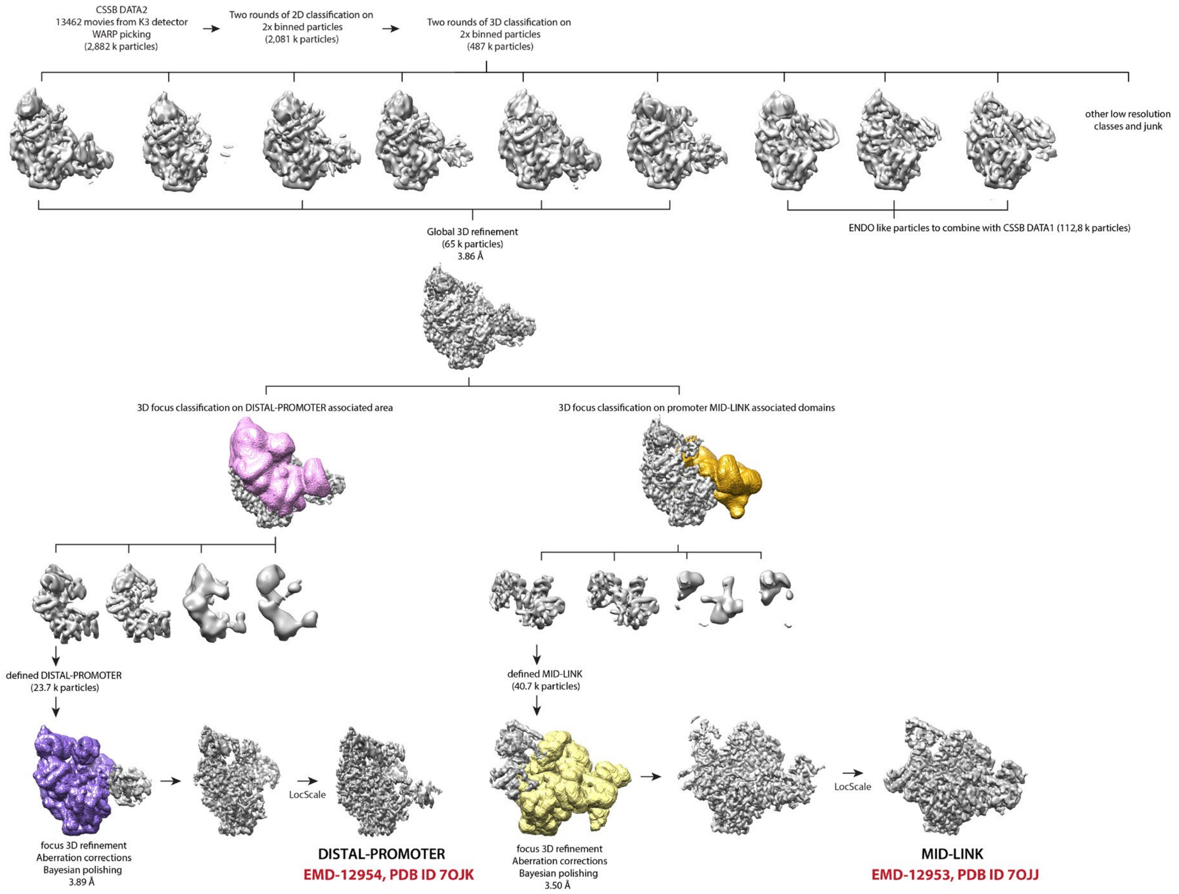


**Figure EM-8: Cryo-EM data 3D classification and refinement scheme LASV L protein incubated with a truncated promoter**

Summary of the cryo-EM 3D classification and refinement scheme of the LASV L protein incubated with a truncated promoter. CSSB DATA2 2D classes with well-defined secondary structure features were merged (2,081k particles). The merged particles were classified into twelve 3D classes with angular assignment. Incomplete, low resolution, and damaged particle classes were excluded from further data analyses. Three classes, which possessed defined endo domain, were pooled with CSSB DATA1 (Figure EM2). Four classes, which possessed defined periphery around putative promoter-duplex binding site and mid-link domain, were merged and globally 3D-refined. The data were then in parallel focus 3D-classified around the DISTAL-PROMOTER regions and associated domains (**left**, pink) and MID-LINK-associated domains (**right**, orange). The most defined class of DISTAL-PROMOTER region was focused 3D-refined, excluding the MID-LINK region (purple). The most defined MID-LINK domain class was focus 3D-refined, excluding the DISTAL-PROMOTER region (yellow). Final cryo-EM maps were refined and post-processed with their respective masks in RELION 3.1[3,4] and filtered by LocScale1[5].

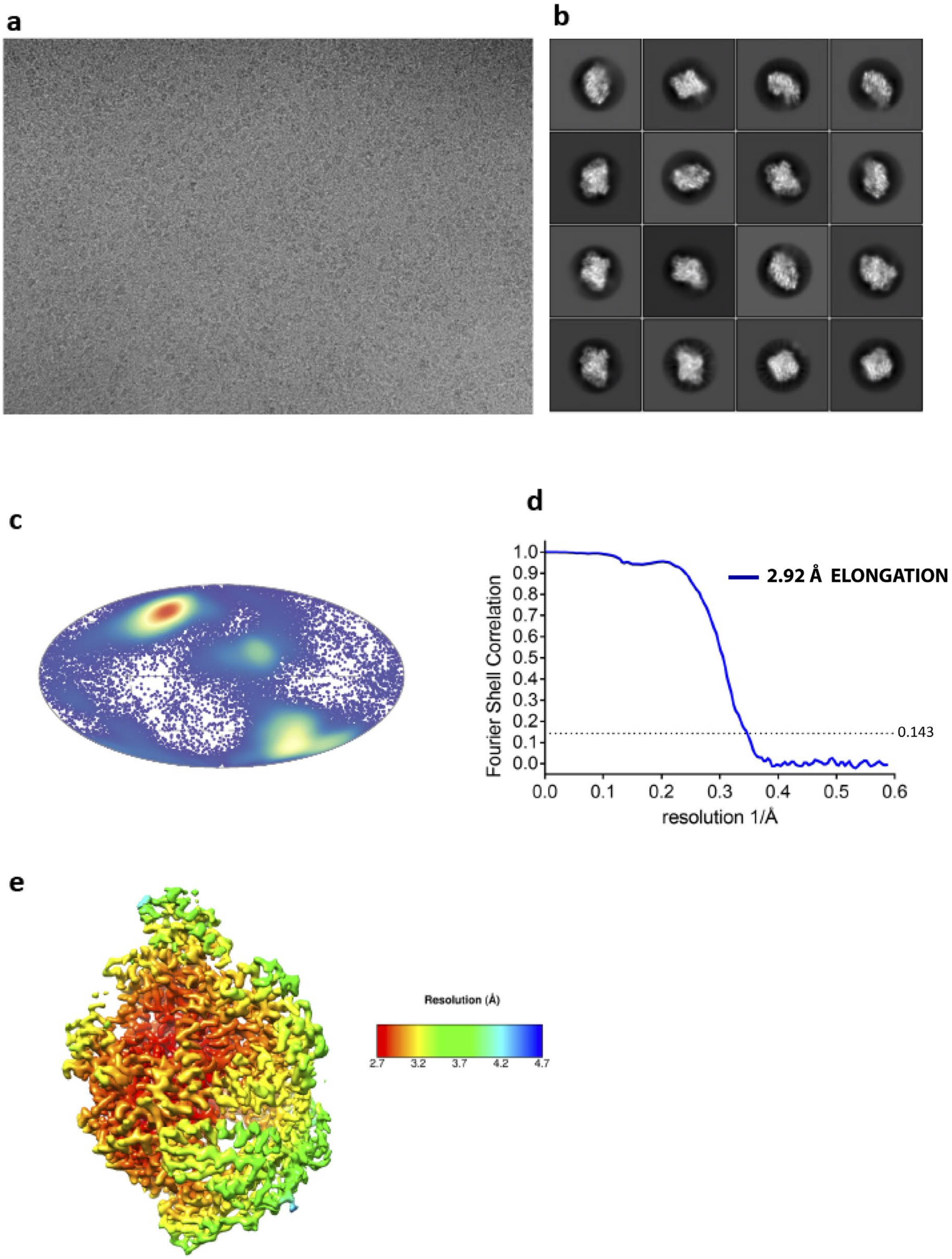


**Figure EM-9: Cryo-EM of LASV L protein ELONGATION complex**

**a**, Representative micrograph of LASV L protein ELONGATION complex in free standing ice after MotionCor2[1] correction at defocus of ~2.0 µm. **b**, 2D-class averages of the of LASV L protein ELONGATION complex **c**, Angular distribution for particle projections of the ELONGATION complex visualized on a globe-like plane. **d**, Fourier shell correlation (FSC) curves for the ELONGATION. The plot of the FSC between two independently refined half-maps shows the overall resolution of the two maps as indicated by the gold standard FSC 0.143 cut-off criteria[2].**e**, Surface representation of local resolution distribution of the ELONGATION. Maps are colored according to the local resolution calculated within the RELION software package. Resolution is as indicated in the color bar.

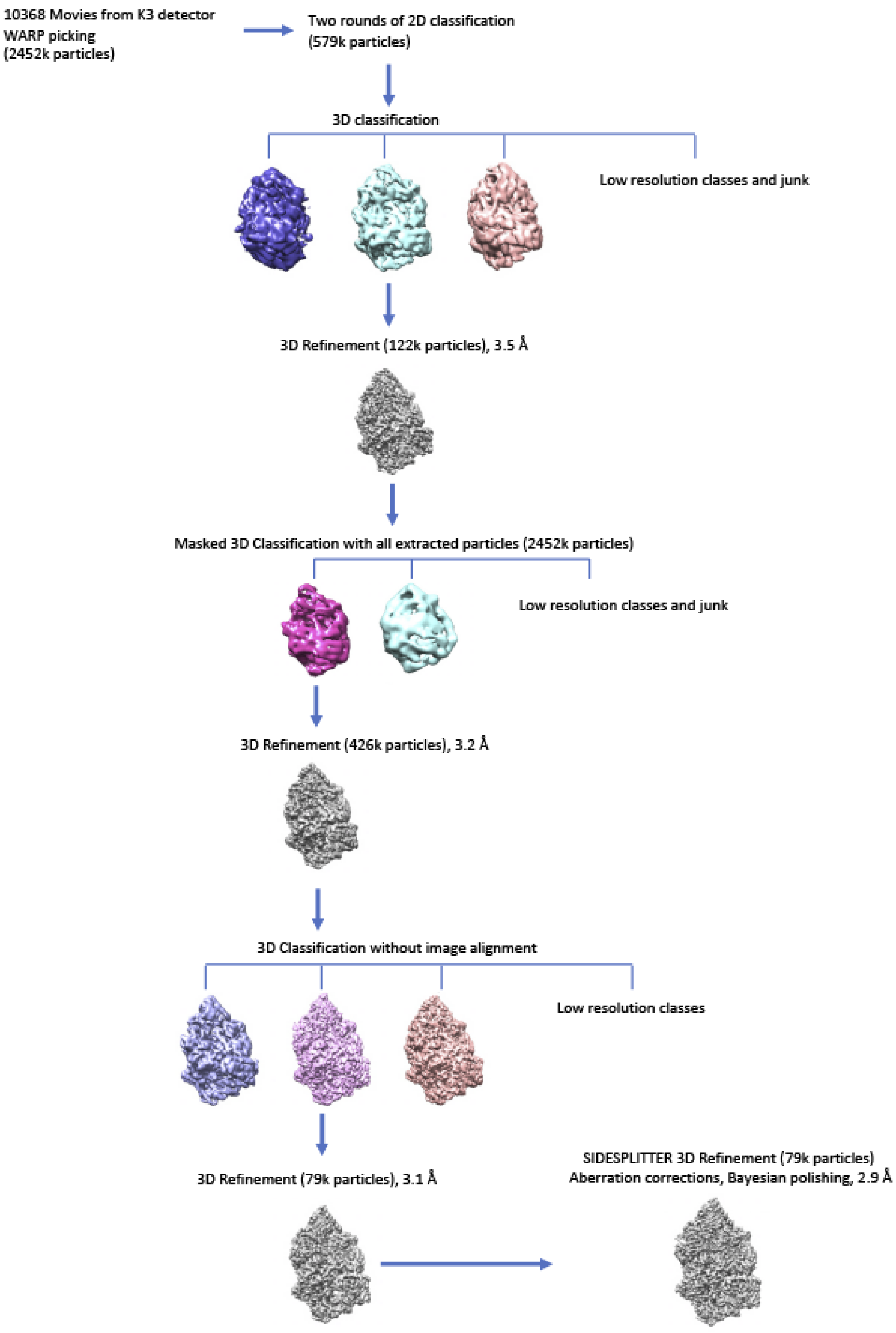
10368 Movies from K3 detector
WARP picking
(2452k particles)
Two rounds of 2D classification
(579k particles)
3D classification
Low resolution classes and junk
3D Refinement (122k particles), 3.5 Å
Masked 3D Classification with all extracted particles (2452k particles)
Low resolution classes and junk
3D Refinement (426k particles), 3.2 Å
3D Classification without image alignment
Low resolution classes
3D Refinement (79k particles), 3.1 Å
SIDESPLITTER 3D Refinement (79k particles)
Aberration corrections, Bayesian polishing, 2.9 Å

143

**Figure EM-10: Cryo-EM data 3D classification and refinement scheme of LASV L ELONGATION complex**

Summary of the cryo-EM 3D classification and refinement scheme of the ELONGATION LASV L protein. CSSB DATA4 2D classes with well-defined secondary structure features were merged (579k particles). The merged particles were classified into ten 3D classes with angular assignment. The highest resolution class was 3D-Refined and then used as a reference for 3D classification with ten classes of all extracted particles (2452k). The most prominent 3D class was globally 3D-refined then further 3D classified then further 3D classified. The most defined class was globally 3D-refined. The final cryo-EM map was refined with SIDESPLITTER[6]and post-processed with their respective mask in RELION 3.1[3,4].

### Refrences:


1 Zheng, S. Q. *et al.* MotionCor2: anisotropic correction of beam-induced motion for improved cryo-electron microscopy. *Nat Methods* **14**, 331-332, doi:10.1038/nmeth.4193 (2017).

2 Rosenthal, P. B. & Henderson, R. Optimal determination of particle orientation, absolute hand, and contrast loss in single-particle electron cryomicroscopy. *J Mol Biol* **333**, 721-745 (2003).

3 Zivanov, J. *et al.* New tools for automated high-resolution cryo-EM structure determination in RELION-3. *Elife* **7**, doi:10.7554/eLife.42166 (2018).

4 Zivanov, J., Nakane, T. & Scheres, S. H. W. Estimation of high-order aberrations and anisotropic magnification from cryo-EM data sets in RELION-3.1. *IUCrJ* **7**, 253-267, doi:10.1107/S2052252520000081 (2020).

5 Jakobi, A. J., Wilmanns, M. & Sachse, C. Model-based local density sharpening of cryo-EM maps. *Elife* **6**, doi:10.7554/eLife.27131 (2017).

6 Ramlaul, K., Palmer, C. M., Nakane, T. & Aylett, C. H. S. Mitigating local over-fitting during single particle reconstruction with SIDESPLITTER. *J Struct Biol* **211**, 107545, doi:10.1016/j.jsb.2020.107545 (2020).
