## Supplementary Fig for "Conformational changes in Lassa virus L protein associated with promoter binding and RNA synthesis activity"

### Supplement

Supplementary Fig. 1

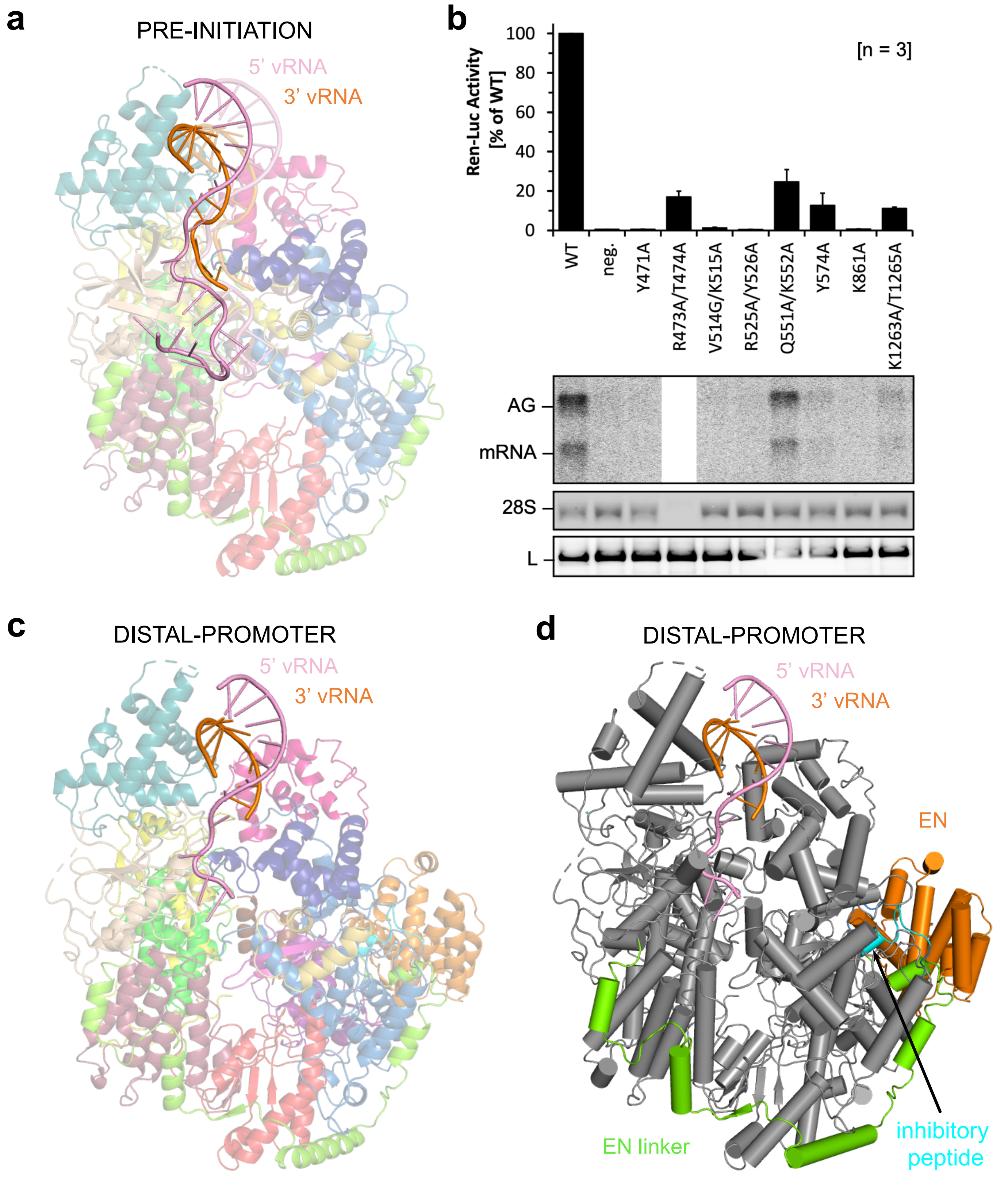

Supplementary Fig. 2

a

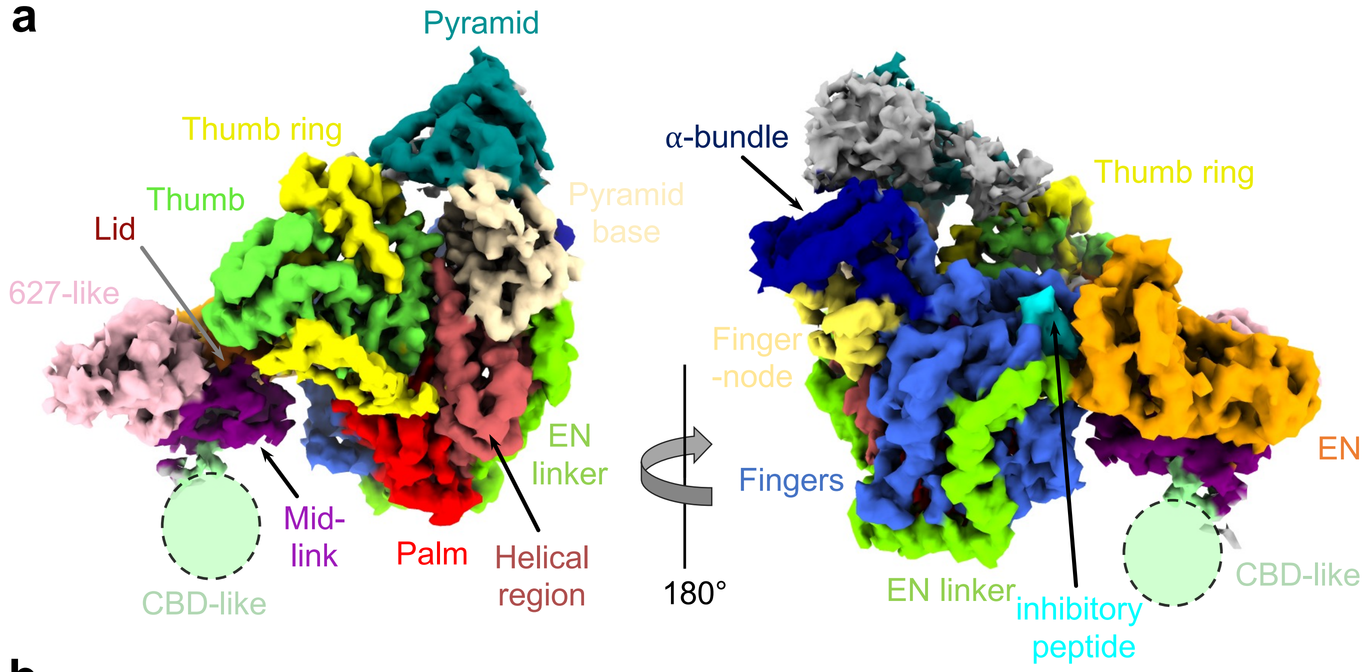


b

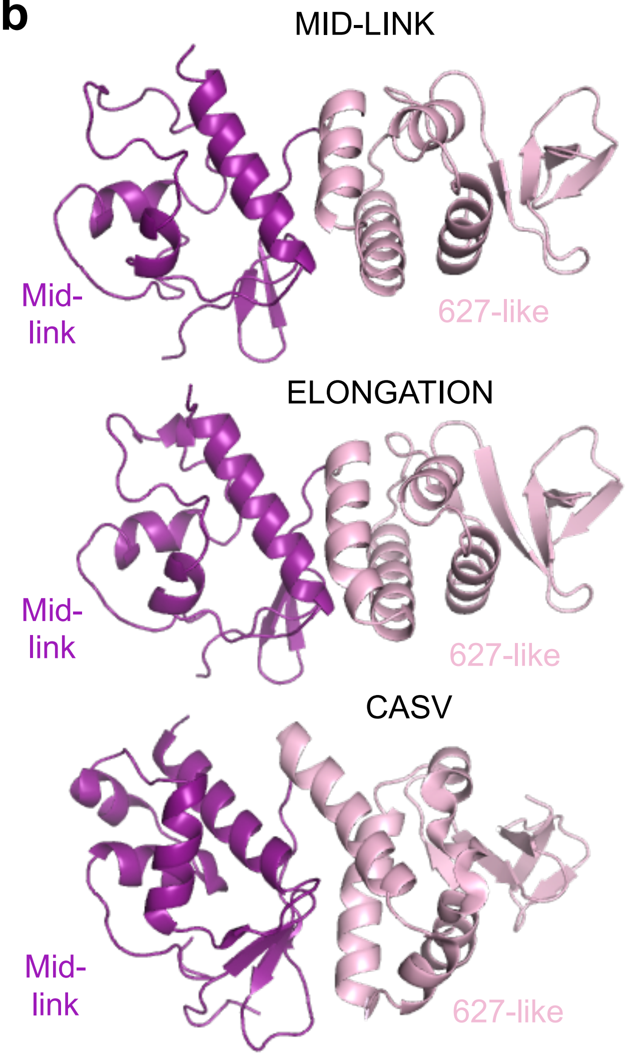

### Supplementary Fig. 3

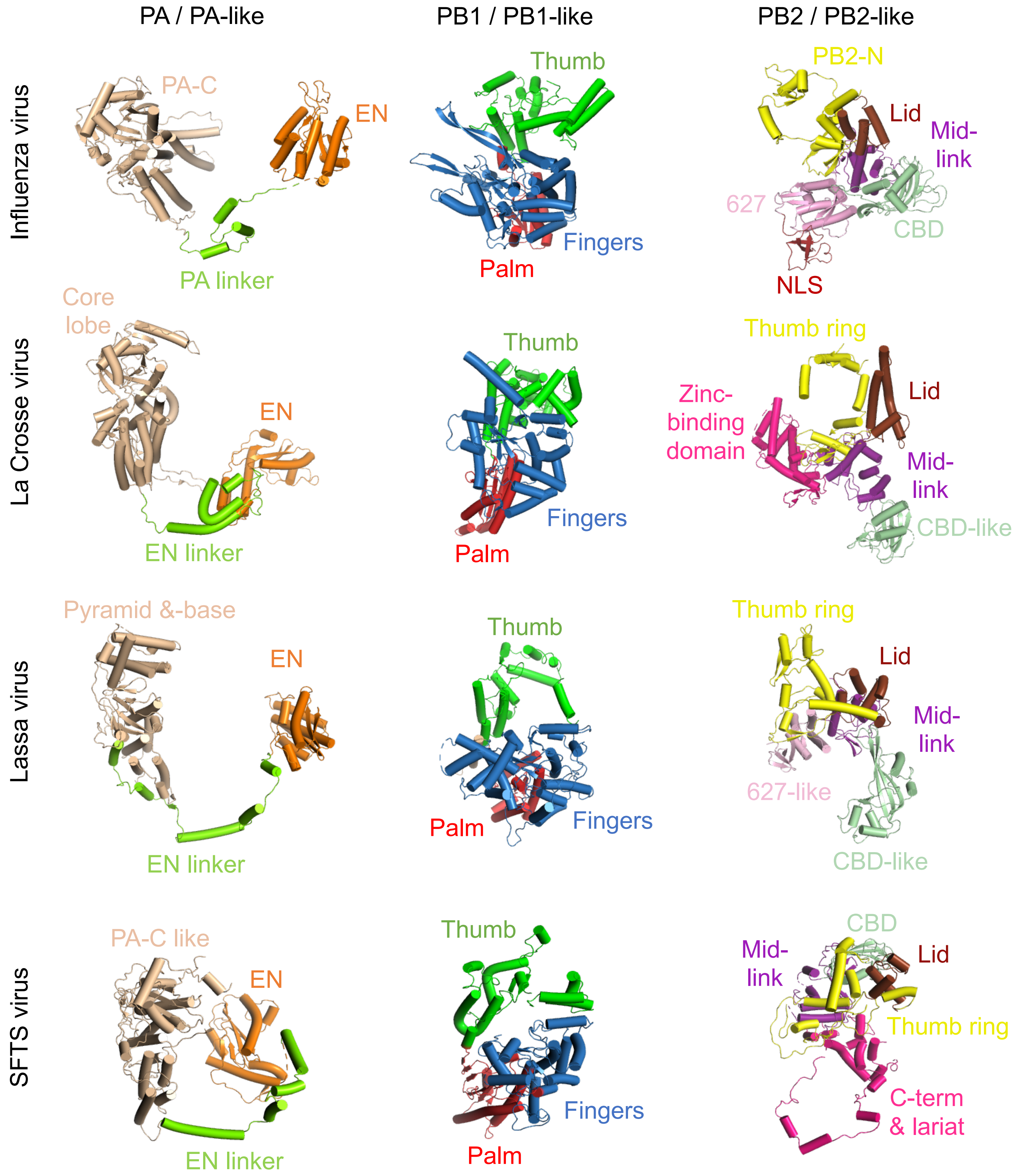

### Supplementary Fig. 4

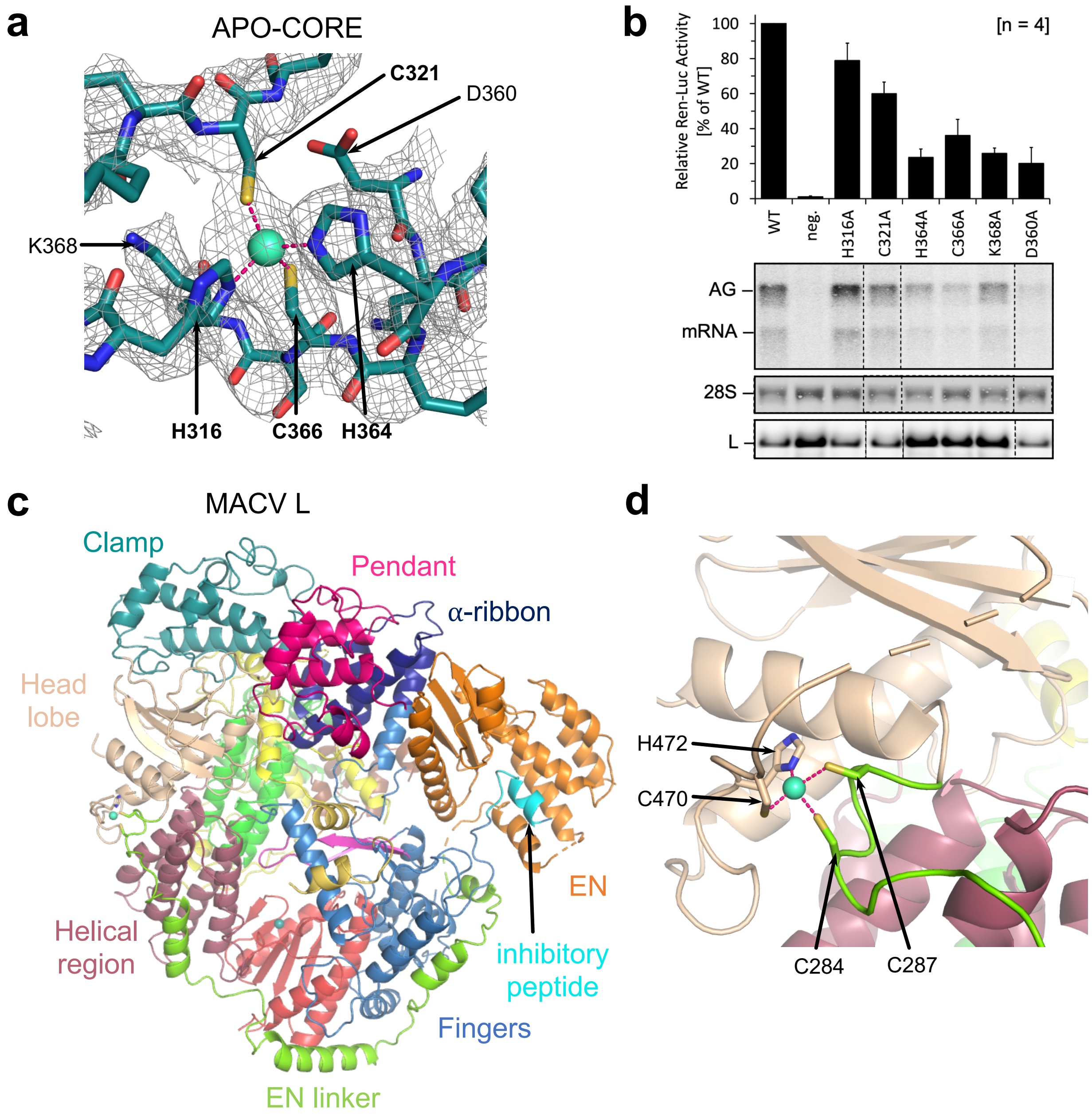

### Supplementary Fig. 5

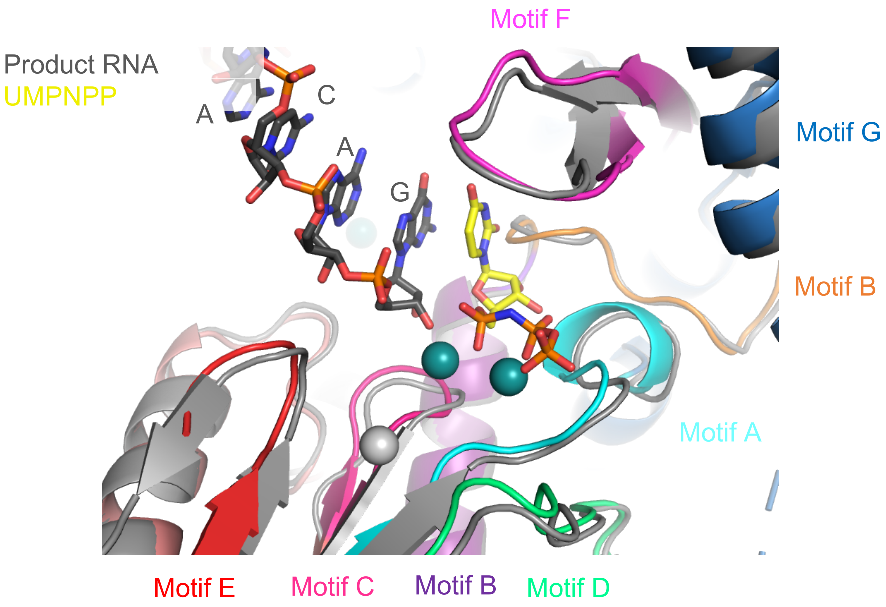

### Supplementary Fig. 6

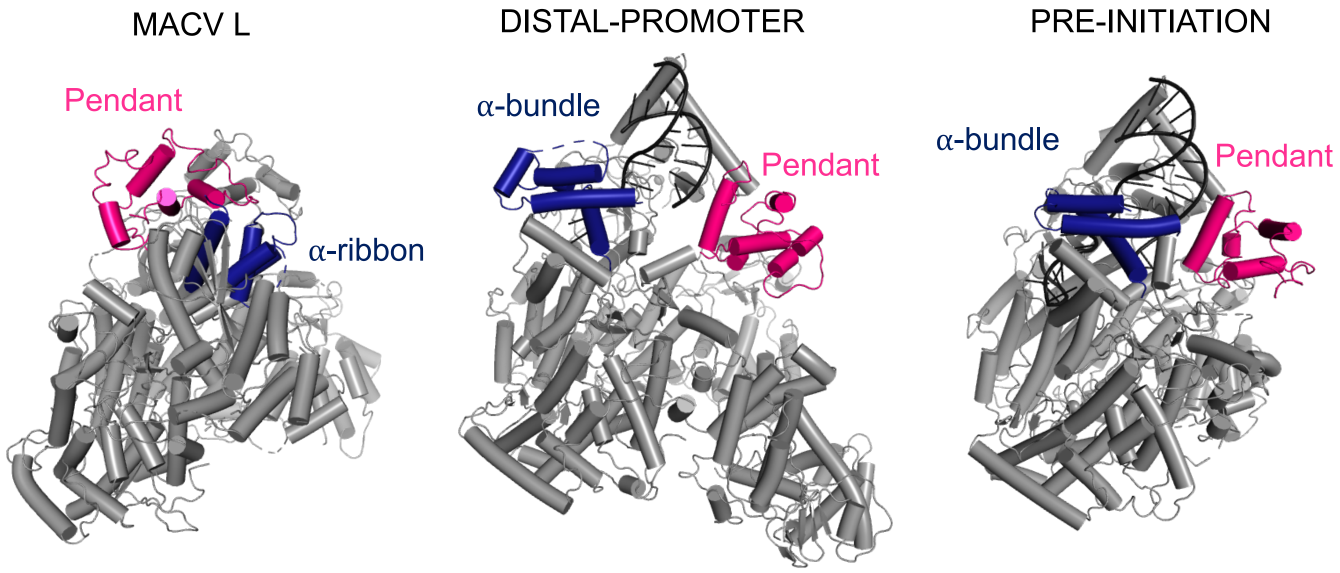

Supplementary Fig. 7

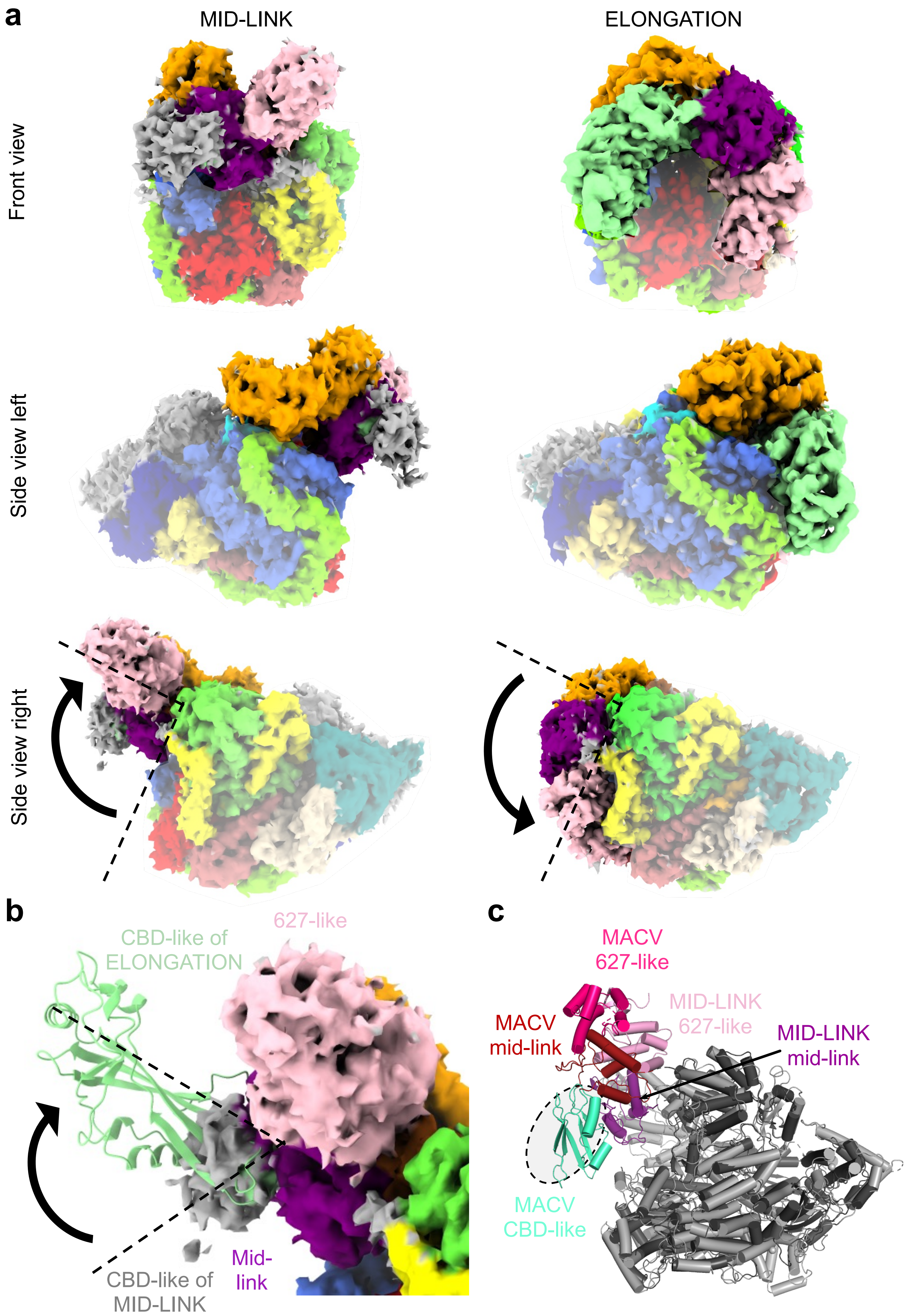

### Supplementary Fig. 8

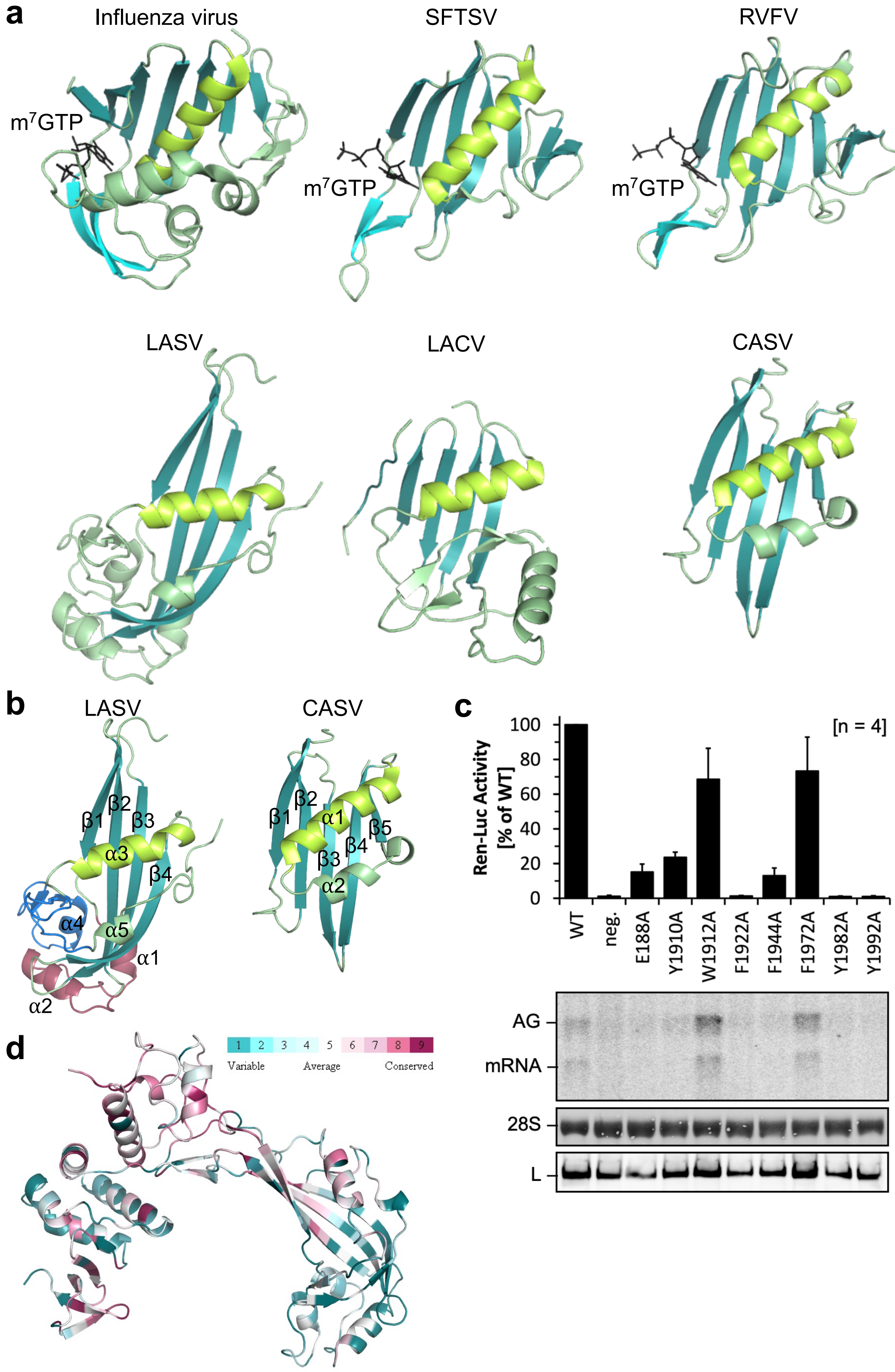

Supplementary Fig. 9

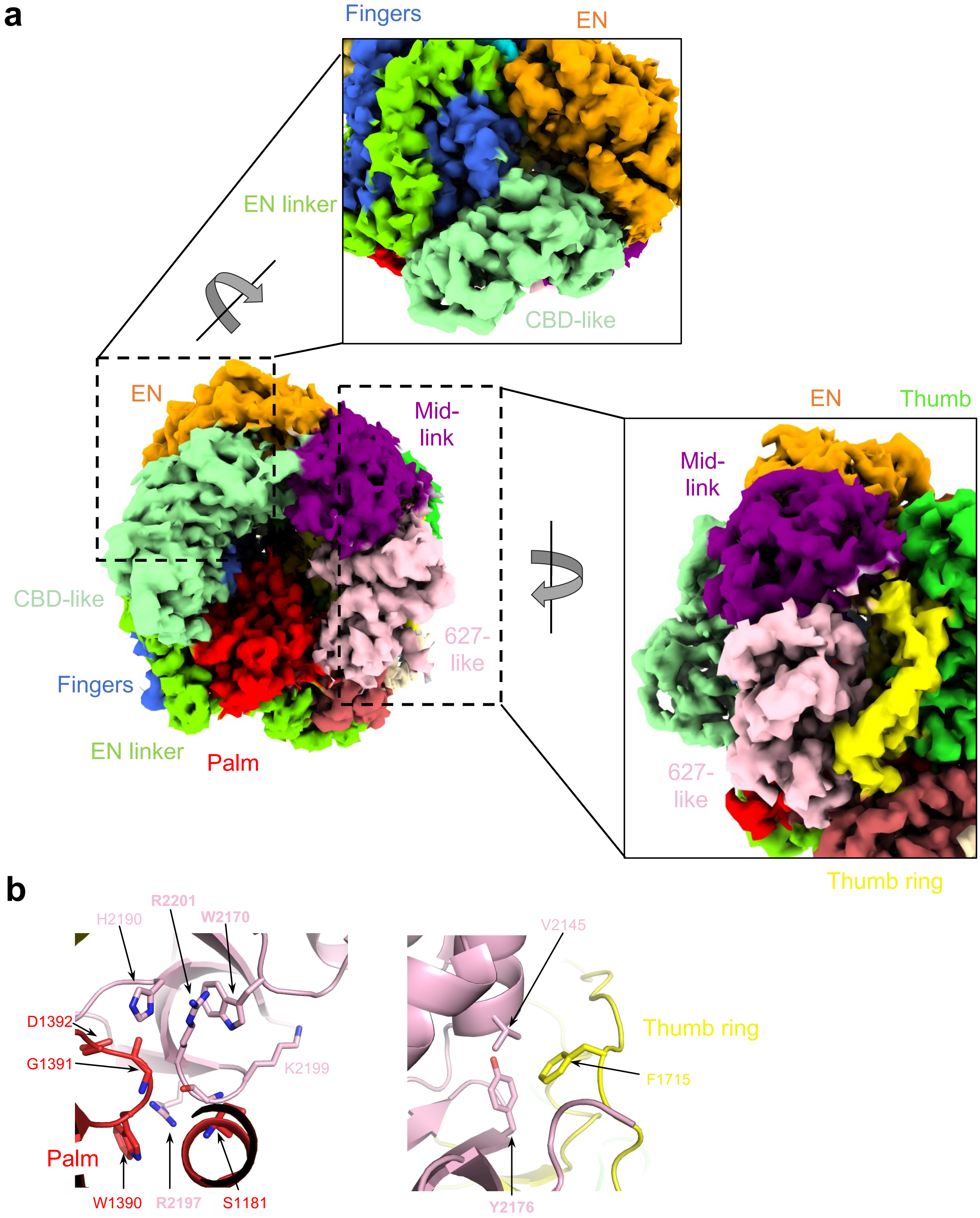

### Supplementary Fig. 10

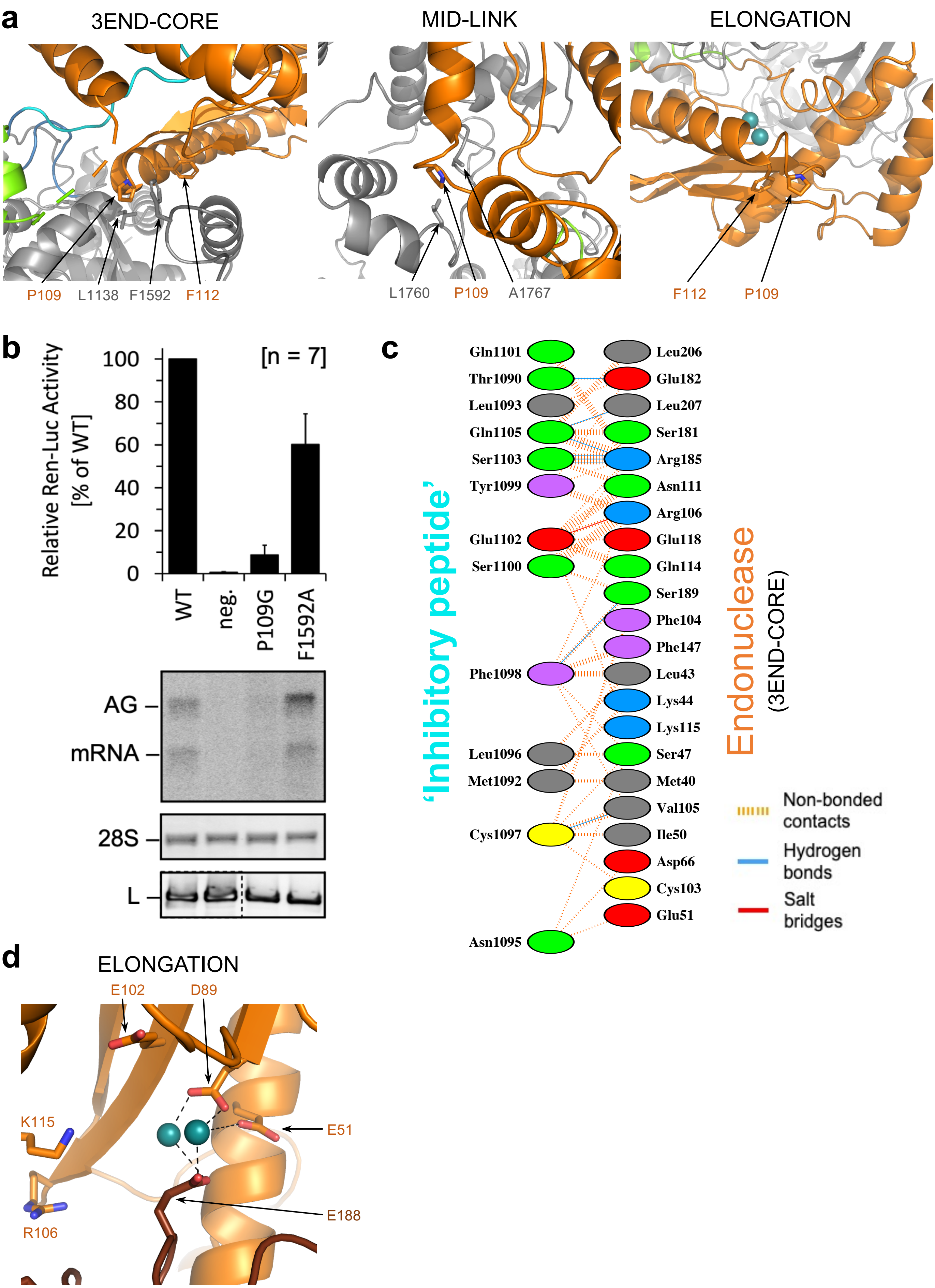

### Supplementary Fig. 11

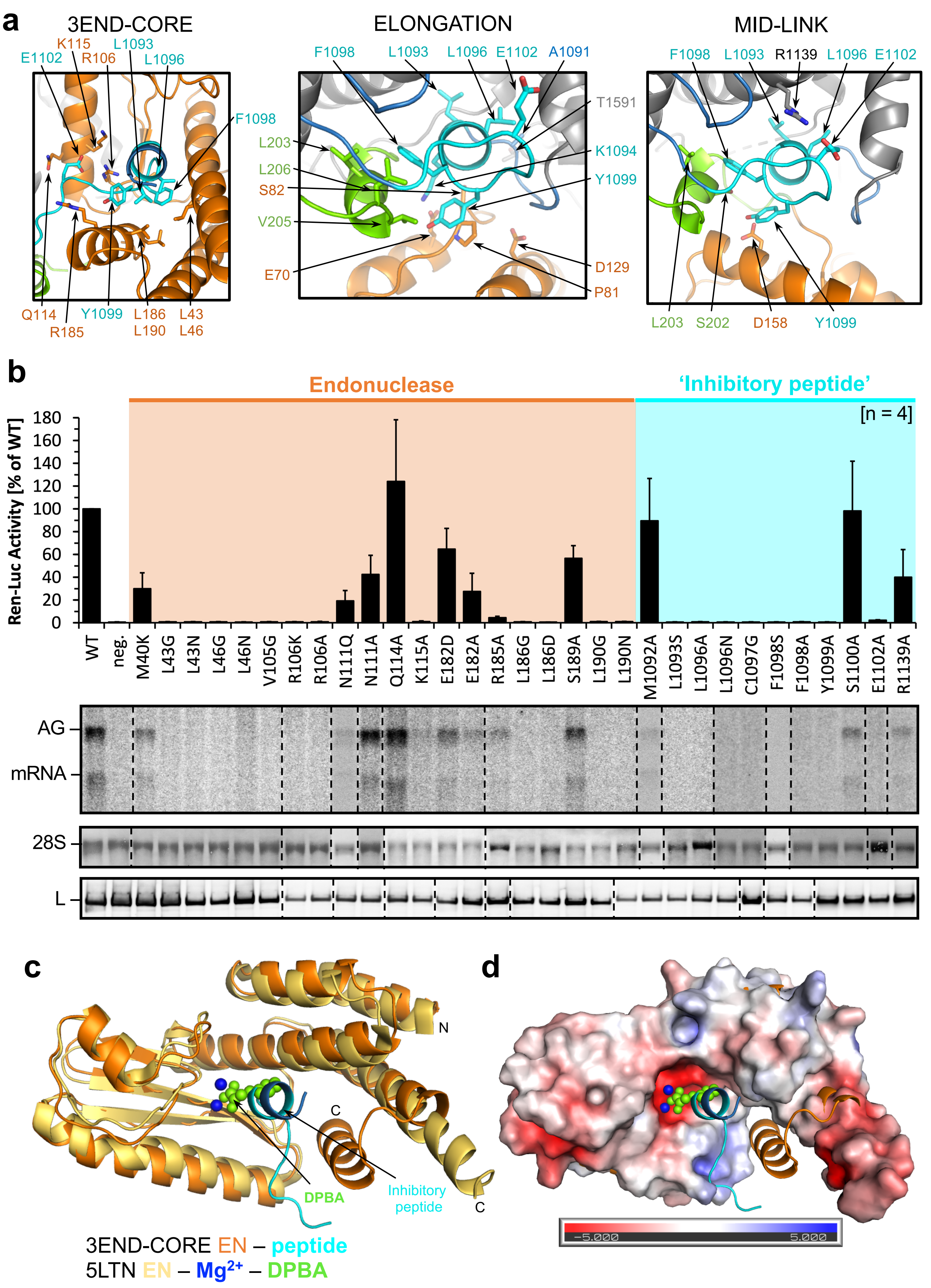

Supplementary Fig. 12

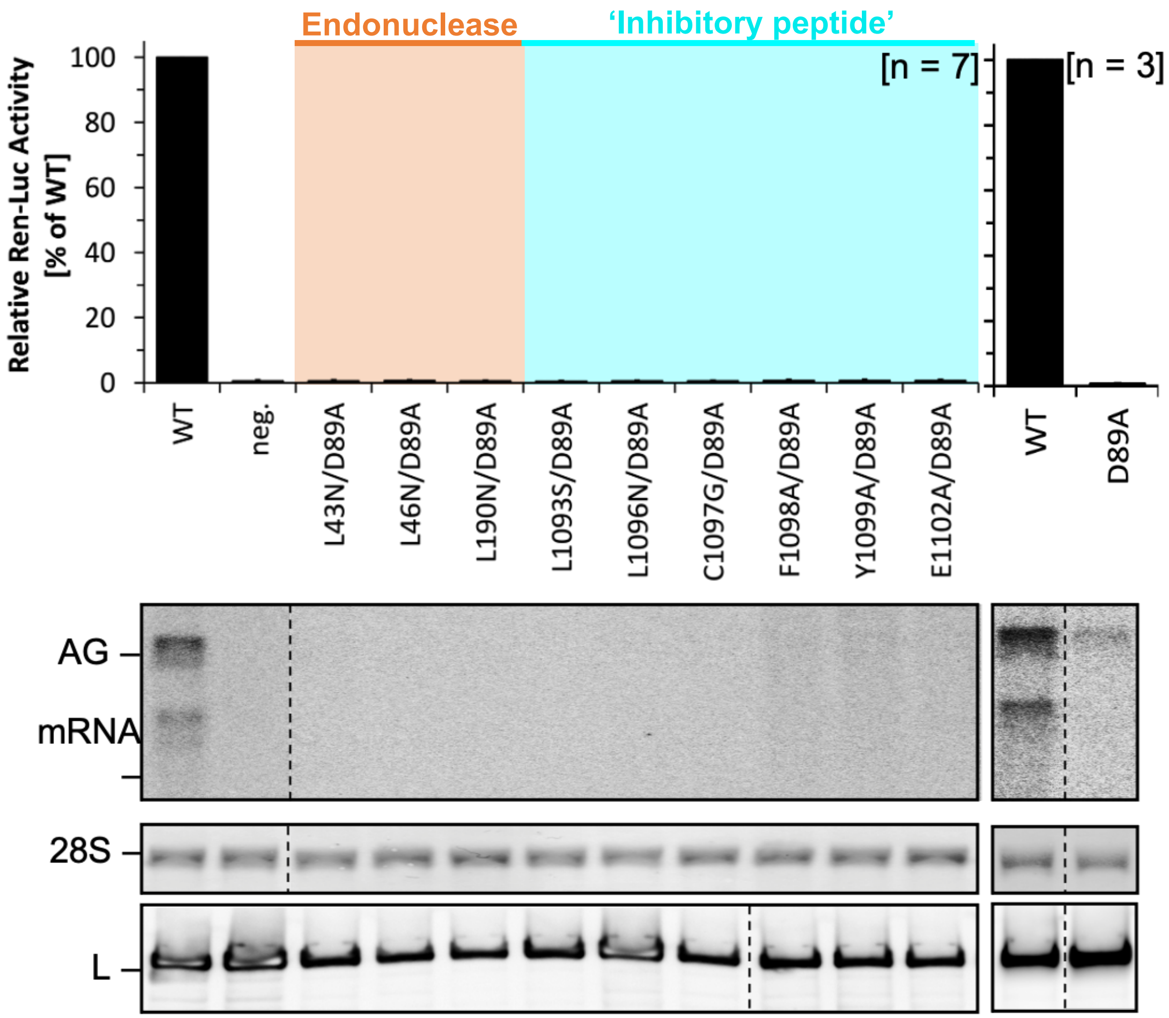

### Supplementary Fig. 13

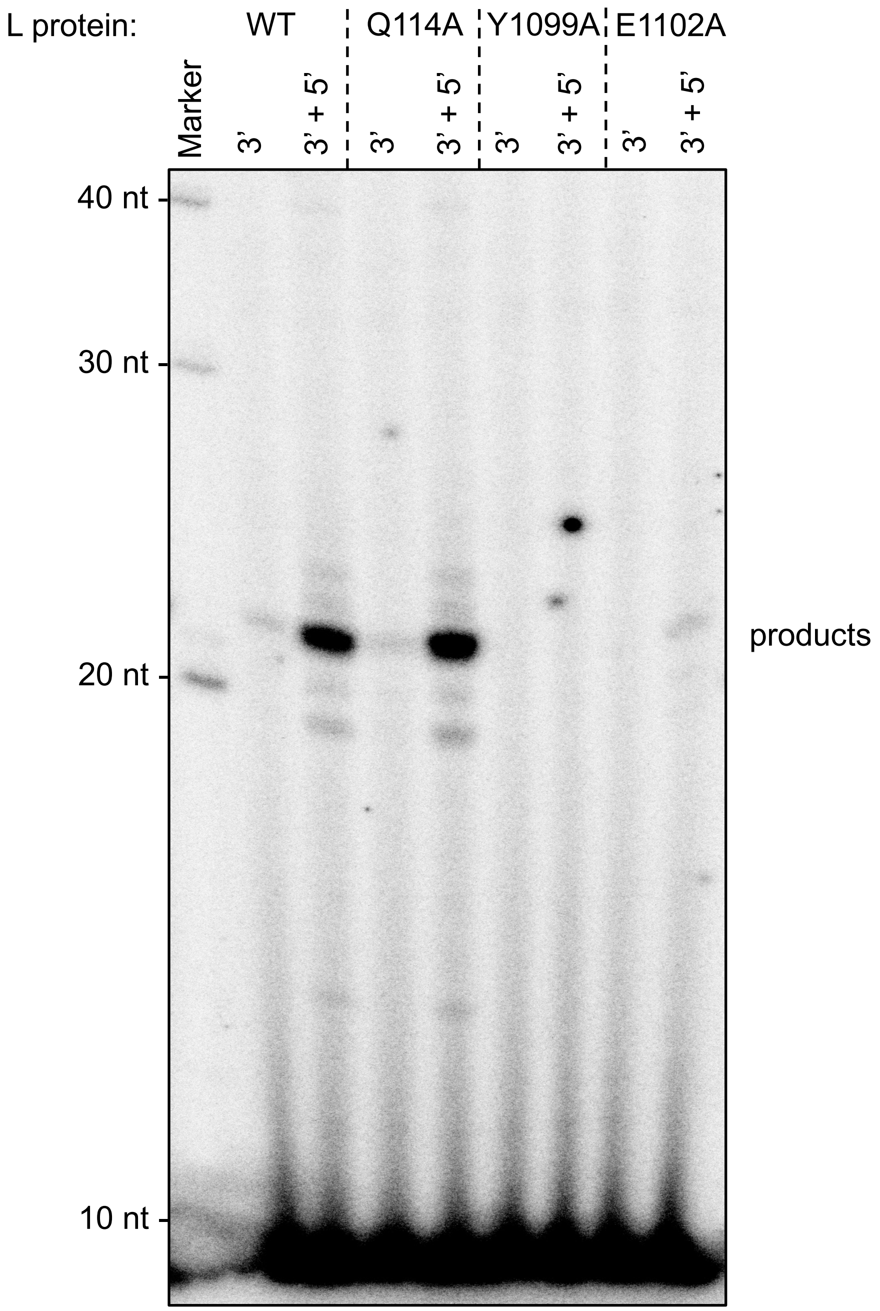

Supplementary Fig. 14

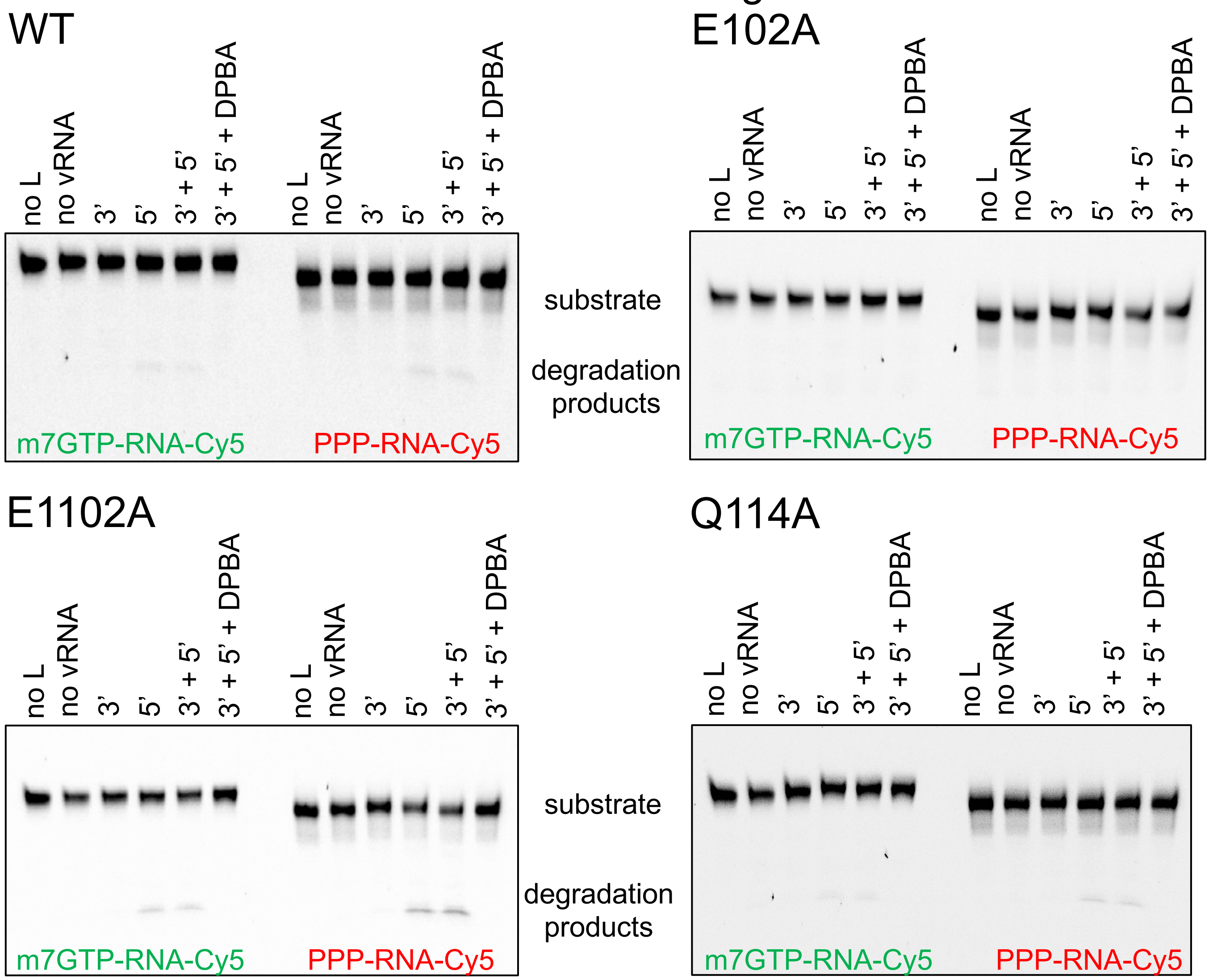

Supplementary Fig. 15

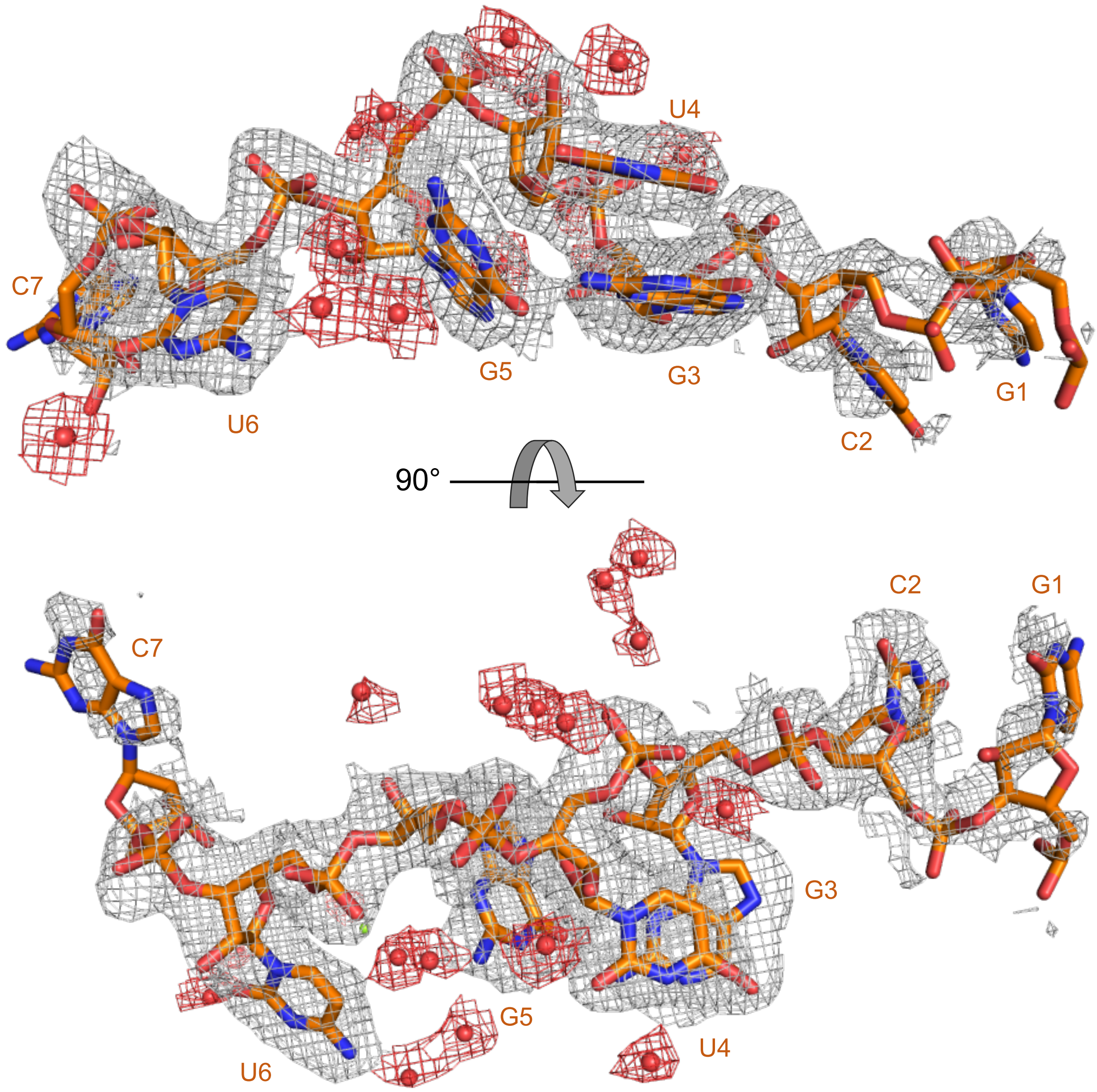

### Supplementary Fig. 16

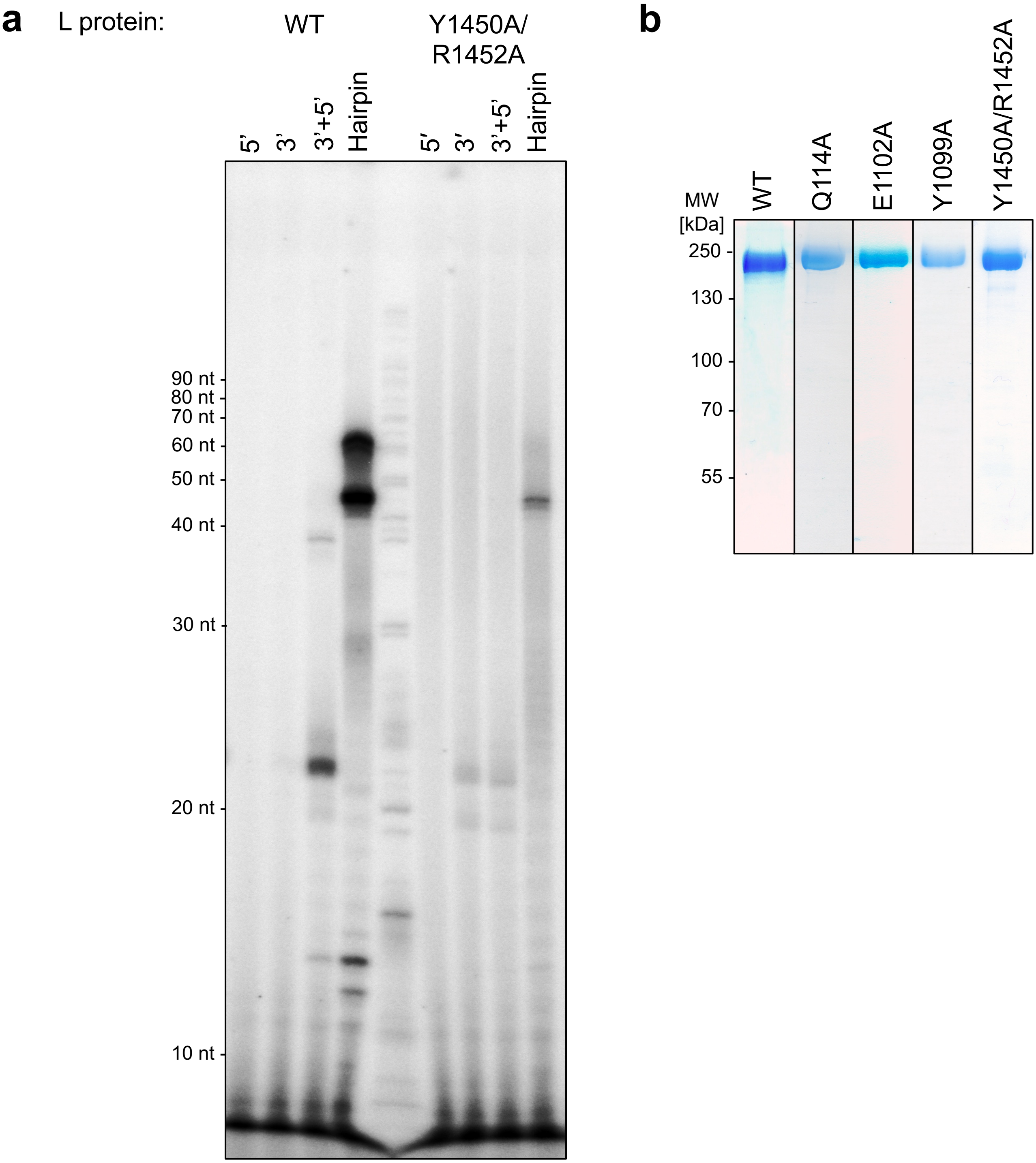

Supplementary Fig. 17

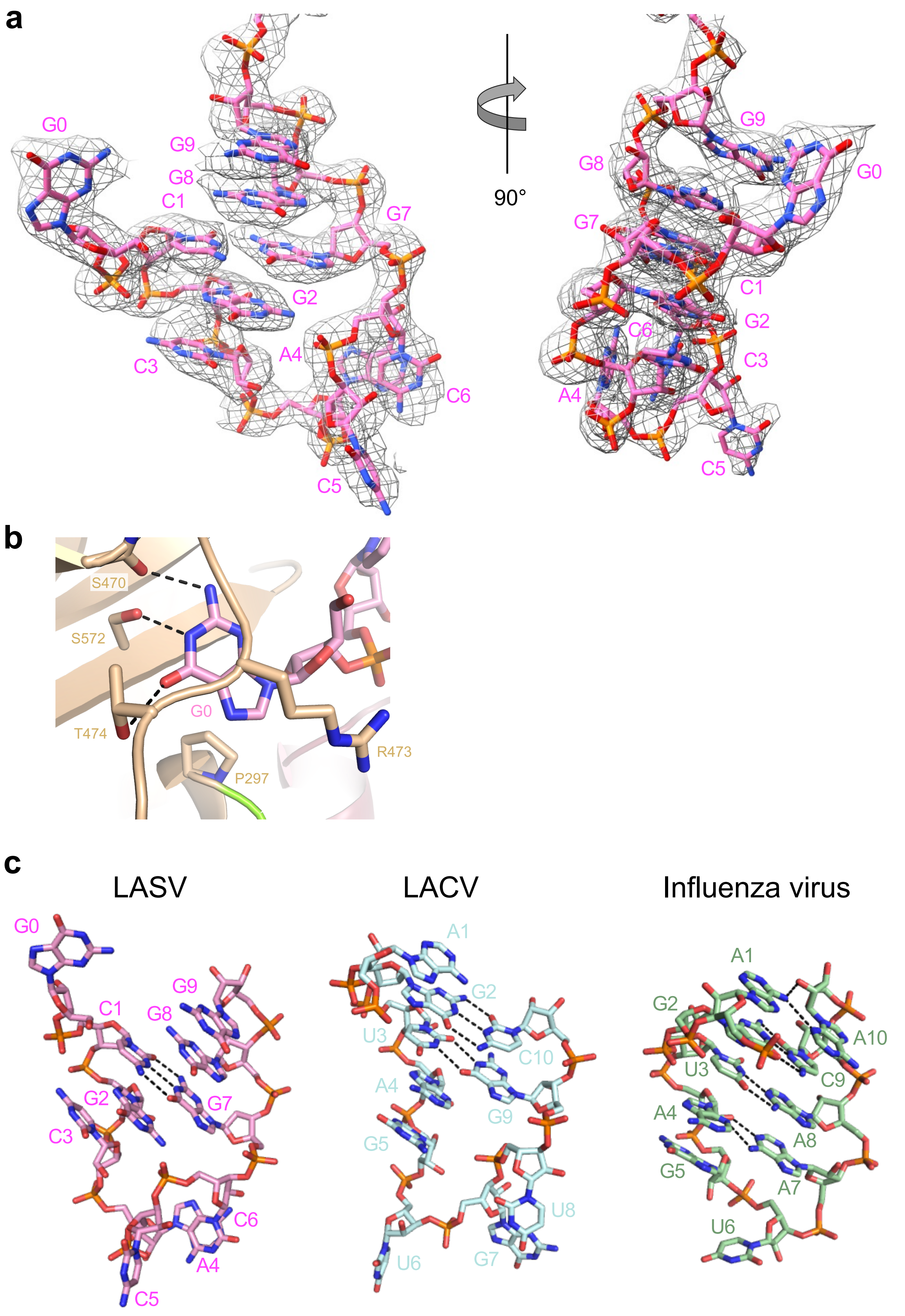

### Supplementary Fig. 18

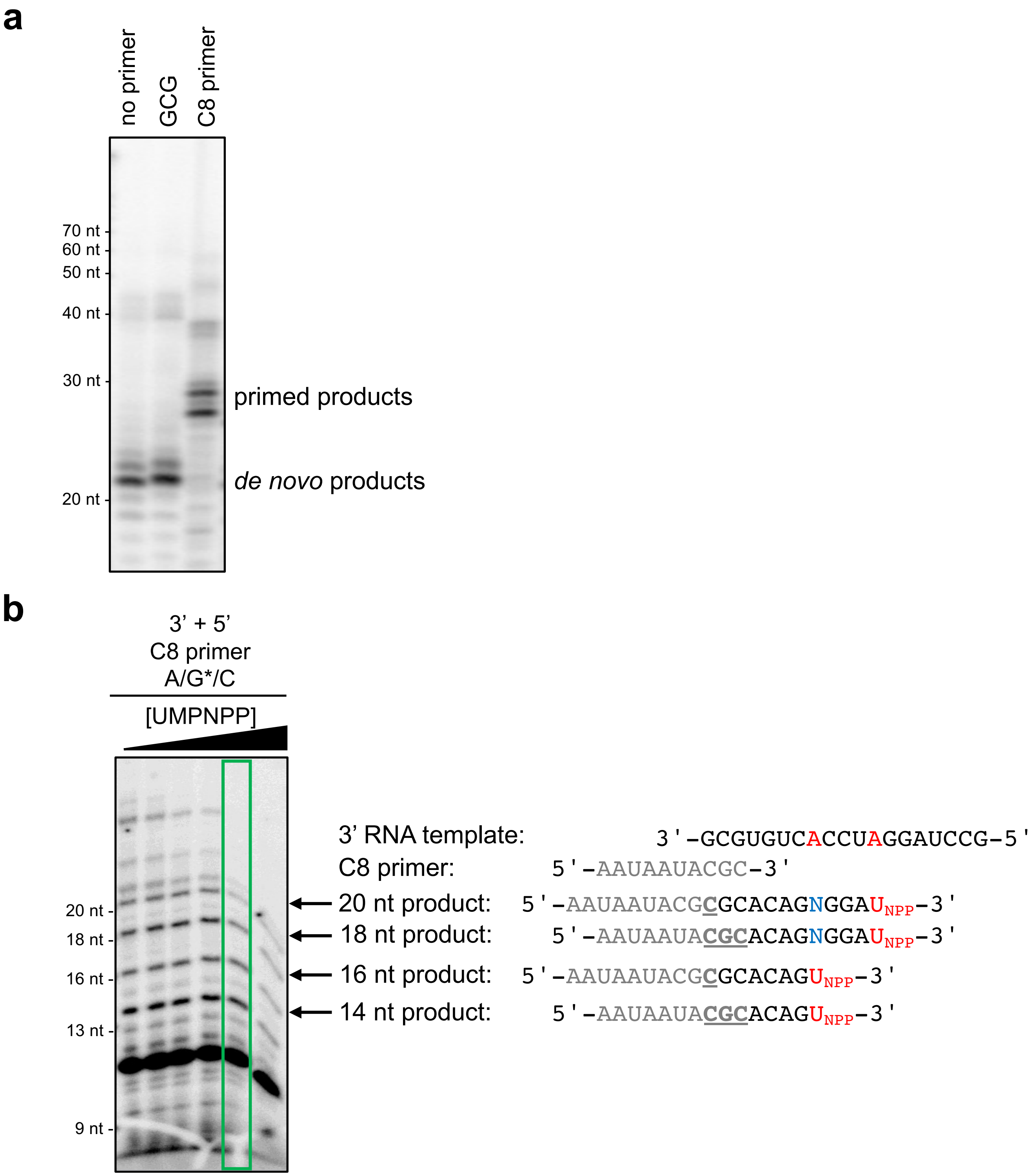

Supplementary Fig. 19

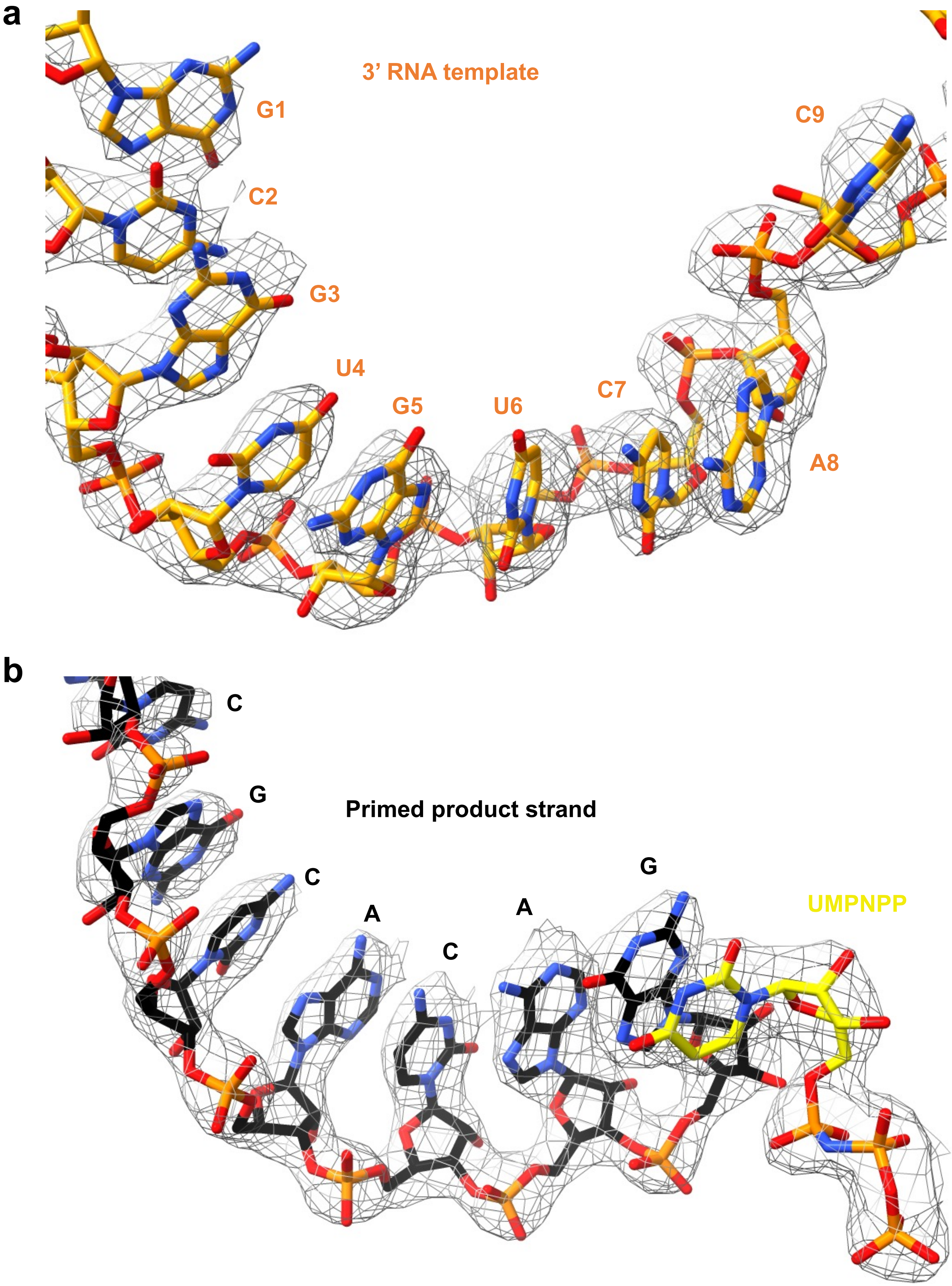

### Supplementary Fig. 20

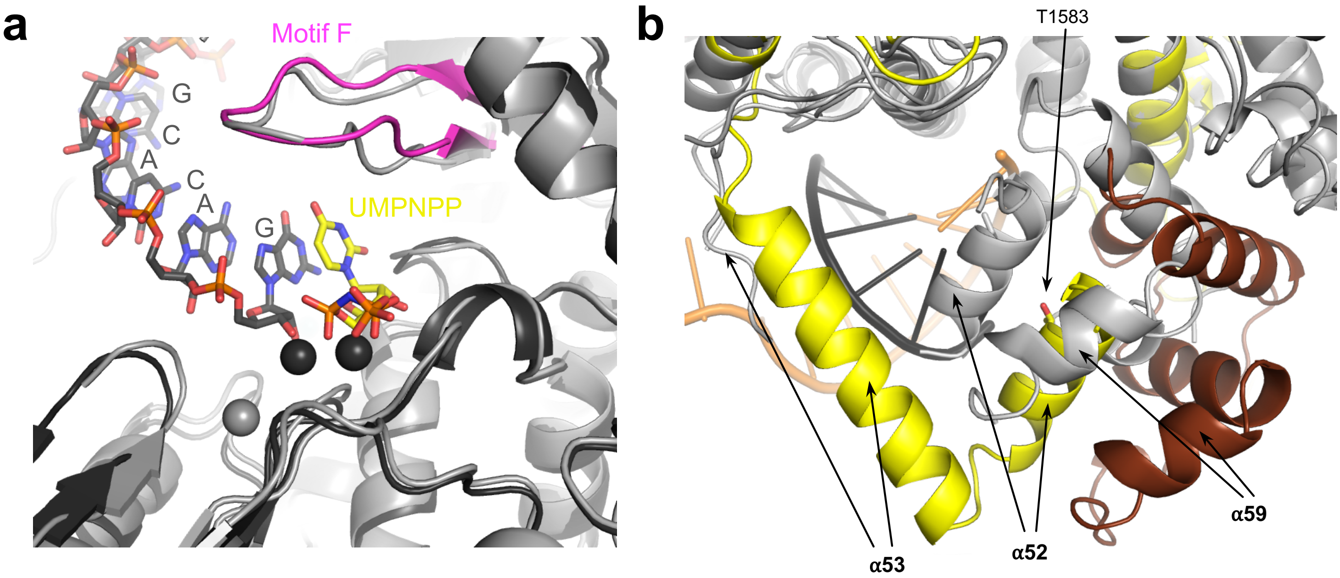

Supplementary Fig. 21

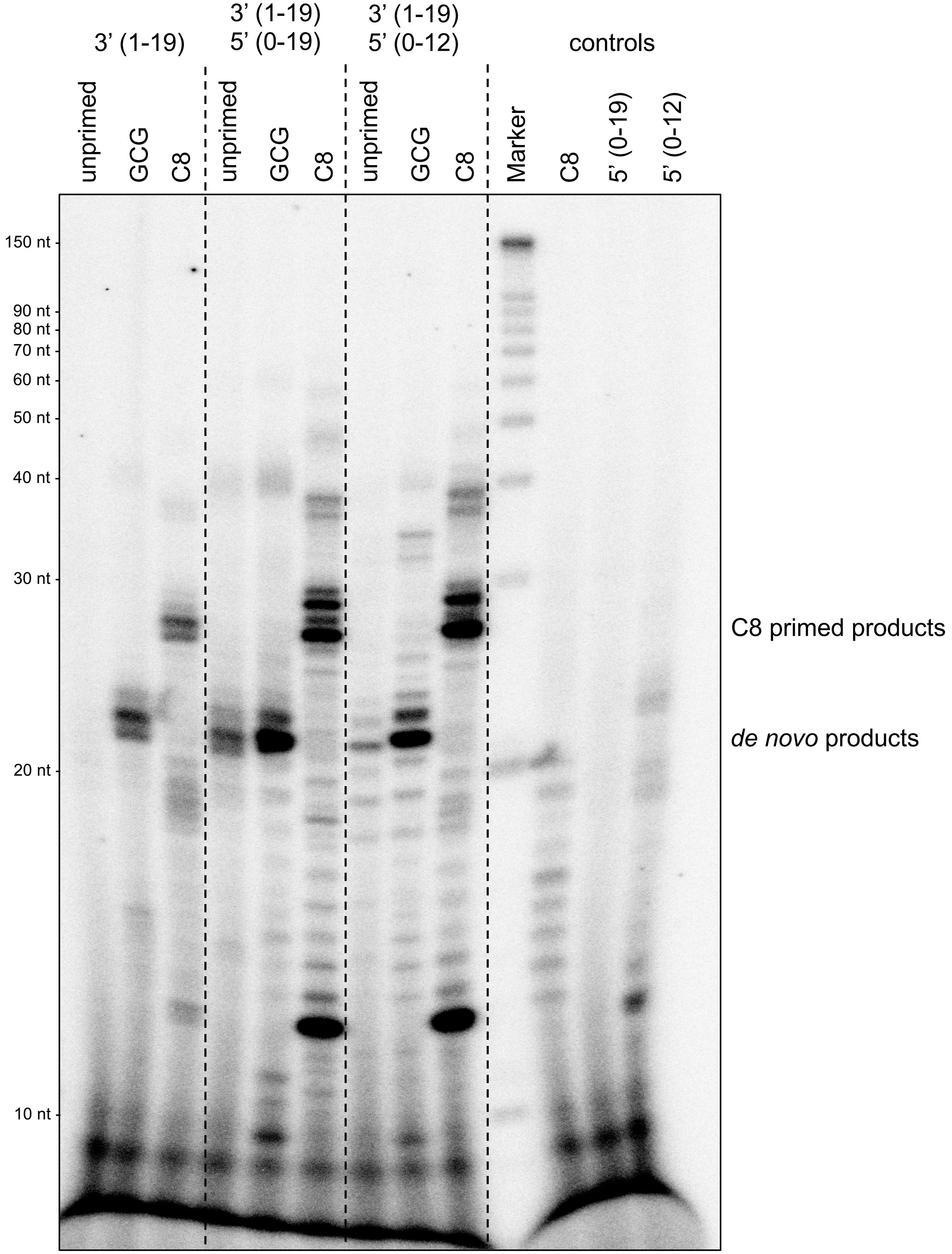

### Supplementary Fig. 22

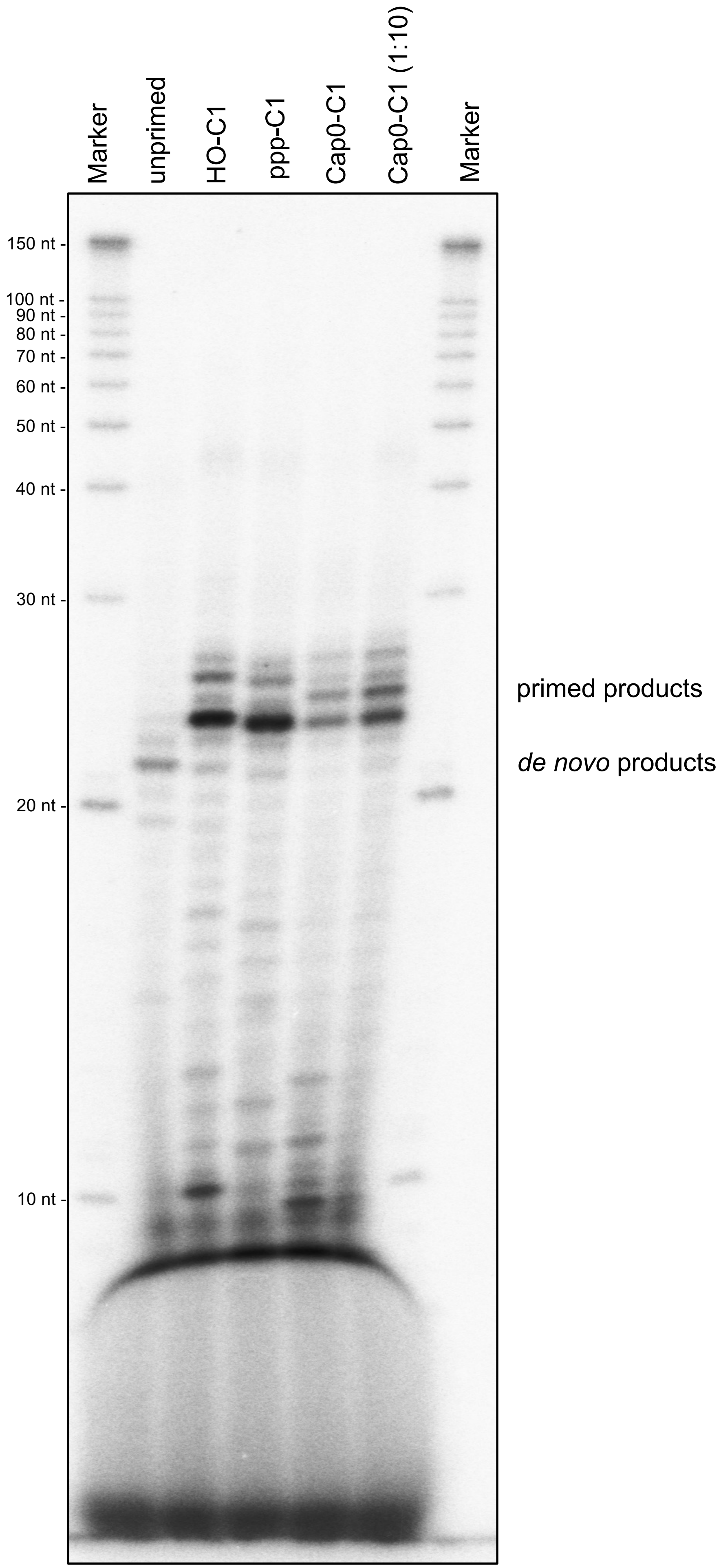

### Supplementary Fig. 23

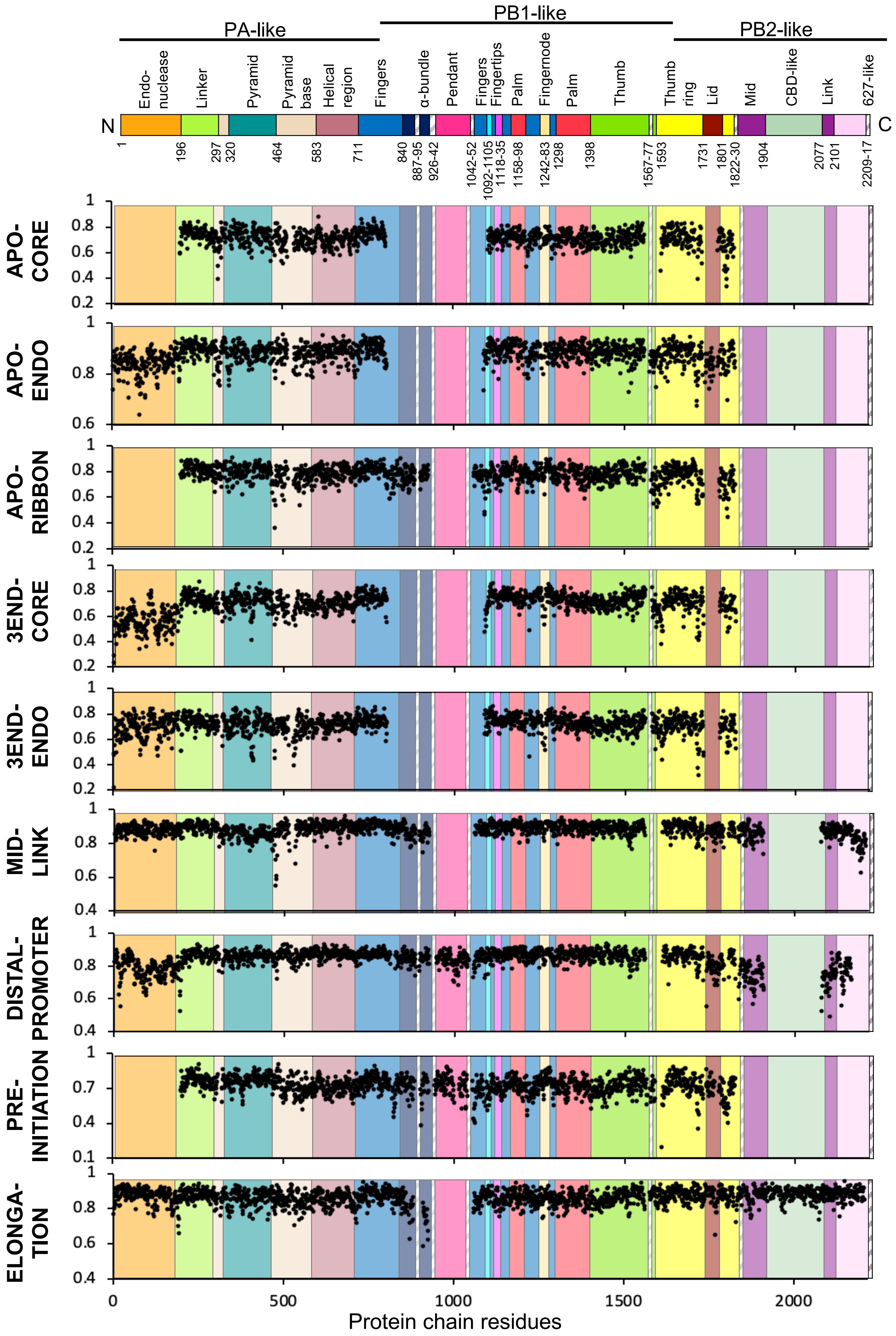
