## Supplementary Table 1 for "Conformational changes in Lassa virus L protein associated with promoter binding and RNA synthesis activity"

**Supplementary Table 1. Cryo-EM data collection, refinement and validation statistics.**

This table provides the statistics for the data collection, refinement and structure validation of the cryo-EM structures. Refinement statistics were generated using Phenix comprehensive validation software.

|  | **APO-CORE** | **APO-ENDO** | **APO-RIBBON** | **3’END-CORE** | **3’END-ENDO** | **PRE-INITIATION** | **DISTAL-PROMOTER** | **MID-LINK** | **ELONGATION** |
| --- | --- | --- | --- | --- | --- | --- | --- | --- | --- |
| PDB | **7OCH** | **7OE3** | **7OE7** | **7OEA** | **7OEB** | **7OJL** | **7OJJ** | **7OJK** | **7OJN** |
| EMDB | **EMD-12807** | **EMD-12860** | **EMD-12861** | **EMD-12862** | **EMD-12863** | **EMD-12955** | **EMD-12953** | **EMD-12954** | **EMD-12956** |
| **Data collection and processing** | CSSB DATA 1 | | | ESRF DATA | | CSSB DATA 3 | CSSB DATA 2 | | CSSB DATA 4 |
| Microscope | CSSB FEI Titan Krios | | | ESRF FEI Titan Krios | | CSSB FEI Titan Krios | CSSB FEI Titan Krios | | CSSB FEI Titan Krios |
| Voltage (kV) | 300 | | | 300 | | 300 | 300 | | 300 |
| Camera | Gatan K3 BioQuantum | | | Gatan K2 Quantum | | Gatan K3 BioQuantum | Gatan K3 BioQuantum | | Gatan K3 BioQuantum |
| Magnification | 105,000 | | | 165,000 | | 105,000 | 105,000 | | 105,000 |
| Nominal negative defocus range (μm) | 0.4 - 3.1 | | | 0.3 - 2.8 | | 0.5 - 2.0 | 0.4 - 3.1 | | 0.6 - 2.6 |
| Exposure time (s) | 2.5 | | | 5.06 | | 2.5 | 2.5 | | 2.5 |
| Electron exposure (e–/Å^2^) | 49.5 | | | 46 | | 53 | 49.5 | | 51.7 |
| Number of frames collected (no.) | 50 | | | 46 | | 50 | 50 | | 50 |
| Number of frames processed (no.) | 23 | | | 26 | | 50 | 25 | | 50 |
| Pixel size (Å) | 0.87 | | | 0.827 | | 0.85 | 0.87 | | 0.85 |
| Micrographs (no.) | 15,448 | | | 6,616 | | 13,204 | 13,462 | | 10,368 |
| Total particle images (no.) | 2,651,000 | | | 1,468,000 | | 2,469,716 | 2,882,000 | | 2,452,281 |
| **Refinement** |  |  |  |  |  |  |  |  |  |
| Particles per class (no.) | 67.5 k | 74 k | 35.4 k | 84.5 k | 40.2 k | 119.9 k | 23.7 k | 40.7 k | 79.6 k |
| Map resolution (Å), 0.143 FSC | 3.14 | 3.35 | 3.73 | 2.70 | 3.04 | 3.34 | 3.89 | 3.50 | 2.92 |
| Map sharpening *B* factor (Å^2^) | -80.2 | -87.1 | -93.5 | -42.8 | -56.7 | -104.1 | -124.6 | -104.3 | -45.9 |
| Map versus model cross-correlation  CC-MASK | 0.8337 | 0.8783 | 0.8250 | 0.8374 | 0.8114 | 0.8361 | 0.8245 | 0.8804 | 0.8635 |
| **Model composition** |  |  |  |  |  |  |  |  |  |
| Non-hydrogen atoms | 9516 | 11904 | 11104 | 11703 | 11682 | 12730 | 15344 | 14330 | 16984 |
| Protein residues | 1181 | 1478 | 1381 | 1432 | 1434 | 1485 | 1847 | 1785 | 2020 |
| Nucleotide residues | 0 | 0 | 0 | 7 | 6 | 33 | 21 | 0 | 39 |
| Water | 0 | 0 | 0 | 17 | 1 | 0 | 0 | 0 | 0 |
| Ligands | 2 | 2 | 2 | 2 | 2 | 1 | 1 | 2 | 6 + NTP |
| ***Mean B* factors (Å^2^)** |  |  |  |  |  |  |  |  |  |
| Protein | 54.51 | 119.18 | 66.77 | 69.63 | 44.51 | 40.31 | 158.11 | 124.91 | 41.45 |
| Nucleotide | - | - | - | 50.71 | 37.82 | 55.08 | 141.68 | - | 46.47 |
| Ligand | 75.78 | 115.19 | 95.22 | 70.43 | 64.00 | 13.19 | 174.58 | 137.47 | 32.66 |
| Water | - | - | - | 30.48 | 17.56 | - | - | - | - |
| **R.m.s. deviations** |  |  |  |  |  |  |  |  |  |
| Bond lengths (Å) | 0.003 | 0.003 | 0.005 | 0.004 | 0.004 | 0.003 | 0.003 | 0.006 | 0.003 |
| Bond angles (°) | 0.537 | 0.519 | 0.641 | 0.772 | 0.557 | 0.600 | 0.716 | 0.663 | 0.460 |
| **Validation** |  |  |  |  |  |  |  |  |  |
| MolProbity score | 2.05 | 2.22 | 1.95 | 1.66 | 1.71 | 2.06 | 2.33 | 2.08 | 1.54 |
| All-atom clashscore | 3.89 | 5.91 | 6.06 | 6.04 | 7.67 | 11.71 | 13.28 | 9.33 | 3.90 |
| Poor rotamers (%) | 3.8 | 3.39 | 0.08 | 0.08 | 0.53 | 0.15 | 0.0 | 0.12 | 0.98 |
| **Ramachandran plot** |  |  |  |  |  |  |  |  |  |
| Favored (%) | 93.44 | 91.84 | 87.10 | 95.31 | 95.74 | 92.24 | 82.94 | 88.64 | 94.78 |
| Allowed (%) | 6.56 | 8.16 | 12.82 | 4.47 | 4.04 | 7.62 | 16.73 | 11.36 | 5.17 |
| Outliers (%) | 0 | 0 | 0.07 | 0.21 | 0.21 | 0.14 | 0.33 | 0 | 0.05 |
