## Supplementary Table 2 for "Conformational changes in Lassa virus L protein associated with promoter binding and RNA synthesis activity"

**Supplementary Table 2: Synthetic RNA oligos used in the assays.** The table lists the RNA oligonucleotides that were used in the described assays. The specific sequence, length in nucleotides and the identifier used to label the RNA in the experimental descriptions are given. Underlined nucleotides in the 5' (nts 0-19) and 3' (nts 1-19) indicate mismatches between the two otherwise complementary RNA strands. All RNAs were synthesized by Biomers, TriLink BioTechnologies or Chemgenes.

|  | **Length [nt]** | **Identifier** |  | **Sequence** |  |
| --- | --- | --- | --- | --- | --- |
| vRNA | 20 | 5' (0-19) | 5' HO- | GCGCACCGGGGAUCCUAGGC | -OH 3' |
|  | 20 | 5' (0-19) Pi | 5' HO- | GCGCACCGGGGAUCCUAGGC | -PO_4_ 3' |
|  | 10 | 5' (10-19) | 5' HO- | GAUCCUAGGC | -OH 3' |
|  | 19 | 3' (1-19) | 3' HO- | GCGUGUCACCUAGGAUCCG | -OH 5' |
|  | 19 | 3' (1-19) Pi | 3' PO_4_- | GCGUGUCACCUAGGAUCCG | -OH 5' |
|  | 16 | 3' (1-16) | 3' HO- | GCGUGUCACCUAGGAU |  |
| EMSA | 10 | 3' (1-10) | 3' HO- | GCGUGUCACC | -Cy3 5' |
| EN | 17 | PPP-RNA | 5' ppp- | AAACGCAACAACAACAC | -Cy5 3' |
|  | 18 | Cap-RNA | 5' Cap^0^- | AAACGCAAGCAACAACAC | -Cy5 3' |
| Primer | 6 | St1 | 5' HO- | GCGCAC | -OH 5' |
|  | 6 | C1 | 5' HO- | AAACGC | -OH 3' |
|  | 6 | C1ppp | 5' ppp- | AAACGC | -OH 3' |
|  | 6 | C1cap | 5' Cap^0^- | AAACGC | -OH 3' |
|  | 10 | C8 | 5' HO- | AAUAAUACGC | -OH 5' |
